# Transcriptome landscape of the developing olive fruit fly embryo delineated by Oxford Nanopore long-read RNA-Seq

**DOI:** 10.1101/478172

**Authors:** Anthony Bayega, Spyros Oikonomopoulos, Eleftherios Zorbas, Yu Chang Wang, Maria-Eleni Gregoriou, Konstantina T Tsoumani, Kostas D Mathiopoulos, Jiannis Ragoussis

## Abstract

The olive fruit fly or olive fly (*Bactrocera oleae*) is the most important pest of cultivated olive trees. Like all insects the olive fly undergoes complete metamorphosis. However, the transcription dynamics that occur during early embryonic development have not been explored, while detailed transcriptomic analysis in the absence of a fully annotated genome is challenging. We collected olive fly embryos at hourly intervals for the first 6 hours of development and performed full-length cDNA-Seq using a purpose designed SMARTer cDNA synthesis protocol followed by sequencing on the MinION (Oxford Nanopore Technologies). We generated 31 million total reads across the timepoints (median yield 4.2 million per timepoint). The reads showed 98 % alignment rate to the olive fly genome and 91 % alignment rate to the NBCI predicted *B. oleae* gene models. Over 50 % of the expressed genes had at least one read covering its entire length validating our full-length RNA-Seq procedure. Expression of 68 % of the predicted *B. oleae* genes was detected in the first six hours of development. We generated a *de novo* transcriptome assembly of the olive fly and identified 3553 novel genes and a total of 79,810 transcripts; a fourfold increase in transcriptome diversity compared to the NCBI predicted transcriptome. On a global scale, the first six hours of embryo development were characterized by dramatic transcriptome changes with the total number of transcripts per embryo dropping to half from the first hour to the second hour of embryo development. Clustering of genes based on temporal co-expression followed by gene-set enrichment analysiss of genes expressed in the first six hours of embryo development showed that genes involved in transcription and translation, macro-molecule biosynthesis, and neurodevelopment were highly enriched. These data provide the first insight into the transcriptome landscape of the developing olive fly embryo. The data also reveal transcript signatures of sex development. Overall, full-length sequencing of the cDNA molecules permitted a detailed characterization of the isoform complexity and the transcriptional dynamics of the first embryonic stages of the *B. oleae*.

## Introduction

The olive fruit fly or olive fly (*Bactrocera oleae*), is an insect of huge economic importance in the olive agribusiness industry costing an estimated 800 million US dollars annually[1]. Adult female flies lay eggs in the olive fruits where the eggs hatch and develop into larvae. The larvae feed on the olive sap causing serious qualitative and quantitative damages to the fruits, making olive flies the most important pest for both wild and cultivated olive fruits.

Olive flies are Dipteran insects belonging to the Tephritidae family. Although olive flies are holometabolous transcriptional events that occur during early embryo development have not been studied. Delineating the transcriptional dynamics in the first few hours of embryo development is important to determine the mechanisms of tissue development and the genes required in the early life of the embryo. These transcriptional dynamics can also be used to determine the mechanism of sexual commitment in the olive fly, a process known to be initiated within the first 6 hours of embryo development[2] and indeed, a fundamental question that has been at the center of olive fly research for decades[2].

Gene expression is dynamic and tightly regulated. In rapidly changing systems like developing embryos precise and coordinated dramatic shifts in transcriptome kinetics occur in quick succession. Although relative normalization is a common method of measuring and comparing transcriptional changes, direct absolute normalization has been shown to perform superiorly to relative normalization in capturing transcriptional dynamics[3]. Coupled with time-course experimentation, absolute normalization enables direct measurement of transcript kinetics thus providing a quantitative understanding of the rate of change of transcript copy numbers with time. Such a quantitative understanding of the transcriptome during embryogenesis can greatly improve our understanding of the impact of different transcript regulation strategies on gene expression.

cDNA sequencing using short-read technologies such as sequence-by-synthesis (Illumina Inc., USA) is currently the standard in high throughput transcriptional exploration. Due to their short-read nature however, these technologies are inefficient in delineating the full depth of transcriptome landscape particularly regarding isoform diversity[4, 5]. We and others have shown that long-read technologies provide full-length transcript resolution[6, 7] and enable identification of hitherto unknown genes and isoforms [8-11]. This can be particularly useful for organisms that are not well characterized at a genomics level. Among long-read technologies, the Oxford Nanopore Technologies (ONT, UK) protein nanopore sequencing is particularly attractive due to higher throughput, simpler library preparation workflow, and gene expression quantification that is comparable to current standards [8-11].

Here we used the ONT sequencing technology and the MinION sequencer to explore the transcriptome of olive fly embryos collected at hourly intervals for the first 6 hours post oviposition (hpo). We refine the *B. olea*e current NCBI predicted gene models by providing a rich set of isoform models found in the first six hours of embryonic development and identify mis-annotations in the currently available NCBI gene model annotations. We also measured transcript kinetics, which can be linked to biological processes occurring during embryogenesis. We provide a set of co-regulatory genes across the different time points that can help to understand the sequence of events during early development. We also explored a range of data analysis tools currently available for long-read technologies.

## Results

### The current *B. oleae* genome assembly and genome annotation

The olive fly has six pairs of chromosomes which include a pair of heterochromatic sex chromosomes with the male being the heterogametic sex[12]. The *B. oleae* genome size was initially estimated by qPCR to be in the range of 322 Mb[13]. We previously submitted to NCBI a *B. oleae* genome assembled from short-reads and long-read scaffolding. This assembly (GeneBank accession GCA_001188975.2) has a total size of 471,780,370 bases and is contained in 36,198 scaffolds, with a contig N50 of 135,231 bases. This genome was annotated using the NCBI Eukaryotic Genome Annotation Pipeline yielding the Bactrocera oleae Annotation Release 100. This annotation contains a total of 13936 genes and pseudogenes of which 13198, 392, 346 are predicted to be protein-coding, non-coding, and pseudogenes, respectively. Further, 2,759 genes are predicted to have variants (isoforms). In total, the *B. oleae* genome was predicted to contain 19,694 transcripts of which 18702, 411, 393, 188 are mRNA (with CDSs), tRNA, lncRNAs, and miscellaneous RNAs, respectively. Whereas the mean length of the genes and transcripts is 9,597 bp and 2,259 bp, respectively, the longest gene is 497,921 bp while the longest transcript is 59,475 bp. All alignments and genes reported in this article refer to these references: genome GCA_001188975.2 and annotation NCBI Bactrocera oleae Annotation Release 100, unless otherwise stated.

Because only 14,555 mRNA (with CDSs) out of the 18,702 in the NCBI annotation (77.8%) were assigned a gene product corresponding to their respective *Drosophila melanogaster* homolog, we obtained the NCBI predicted proteins and re-determined the Uniprot homologues to *D. melanogaster* (see Materials and Methods). Of the 13198 protein coding genes 12494 (95%) were assigned a *D. melanogaster* homologue (E-value ≤ 1e-3). Of these, 57% were identified in the UniProtKB/Swiss-prot database, which comprises high quality manually annotated and nonredundant proteins while the remaining 43% were identified in the UniProtKB/TrEMBL database which contains high quality computationally annotated and classified proteins.

### *De novo* genome-guided transcriptome assembly of the olive fly identifies new genes and isoforms

We performed cDNA synthesis for mixed sex *B. oleae* embryos collected at one-hour intervals for the first six hours of development using an optimized and customized SMARTer protocol[14] aimed at capturing poly(A)+ RNA(Figure 1**A**, See Supplementary protocol, Supplementary Table 1). cDNA sequencing libraries were generated following the SQK-LSK108 protocol (ONT) and sequenced on the MinION using 1D R9.4 flow cells (see Supplementary Table 2 and 3 for sequencing statistics). We also included 2 cDNA libraries separately generated from the heads of adult male and female flies.

**Figure 1:**
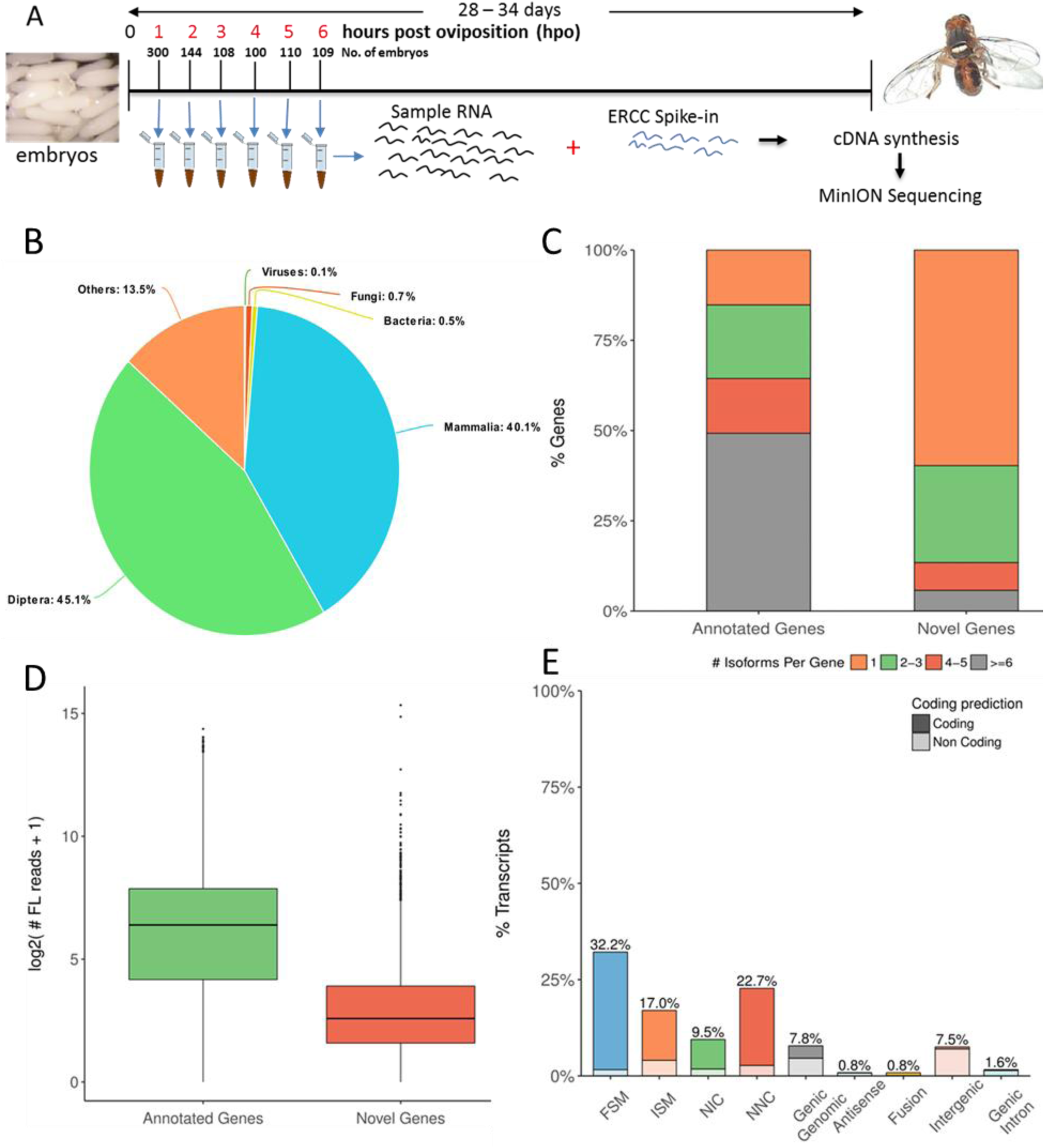
Exploration of the transcriptome assembly. **A)** Schematic of cDNA library generation and sequencing. Embryos were collected at hourly point post oviposition (hpo), counted and total RNA extracted using the Trizol method. At cDNA synthesis step, external RNA standards (ERCC) were added to each sample commensurate to the number of embryos that were used. The Smart-Seq2 protocol was used to generate full length cDNA, followed by PCR amplification of the cDNA. The Oxford Nanopore Technologies (ONT) SQK-LSK108 protocol for library preparation was then followed, albeit with some custom changes. The library was then sequenced on the ONT MinION, followed by basecalling using ONT Albacore basecaller. **B**) Distribution of top blastp hits to Uniprot-Swiss prot database. C) Distribution of isoform among NCBI annotated genes and Novel genes D) Gene expression levels of annotated and novel genes using long-read counts. C) Transcript length distribution by structural classification. E) Distribution of transcripts among structural categories (SQANTI[18]). FSM; full splice match, ISM; incomplete splice match, NIC; novel in catalogue, NNC; novel not in catalogue (see Supplementary Table 6 for explanation).

We developed a computational pipeline (Supplementary Figure 1) to perform *de novo* transcriptome assembly. We generated 31 million reads of which 22 million reads (71 %) were error-corrected using Canu[15]. We then focused only on full-length reads identified as those that possessed both 5’ adapters and both poly(A) and 3’ adapter (Supplementary Figure 1). These were then pre-processed to orient the strands, trim adapters, and correct errors using short-reads. First, we used 3 million reads generated from the 5-hour timepoint to compare three genome-guided *de novo* transcriptome assembly tools; TAMA (https://github.com/GenomeRIK/tama), Cupcake ToFU (https://github.com/Magdoll/cDNA_Cupcake/wiki/Cupcake-ToFU%3A-supporting-scripts-for-Iso-Seq-after-clustering-step), and TAPIS[16]. We opted for Cupckae ToFU due to its computational efficiency, high sensitivity and precision (see Supplementary Materials for details).

Of the 31 million reads, 14.7 million reads passed pre-processing and were aligned to the *B. oleae* genome using GMAP[17]. Because the ToFU software of our assembly workflow was set to consider reads that were aligned at least 99 % in length and with at least 95 % identity, out of the total 14.7 million reads, 3.9 million were used to derive the transcriptome assembly. The *de novo* assembled transcriptome was analyzed using SQANTI[18] and PRAPI[19]. The ToFU transcriptome assembly contained a total of 11,883 genes and 79,810 isoforms (Table 1) of which 8330 genes matched the NCBI annotated genes while 3553 genes were novel. All these correspond to a four-fold expansion of the olive fly transcriptome at the isoform level over the current NCBI annotation. A blast search of the predicted protein sequences against Uniprot Swiss-prot database showed the top hits were of the order Diptera (45.1%, Figure 1B), followed by class Mammalia (40.1%). Further analsysis, revealed that 99% of the hits to Mammalia also had hits to Diptera. Hits to Viruses, Fungi, and Bacteria accounted for only 1.3%.

**Table 1:**
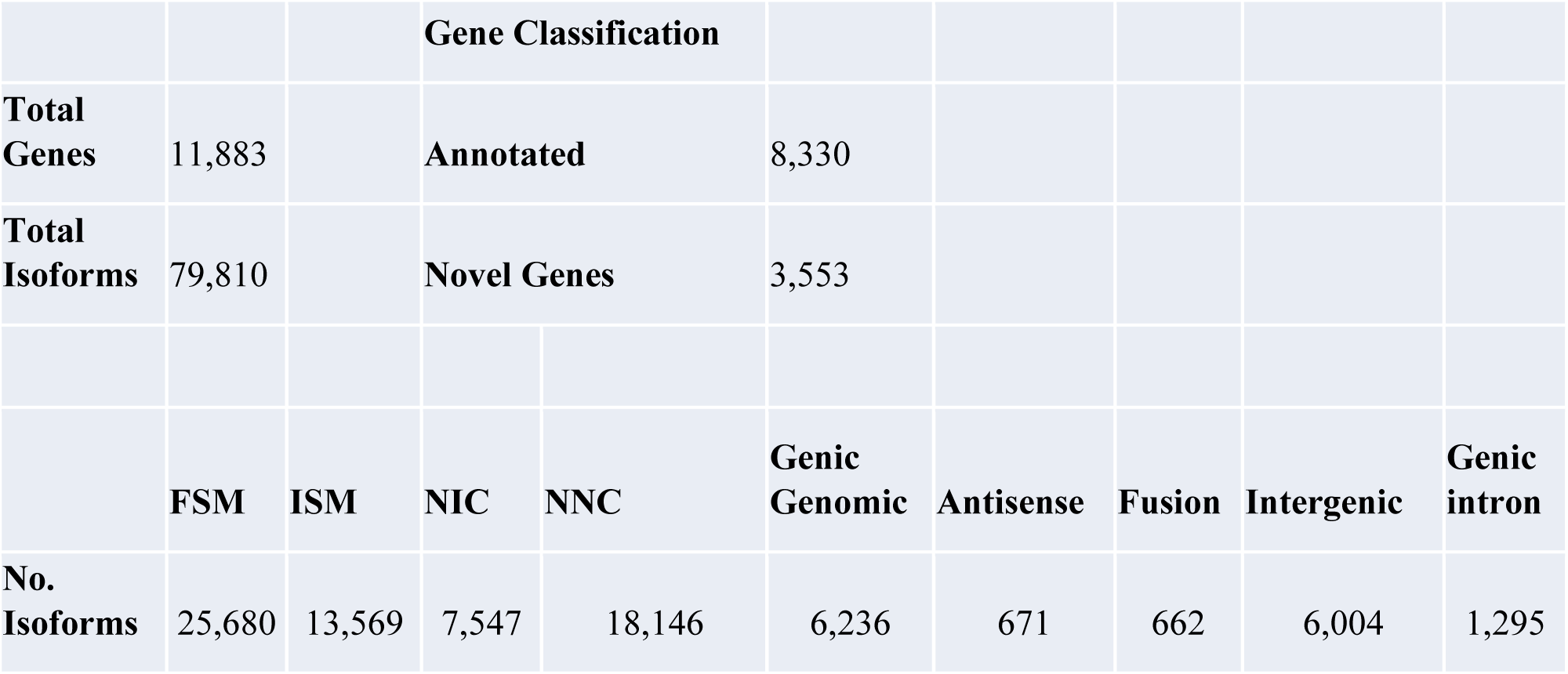
Summary of the long-read genome guided *de novo* transcriptome assembly generated. The assembly was generated from full-length Canu[15] consensus corrected and Lordec[44] short-read corrected reads using Cupcake ToFU* tool. The full-length reads used included embryo and adult male and female heads. The transcriptome assembly was analyzed using SQANTI[18]. Isoform categories FSM; full splice match, ISM; incomplete splice match, NIC; novel in catalogue, NNC; novel not in catalogue (see Supplementary Table 6 for explanation). *(https://github.com/Magdoll/cDNA_Cupcake/wiki/Cupcake-ToFU%3A-supporting-scripts-for-Iso-Seq-after-clustering-step)

|  |  | Gene Classification |  |  |  |  |  |  |  |
| --- | --- | --- | --- | --- | --- | --- | --- | --- | --- |
| Total Genes | 11,883 | Annotated |  |  | 8,330 |  |  |  |  |
| Total Isoforms | 79,810 | Novel Genes |  |  | 3,553 |  |  |  |  |
|  | FSM | ISM | NIC | NNC | Genic Genomic | Antisense | Fusion | Intergenic | Genic intron |
| No. Isoforms | 25,680 | 13,569 | 7,547 | 18,146 | 6,236 | 671 | 662 | 6,004 | 1,295 |

Surprisingly, although 3553 novel genes were identified, only 269 were predicted to contain open reading frames. Over 50% of novel genes were mono-exon compared to annotated genes where over 80% of the genes were multi-exon and also contained much higher percentage of mono-isoform genes (>50% compared to 86% among annotated genes, **Figure 1B**). Novel genes also showed lower expression compared to annotated genes (**Figure 1**D). Structurally, SQANTI categorizes the transcripts into 9 groups depending on their splice junction and genomic coordinate. These include; full splice match (FSM), incomplete splice match (ISM), novel in catalogue (NIC), novel not in catalogue (NNC), genic genomic, antisense, fusion, intergenic, genic intron. Most of the ToFU collapsed transcripts were a FSM (32.2%), followed by transcripts containing alternatively spliced junctions (22.7%, Table 1, **Figure 1E**). A more detailed exploration of the transcriptome is shown in Extended material.

Using PRAPI, we identified 63 genes that were miss-annotated as two or more separate genes, but we could find single transcript reads covering the two genes (see Supplementary Figure 2).

### Direct Absolute Normalization of RNA-Seq Data outperforms relative normalization

The method and justification for absolute normalization have been previously reported by Owens et al.[3]. To obtain absolute transcript numbers we added ERCC internal spike-in RNA standards in our cDNA synthesis step at a constant ratio per number of embryos used in each timepoint. Following sequencing and alignment, absolute normalization was achieved following 2 steps; 1) relative normalization for sequencing depth using Mandalorion[8], resulting in transcript abundances in number of reads per gene per 10000 mapped reads (RP10K). We noted that the relative normalized abundances of our RNA standards varied with time most likely due to changes in the amount of poly(A) RNA in the embryo (**Figure 2A**). In the second step, we used the standard curve generated from ERCC standards to transform our relative counts to absolute counts per embryo. Here we noted that the ERCC standards did not vary significantly across timepoints (**Figure 2B**, Supplementary Figure 3).

**Figure 2:**
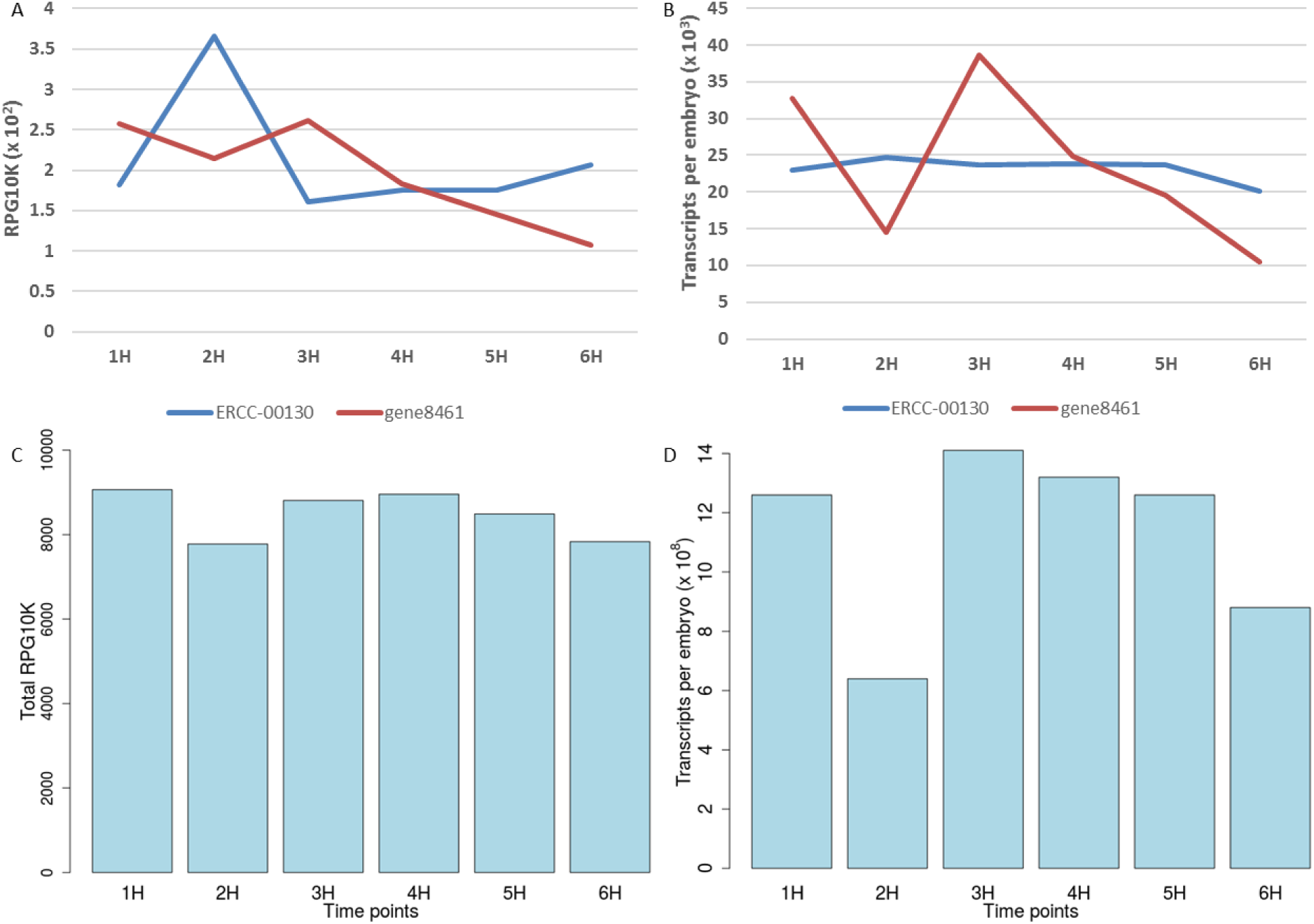
Comparison of relative and absolute normalization. A) Relative gene expression of ERCC 00130 (blue) and gene8461 (red) as obtained from Mandalorion software[8]. For each gene Mandalorion reports its quantification as reads per gene per 10000 mapped reads (RPG10K). B) Same as ‘A’ but showing absolute gene expression levels per embryo. C) Summed RPG10K for each timepoint D) Same as C but showing total number of transcripts per embryo across the six timepoints following absolute normalization.

Interestingly, in contrast to the relative expression, when the absolute number of transcript per embryo for all genes were summed and plotted across timepoints (**Figure 2** C and D), the profile mirrored that of cDNA generated per embryo (Supplementary Figure 4A), thus validating the absolute normalization approach. Further, as we had anticipated, the neighboring timepoints showed higher gene expression correlation than distant timepoints (Supplementary Figure 5) with Spearman correlation for successive samples consistently equal or above 0.96. This suggests that our sampling was close enough to capture transcriptional dynamics across the sampling time. We also calculated the lower limit detection limit setting our sensitivity to RPG10K of 0.01 which corresponds to ~2 mapped reads. Averaged over the 6 samples, the detection limit was 1038 transcripts per embryo.

### Total mRNA Content of the Embryo and Biological replication validate absolute quantification

During the cDNA synthesis step of our protocol we added an equal amount of internal RNA standards (ERCCs) per number of embryos in order to calibrate read counts and obtain absolute numbers of transcripts per embryo. We computed the total mRNA per embryo by summing all transcripts per gene per embryo and calculating the equivalent in nanograms (**Figure 3 A**). The total mRNA dropped from 1.26 ng/embryo at 1 hpo to 0.61 ng/embryo at 2 hpo and then increased to 1.49 ng/embryo at 3 hpo before dropping to 0.93 ng/embryo at the end of our sampling at 6 hpo. The pattern mirrors the total transcripts per embryo (**Figure 2 D**). The mRNA levels agree with the total RNA yields we obtained per embryo (~33 ng/embryo at 1 hpo and 53 ng/embryo at 6 hpo, Supplementary Table 1) assuming 2-5% of total RNA is polyadenylated. We also compared the volume of *B. oleae* embryos to *Xenopus tropicalis* (0.025 mm^3^ versus 0.268 mm^3^, respectively, giving a volume ratio of 10.7). *X. tropicalis* embryos contain 10 – 15 ng of mRNA/embryo at fertilization[3] which closely matches the *B. oleae* 13.5 ng mRNA/embryo at 1 hpo, after account for the volume ratio.

**Figure 3:**
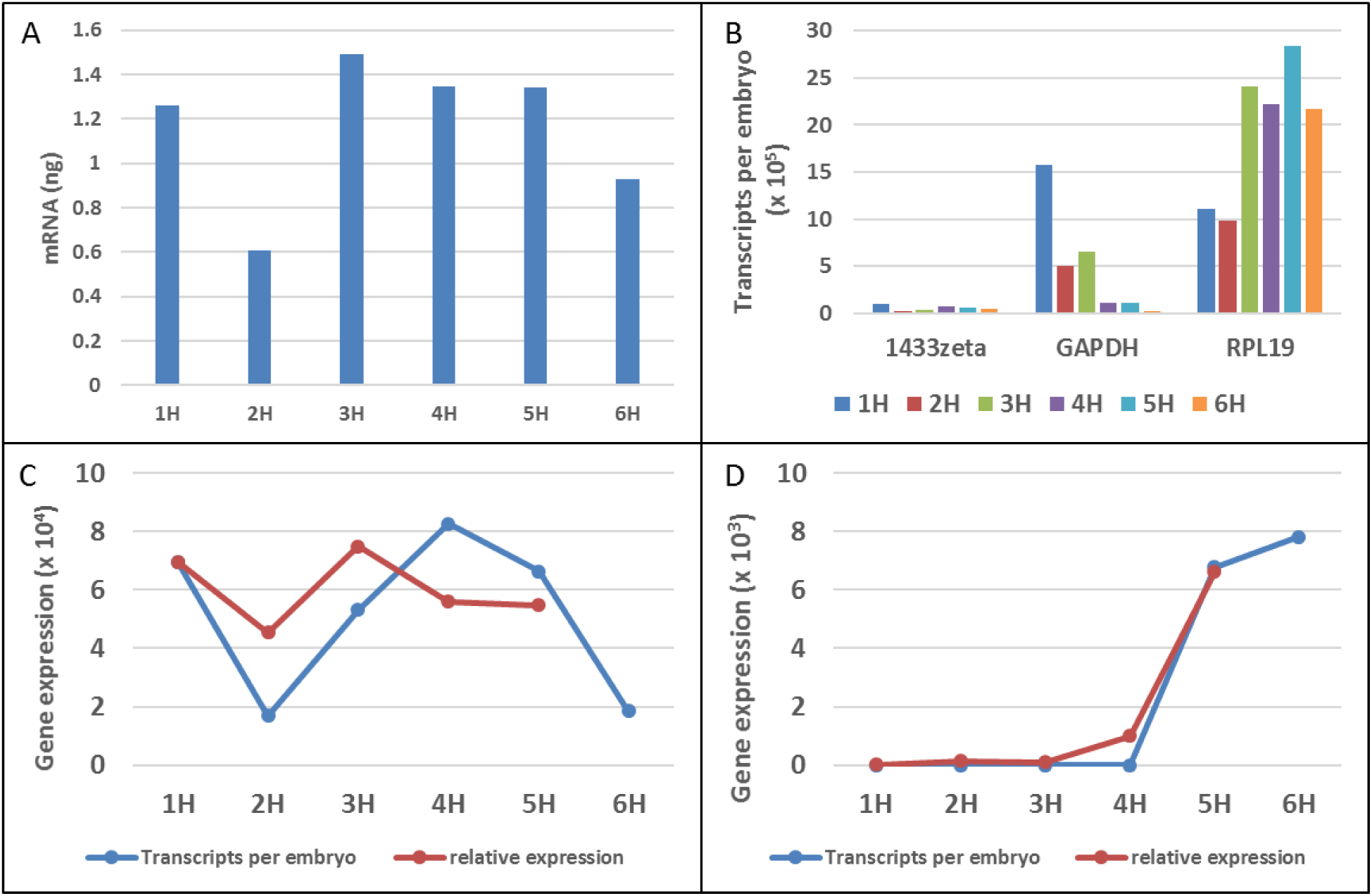
Validation of absolute normalisation. A) Total mRNA (ng) of the embryo across timepoints derived from conversion of transcript molecules to nanograms. B) Absolute gene expression of 3 genes proposed by Sagri et al.[20] as candidate housekeeping genes (14.3.3.zeta, GAPDH, RPL19). C) Absolute expression (blue) and qPCR expression (red) of HID. qPCR expression values were multiplied by a scaling factor. D) Same as C but for SRY.

We further sought to determine whether the expression patterns observed could be replicated in a different set of biological samples and using real-time quantitative PCR (qPCR) which is the current standard method for quantifying gene expression. However, the relative method of qPCR gene expression requires identification of a reference gene set whose expression remains stable across samples. Sagri et al.[20] evaluated several commonly used reference genes and identified 14-3-3zeta and RPL19 *B. oleae D. melanogaster* homologues to be optimal reference genes in *B. oleae* eggs collected at 15 minutes following oviposition. We used our data to determine the variation in expression levels of these common reference genes as well as GAPDH (Supplementary Figure 6). In agreement with Sagri et al., 14-3-3zeta had minimal variability in expression and could be used as reference genes for qPCR. We then evaluated the qPCR expression of 3 genes; SRY, HID, and LINGERER, in a different set of biological replicate samples (skipping 6-hour timepoint) using RPL19 and 14-3-3zeta as reference genes. We observed similar trends of gene expression (particularly with 14-3-3zeta compared with RPL19 (**Figure 3 C** and D).

We then, sought to find other genes that could be used as reference genes. We found 66 genes in poly(A)+ (1–6 hpf) that had less than a 0.4-fold change.

To further explore the expression profiles, we performed principal component analysis (PCA) and hierarchical clustering using the most differentially expressed genes. Projection of the expression onto the first two principle components showed temporal correlation of expression across the different timepoints. The first principle component separated the first 3 timepoints from the last 3 timepoints (**Figure 4**). Hierarchical clustering further showed that the first 3 and last 3 timepoints were separately co-clustered (Figure 5).

**Figure 4:**
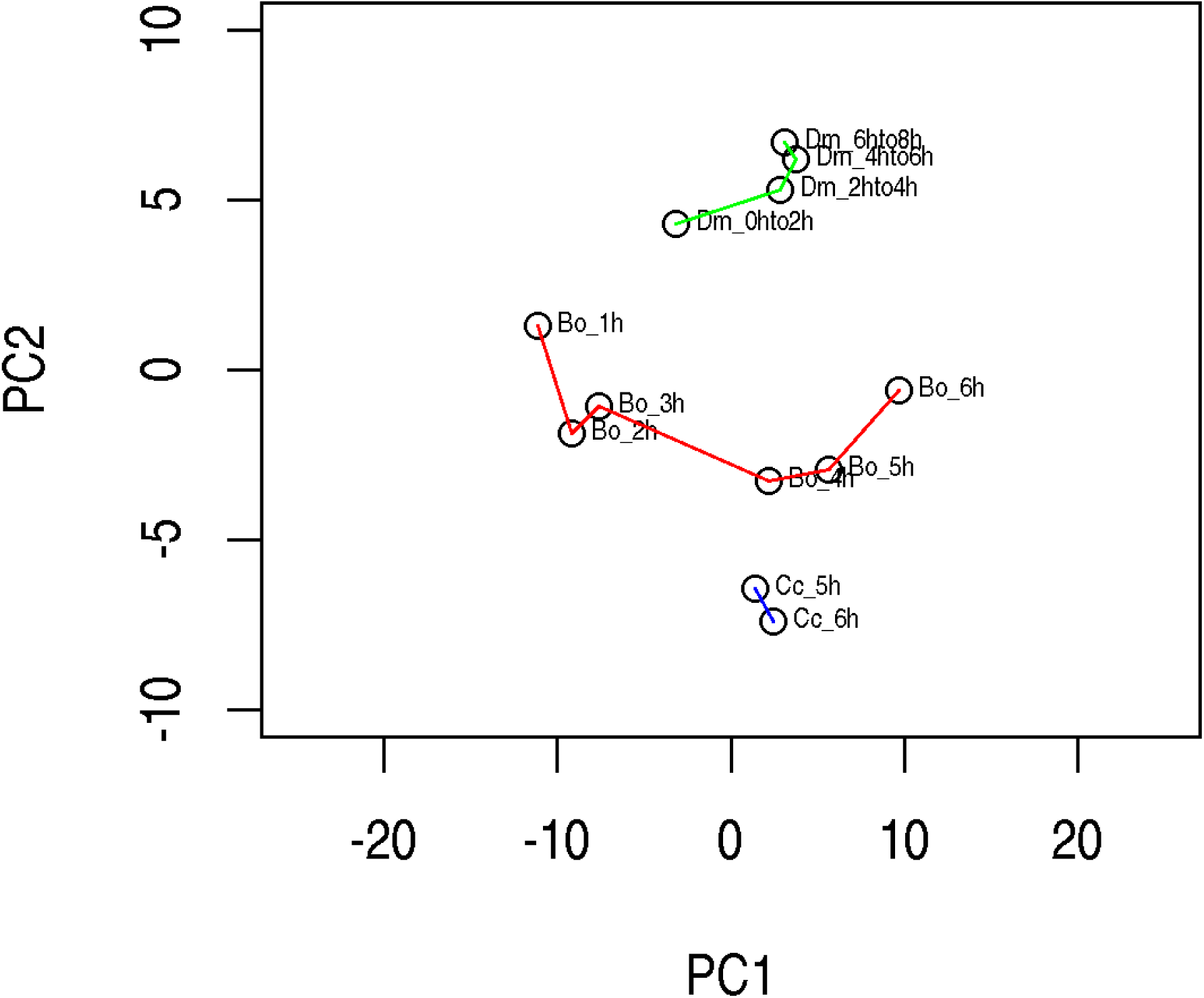
Principal component analysis (PCA) analysis of early embryo development of *B. oleae* (red), *D. melanogaster* (green), and *C. capitata* (blue). PCA was performed using the 100 most variable genes. For each organism the individual points are labelled with the corresponding scientific name initials and the hours post oviposition (Bo_1h for example refers to B. oleae 1 hour post oviposition). Data for *D. melanogaster* was downloaded from the FlyBase.

**Figure 5:**
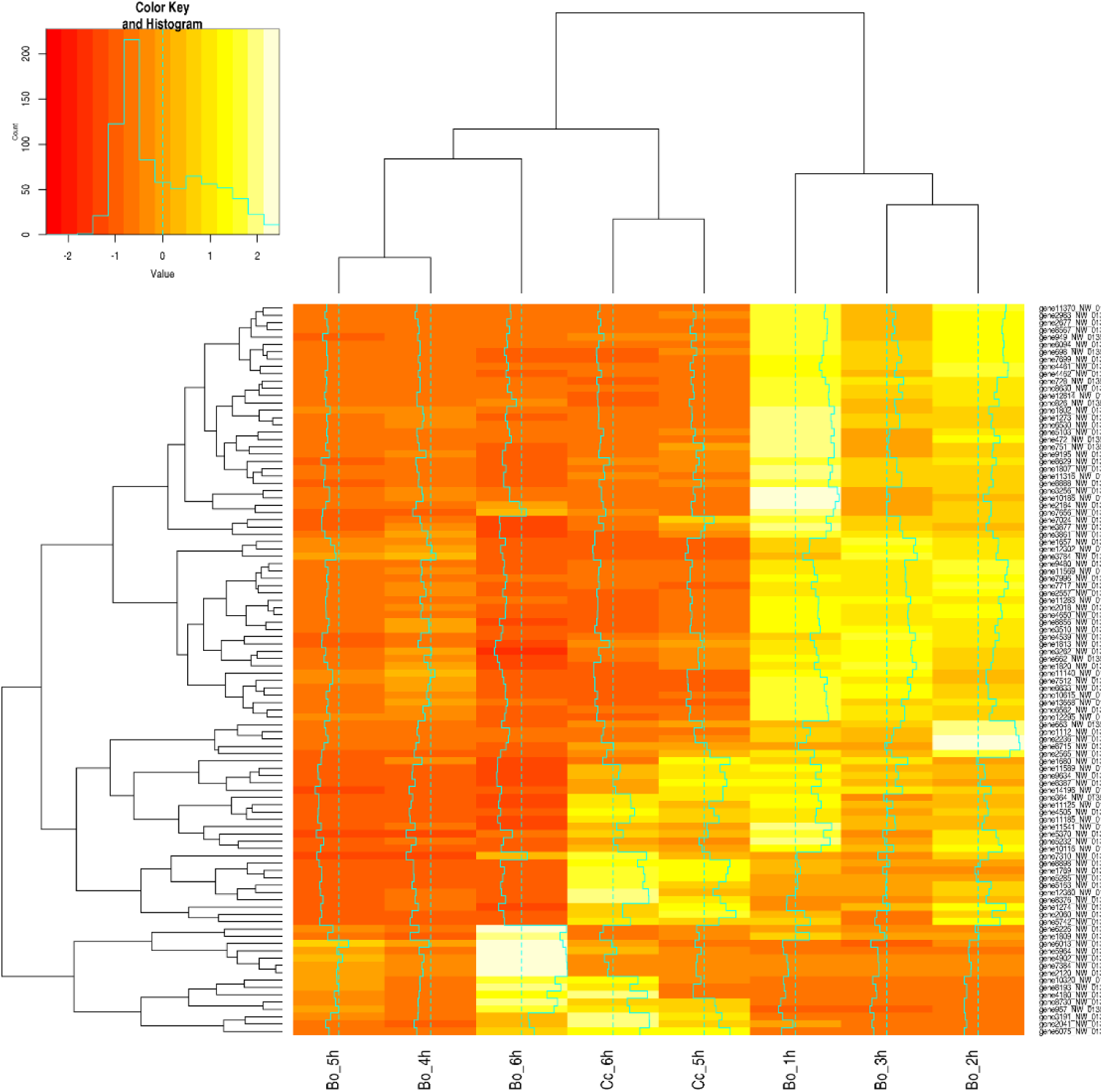
Hierarchical clustering of *B. oleae* embryo timepoints. Clustering was done based on the 100 most variable genes from the 6 different experimental time points. For comparison the gene expression data from the *C. capitata* at 5 hpo and 6 hpo developmental time points were also included in the hierarchical clustering analysis.

We also downloaded *Drosophila melanogaster* RNA-Seq data from Flybase database[21] and projected them together our *B. oleae* data using. Projecting the expressions onto the first principle component showed that *D. melanogaster* embryos at 2 hours post fertilization (hpf) clustered with *B. oleae* 1-3 hpo timepoints. Similarly, the first principle component co-clustered *D. melanogaster* 2-8 hpf and *B. oleae* 4-6 hpo (**Figure 4**). This suggested close similarity in gene expression patterns between the closely related *D. melanogaster* and *B. oleae* Dipteran insects.

### The Maternal-to-Zygotic Transition shows dramatic shift in embryo mRNA content

The developing embryos of many metazoans including echinoderms, nematodes, insects, fish, amphibians, and mammals, are characterized by dramatic transcriptional changes one of which is the change of embryo dependence from maternal to zygotic transcripts; the maternal-to-zygotic transition (MZT, reviewed in Tadros and Lipshitz[22] and Langley et al., [23]). MZT involves 2 main stages; first is the clearance of a large proportion of maternal transcripts and proteins originally loaded into the oocyte during oogenesis, and then initiation of zygotic transcription[22]. In *Drosophila melanogaster* where MZT has been extensively studied, the embryo is dependent on maternal transcripts and proteins up to cell cycle 14 (2 – 3 hours post fertilization, hpf). However, during MZT up to 20% of maternally supplied transcripts are destabilized by maternally encoded proteins by the end of 2 hpf and another 15% of maternal transcripts are destabilized by zygotically encoded protein by 3 hpf[22]. The *D. melanogaster* maternally destabilized genes are enriched in cell cycle functions while maternally stable genes are enriched in house-keeping functions like metabolism, translation.

In *D. melanogaster* still, the timing of the zygotic genome transcription activation depends on absolute time or developmental stage, the main wave of which starts at 2 hpf. Gene ontology search among strictly zygotic genes has shown them to be enriched in transcriptional factors which are probably needed to guide the development of the embryo.

We used our time-course data to elucidate the mechanism of MZT in *B. oleae*, a process that has not been studied before to the best of our knowledge. We identified an interesting phenomenon when examining the total mRNA content per embryo during development across timepoints (**Figure 3 A**). The total mRNA per embryo dropped 51% at 2 hpo compared to levels at 1 hpo and increased 143% at 3 hpo compared to levels at 2 hpo. Interestingly, the distribution of the number of transcripts per gene at different timepoints (Supplementary Figure 7 A) followed a trend similar to the total number of transcripts per embryo suggesting that the clearance of maternal transcripts seen at 2 hpo and the subsequent activation of zygotic gene expression seen at 3 hpo were global events as opposed to targeting specific genes. Indeed, the number of expressed genes was similar between 1,2 and 3 hpo (11043, 10542, and 10713, respectively). We then hypothesized that cleared genes would be enriched with highly expressed genes. Using our time-course data, we performed differential expression between successive timepoints using GFOLD[24] which is designed for samples without biological replicates. GFOLD log2 fold change has been shown to correlate well with qPCR-determined fold change[25]. Genes were coded as upregulated or downregulated using a Gfold cutoff of ±0.5. We identified 1496 genes that were downregulated at 2 hpo compared to 1 hpo. These genes are enriched in maternal-degraded transcripts and are here referred to as maternal-degraded genes. Indeed, at 1 hpo the expression level of maternal-degraded genes was higher than that of the other genes (Supplementary Figure 7 B). The same genes showed similar expression levels at 2 hpo compared to other genes suggesting that these genes were destabilized down to basal levels of other genes (Supplementary Figure 7 C).

We also identified a group of 215 genes enriched in zygotic transcripts (here referred to as zygotic genes). These genes were not detectable at 1 hpo but detectable at any of the other timepoints suggesting that their transcription was from zygotic genome. Since the mRNA content per embryo at 3 hpo had increased 143% compared to 2 hpo, we sought to determine if there was a set of genes that could explain this increase. Maternal-degraded genes, and purely zygotic genes (215 genes whose expression was at 2 or 3 hpo but not at 1 hpo) could not explain this increase as their expression was either similar or much lower than the other genes, respectively (Extended materials).

We performed gene-set enrichment analysis across 3 categories; maternal-degraded genes, zygotic genes, and maternal stable/upregulated genes. The maternal-degraded genes, which were also among the most highly expressed at 1 hpo were enriched in cellular processes, development, and metabolism (Figure 6). The maternal-degraded genes were also enriched in transcription factors for example DREF, BEAF-32A, PNR which are the corresponding Drosophila homologues (**Figure 6**). More like the maternal-degraded genes, the maternal-stable/upregulated genes were enriched in translation, biosynthesis process, gene expression, metabolic processes among others which reflects the high metabolic activity of the rapidly growing embryo (**Figure 6**). DREF transcription factor was also enriched in these genes (**Figure 6**).

**Figure 6:**
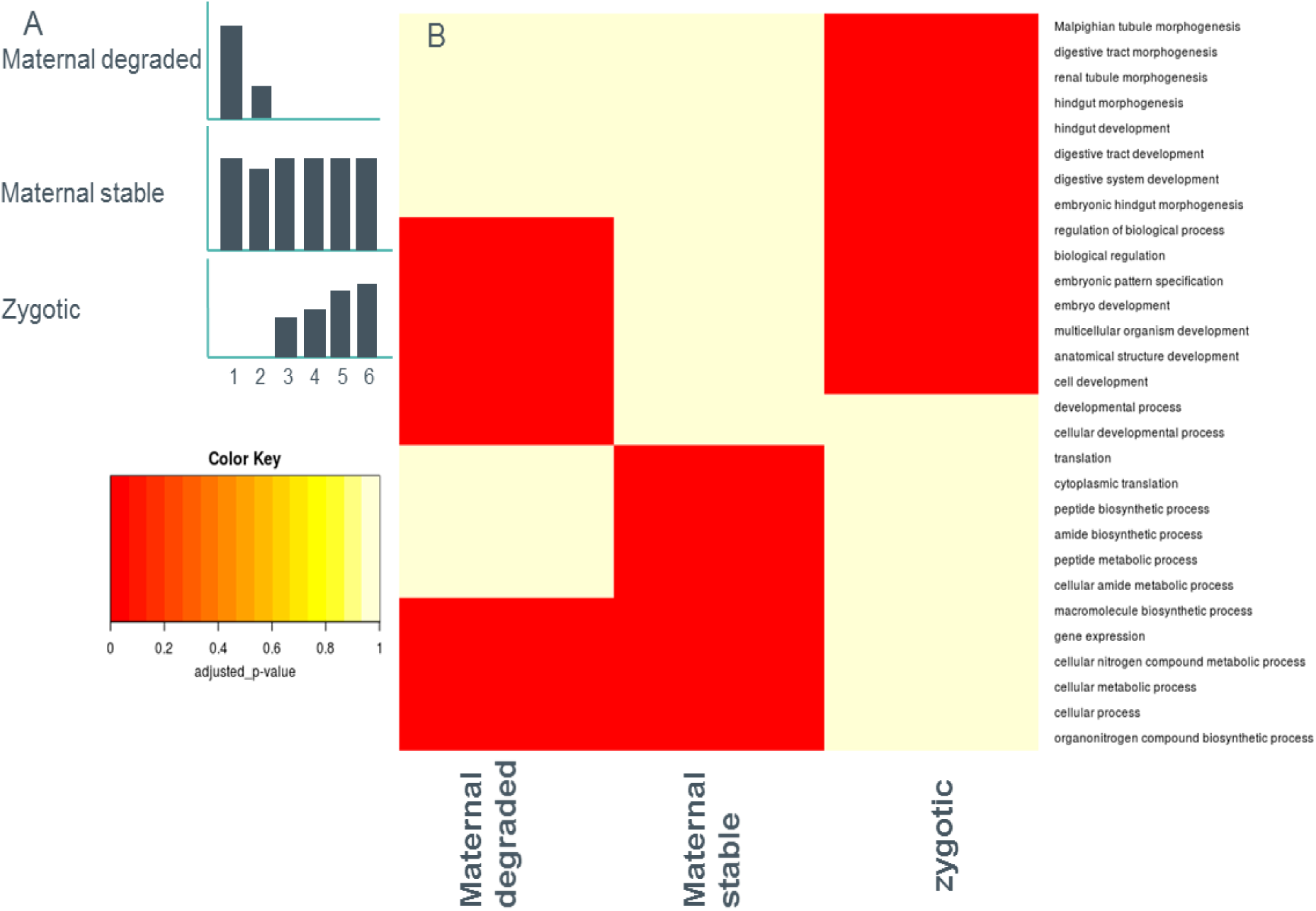
Gene set enrichment analysis of *B. oleae* gene in 3 categories. Gfold[24] was used to perform differential gene expression across the 6 timepoints. A Gfold value of 0.5 was used to identify 2 categories: Maternal degraded (Gfold >0.5 for 1 hpo versus 2 hpo) and Maternal stable (−0.5<Gfold>0.5 for all timepoints). For Zygotic, we identified genes whose expression was detectable at any other timepoint except 1 hpo hpo: hours post oviposition. A) Schematic representation of expression profiles for each of the categories. B) Clustering of gene-set enrichment categories based on the most p-value.

Strikingly, zygotic genes were enriched in specific tissue formation and developing processes including; hindgut development, pattern specification, digestive tract morphogenesis (**Figure 6**).

### Clustering of genes based on temporal expression dynamics

Gene expression is a tightly regulated process. In developing embryos regulation of spatio-temporal dynamics in gene expression is critical for proper organ development. Clustering of genes depending on their temporal expression dynamics not only reduces the complexity of the expression matrix into simple gene groups but can also identify genes with similar biological function as previously shown[26, 27]. Temporal gene expressions have been suggested to follow Guassian distribution[3]. We thus clustered our data using DPGP[28] which jointly models data clusters with a Dirichlet process and temporal dependencies with Gaussian processes. Indeed, we identified genes whose expression peaks at different timepoints, demonstrating transcript kinetics that are highly dynamic and suggesting specific roles for these genes during defined developmental periods (**Figure 7**, Extended material). We further grouped the clusters into 3 groups; 1) genes which peaked at 3 hpo and generally decreased over time, termed early genes (**Figure 7A**), 2) genes whose expression was maintained through 3-5 hpo, termed middle genes (**Figure 7** B and C), and 3) genes whose expression only increased at 5 and/or 6 hpo, termed late genes (**Figure 7** D). Gene-set enrichment showed that indeed, as observed previously under maternal categories of gene, early genes and middle genes were enriched in cellular processes and metabolic process while late genes were enriched in specialized development processes.

**Figure 7:**
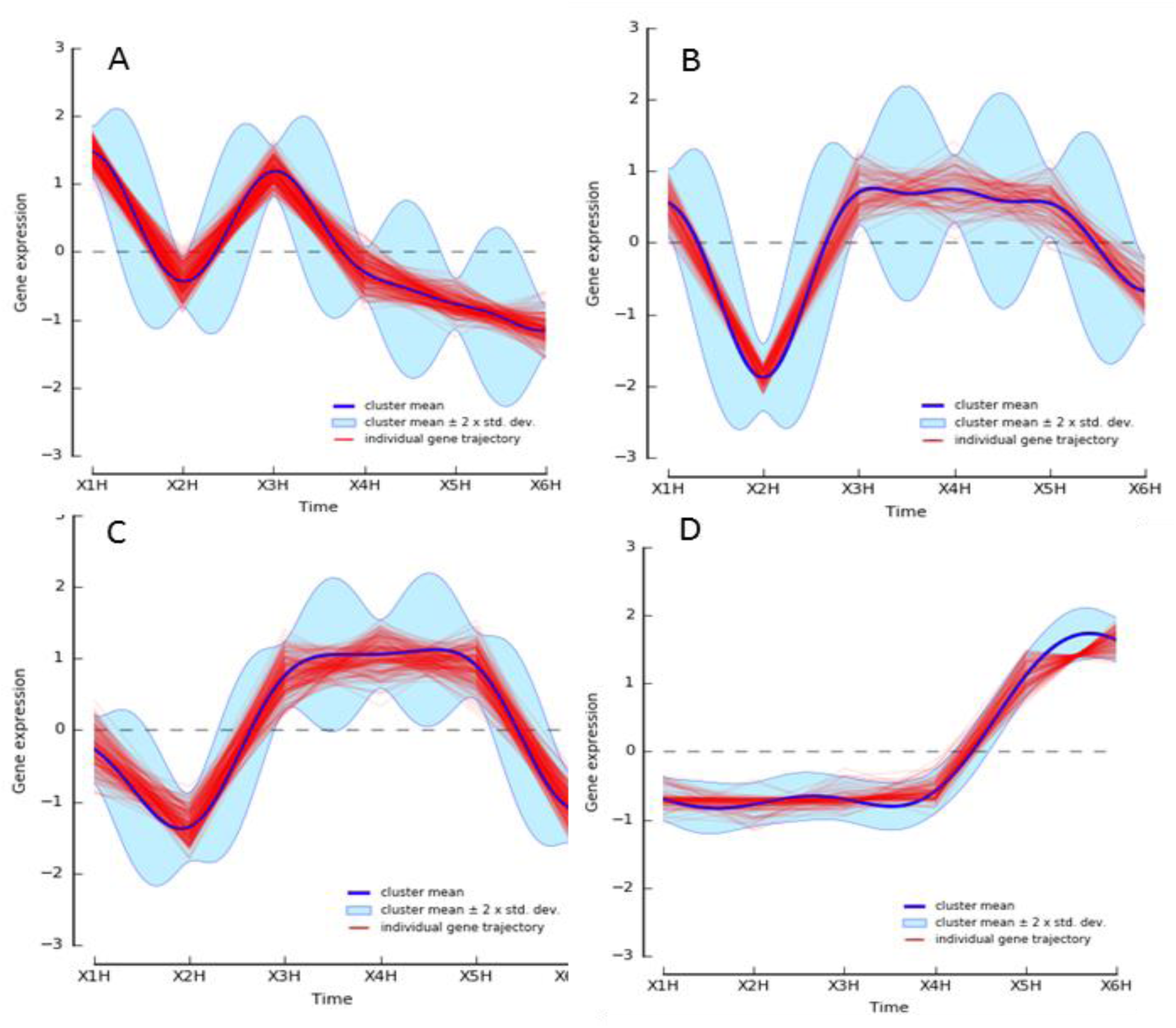
Clustered trajectories of expressed genes across the early embryonic development of *B. oleae*. We used DPGP software[45] to cluster genes according to their temporal expression profile. Out of the 87 clusters returned by DPGP we identified 3 groups of general profiles; early expressed genes (A), stably expressed genes (B and C), and late expressed genes (D). The figures shown here are only samples to show the pattern of expression among the categories we identified. See Supplementary materials for all the clusters.

### Long-read RNA-Seq improves annotation of genes in the sex determination pathway

Dipteran insects share a large part of their sex determination mechanism. In *Drosophila melanogaster* where the sex determination mechanism has been extensively studied (reviewed in[29]), the *sex lethal* gene (*sxl*) acts as the master regulator, mediating sex-specific alternative splicing of both itself and the *transformer* gene (*tra*) depending on ratio of sex chromosomes to autosomes. *tra* in turn mediates sex-specific alternative splicing of *double sex* (*dsx*), the last member of the cascade and the mediator of differential sex development. *B. oleae* homologues of *sxl*, *tra*, and *dsx* have all been identified[30, 31]. However, the master regulator has remained elusive. Sex determination in the olive fly has been suggested to occur within the first 6 hours of embryo development, and to be orchestrated by alternative splicing of *tra* and *dsx* akin to the sex determination mechanism in *D. melanogaster*. In our data, we were able to observe a wide range of alternative splicing for *tra* and *dsx*. We used long-reads from adult male and female heads to identify sex specific isoforms.

In the case of the *dsx* the isoform complexity was significantly different in the early developmental stages compared to the adult ones. In our data we saw isoforms with a different transcription start-site and with a longer length as the prominent isoform present in the early embryonic stages of development (**Figure 8**). These isoforms in the adult head tissue shift to shorter ones with a exon 4 being present in the female but absent from the male (**Figure 8**). Lagos et al [5] have shown that these distinct shorter isoforms are involved in the production of sex-specific proteins important for the sex determination pathway. As we were not able to detect these sex-specific isoforms in the early embryonic stages we argue that their expression starts later during development. Nevertheless, their accumulation at later stages is representative of the sex specification system activated in the early stages of development.

**Figure 8:**
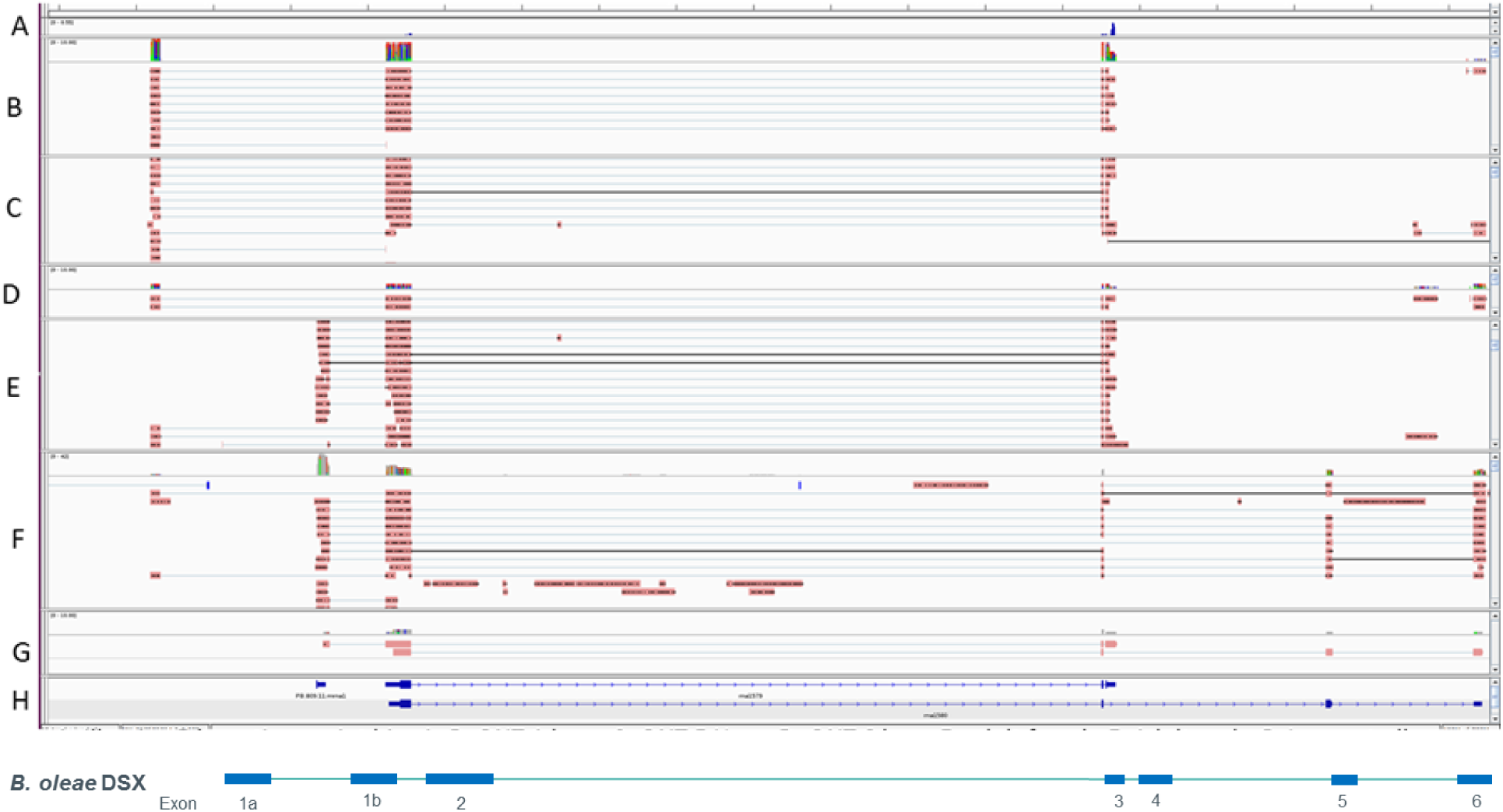
IGV screenshot showing either read alignments to double sex (DSX) gene or isoform models for the DSX gene. Panel A: Short-read Illumina coverage of the 5-hour time point. Panel B - D: ONT sequencing of bulk embryos at 1, 5, 6 hours post oviposition, respectively. Panel E and F: ONT sequencing of female and male adult heads, respectively. Panel G: Male and female specific DSX cDNA as generated by Lagos et al.[30], showing uniform splicing between male and female. H: NCBI predicted gene model for DSX.

## Discussion

Embryonic development has been widely studied in model organisms because of the interest and puzzling nature of the fertilized egg. In insects, embryogenesis has been and continues to be extensively studied in the fruit fly *Drosophila melanogaster*. However, in organisms that are not well characterized long-read RNA-seq has the potential to reveal hitherto unknown genes, correct mis-annotations, and expand isoform diversity. Here, we pooled mixed-sex olive fruit fly (*Bactrocera oleae*) embryos collected at hourly intervals following oviposition for the first six hours of development. This period is interesting to study since previous studies in relatives of *B. oleae* have indicated that the mechanism of sex determination which is mediated by alternative splicing of genes is initiated during this period[2]. Further, evidence from *Ceratitis capitata* suggests that establishment of pole cells, the primordial germ cells, occurs during this period[32]. It is therefore, important to elucidate the transcriptional status during this period. We however, included male and female heads in the generation of the transcriptome (because these were the tissues available to us) in order to expand the transcriptome. cDNA was generated separately from each of the samples using our custom cDNA preparation protocol that enables identification of 5’ and 3’ ends, and addition of ERCC molecular standards to enable absolute quantification of gene expression (see Supplementary material).

We sequenced the cDNA on the MinION (Oxford Nanopore Technologies) using R9.4 1D sequencing chemistry. Details of read yields, alignment statistics, our custom transcriptome assembly workflow, are provided in Extended materials. The *de novo* transcriptome assembly contained 11,883 genes of which 8330 were previously annotated while 3553 are novel genes. Our transcriptomic data provide a rich source of splice variants observed in developing embryos and mature insects, enhancing by 4-fold the isoform models of the current genome assembly. We also provide corrections for 38 genes currently wrongly modelled in the current NCBI annotation. The current *B. oleae* NCBI estimation of 13198 genes is comparable to the number of the closely related *Ceratitis capitata* (14,652[33]). The *C. capitata* genes were however, computationally predicted and thus could face similar limitations as the *B. oleae* predicted gene. The well-studied *D. melanogaster* contains ~17,000 genes. We thus argue that the current *B. oleae* number of genes is underestimated, and our transcriptome provides over 3000 novel genes towards completing the annotation of the genome. Pavlidi et al., [34] previously generated a transcriptome assembly of the olive fly and assembled a total of 14,204 contigs. However, they used a short-read sequencing technology (454 pyrosequencing) and the average contig length was 421 bp compared to an average of 9000 bp in our transcriptome and 9,597 in the NCBI predicted gene models.

Calibration of our gene expression quantification using internal standards enabled absolute quantification of the transcripts per gene. Although the ERCC internal standards exhibit sequencing biases which ultimately affects quantification[3] they have been used successfully to quantify gene expressions. We observed that the total mRNA content of the embryo calculated with absolute normalisation was similar to total yield of cDNA generated per embryo. The remarkable similarity between *B. oleae* total mRNA content and *X. tropicalis* after accounting for embryo volume differences provided further validation of our quantification. Further, the total RNA content of the embryo calculated from absolute normalisation agreed with the actual yield we obtained, thus validating the absolute normalisation method. Further validation of gene expression using qPCR and biological replicates yielded comparable results with our absolute quantified expressions. Furthermore, through the absolute quantification approach we have identified 14-3-3 zeta gene as a suitable normalization controls for qRT-PCR in early embryo samples as previously determined by Sagri et al.

The maternal-to-zygotic transition is a key process in development of many organisms. This process has been extensively studied in *D. melanogaster*. For example, in chromosome ablation and microarray experiments, De Renzis et al.[35], performed a deep characterization the transcriptome of *D. melanogaster* embryos at 0-1 and 2-3 hour post fertilization and provided a rich catalogue of maternal and zygotic genes. However, MZT has not been well studied in other insects. Our sampling captures this period as it covers the events beginning at oviposition (1 hpo) up to the beginning of blastoderm cellularization (6 hpo) when pole cells are already established at the posterior end of the embryo[36]. We show that in *B. oleae*, the MZT is characterized by a dramatic reduction in poly(A) content of embryos, dropping 51% from 1.2 ng/embryo at 1 hpo to 0.6 ng/embryo at 2 hpo and increases 143% to 1.4 ng/embryo at 3 hpo. The distribution of the number of transcripts per gene across timepoints suggested that clearance of poly(A) transcripts and subsequent replenishment are global events as opposed to targeted. Indeed, genes identified as downregulated between 1 and 2 hpo had higher expression at 1 hpo and the same genes had similar expression levels at 2 hpo.

The enrichment of maternal transcripts with translation, cellular process genes, metabolic process genes, and transcription factors like DBREF agrees with the notion that the embryo is dependent on this rich medium for the first 2 hours of development where rapid cell division and expansion occurs [22]. Zygotic genes on the other hand were enriched in developmental process.

However, our approach cannot differentiate between genes directly transcribed from zygotic genome and maternal transcripts which undergo post-transcription modification (for example polyadenylation) and thus making them appear in our quantification. Indeed, in Drosophila, the cytoplasmic poly(A) polymerase encoded by *wispy* promotes poly(A)-tail lengthening during both oocyte maturation and egg activation[37-39]. Therefore, expression profiling methods that do not rely on oligo dT priming or genetic manipulations as for example in De Renzis et al.,[35] who used chromosomal deletion would need to be performed to identify purely zygotic genes in *B. oleae*.

Successful creation of transgenic flies capable of passing on the mutation requires injection of preblastoderm syncytium embryos before establishment of pole cells which eventually become germ cells, typically 20 minutes to 2 hours following oviposition [40-42]. Although the precise timing of pole cell formation has not been reported in olive fly, in the closely related medfly pole cells are established 3-3.5 hours following oviposition[43]. Understanding the structure of genes involved in sex determination at this stage is important as it might reveal avenues of genetic control. Indeed, using a CRISPR-Cas gene drive targeting exon 5 of doublesex gene (*dsx*) in *Anopheles gambiae* enabled near complete suppression of mosquito populations in the lab[42]. *dsx* in *B. oleae* females carries exon 4 which is absent in males and is thus a potential target for genetic control.

We used the full-length reads to improve the annotation of genes in the sex determination pathway. Earlier work by Lagos et al.[30, 31] had identified the most important genes in the sex determination cascade, key components of which rely on alternative splicing. We not only confirm, but also expand the number of isoforms detected, while also being able to characterize them as single molecules rather than as collections of individual PCR products. More importantly, the Y-chromosome localized male determining factor that should be expressed at the early embryonic stages remains elusive. Based on downstream splicing effects we can detect male specific alternative splicing of *tra* from the 5-hour timepoint, which indicates that the sex determining factor must be present in our data. Future efforts will concentrate in its identification and characterization by combining this transcription profiling effort with improved genome assemblies and in particular Y-chromosome assemblies. Overall, our transcription profiling effort provides a rich resource to identify early development genes and transcript isoforms and an extensive set of alternative splicing variants.

## Supporting information

## References

1. Daane, K.M. and M.W. Johnson, Olive fruit fly: managing an ancient pest in modern times. Annu Rev Entomol, 2010. 55: p. 151-69.

2. Morrow, J.L., et al., Expression patterns of sex-determination genes in single male and female embryos of two Bactrocera fruit fly species during early development. Insect Molecular Biology, 2014. 23(6): p. 754-767.

3. Owens, N.D.L., et al., Measuring Absolute RNA Copy Numbers at High Temporal Resolution Reveals Transcriptome Kinetics in Development. Cell Rep, 2016. 14(3): p. 632-647.

4. Angelini, C., D. De Canditiis, and I. De Feis, Computational approaches for isoform detection and estimation: good and bad news. BMC Bioinformatics, 2014. 15: p. 135.

5. Steijger, T., et al., Assessment of transcript reconstruction methods for RNA-seq. Nat Methods, 2013. 10(12): p. 1177-84.

6. Oikonomopoulos, S., et al., Benchmarking of the Oxford Nanopore MinION sequencing for quantitative and qualitative assessment of cDNA populations. Sci Rep, 2016. 6: p. 31602.

7. Weirather, J.L., et al., Comprehensive comparison of Pacific Biosciences and Oxford Nanopore Technologies and their applications to transcriptome analysis. F1000Res, 2017. 6: p. 100.

8. Byrne, A., et al., Nanopore long-read RNAseq reveals widespread transcriptional variation among the surface receptors of individual B cells. Nature Communications, 2017. 8: p. 16027.

9. Clark, M., et al., Long-read sequencing reveals the splicing profile of the calcium channel gene CACNA1C in human brain. bioRxiv, 2018.

10. Liu, H., et al., A high-quality annotated transcriptome of swine peripheral blood. BMC Genomics, 2017. 18(1): p. 479.

11. Singh, N., et al., IsoSeq analysis and functional annotation of the infratentorial ependymoma tumor tissue on PacBio RSII platform. Meta Gene, 2016. 7: p. 70-5.

12. Mavragani-Tsipidou, P., et al., Mitotic and polytene chromosome analysis in Dacus oleae (Diptera: Tephritidae). Genome, 1992. 35(3): p. 373-8.

13. Tsoumani, K.T. and K.D. Mathiopoulos, Genome size estimation with quantitative real-time PCR in two Tephritidae species: Ceratitis capitata and Bactrocera oleae. Journal of Applied Entomology, 2011. 136(8): p. 626-631.

14. Bayega, A., et al., Transcript Profiling Using Long-Read Sequencing Technologies. Methods Mol Biol, 2018. 1783: p. 121-147.

15. Koren, S., et al., Canu: scalable and accurate long-read assembly via adaptive k-mer weighting and repeat separation. Genome Res, 2017. 27(5): p. 722-736.

16. Abdel-Ghany, S.E., et al., A survey of the sorghum transcriptome using single-molecule long reads. Nat Commun, 2016. 7: p. 11706.

17. Wu, T.D. and C.K. Watanabe, GMAP: a genomic mapping and alignment program for mRNA and EST sequences. Bioinformatics, 2005. 21(9): p. 1859-75.

18. Tardaguila, M., et al., SQANTI: extensive characterization of long-read transcript sequences for quality control in full-length transcriptome identification and quantification. Genome Res, 2018.

19. Gao, Y., et al., PRAPI: post-transcriptional regulation analysis pipeline for Iso-Seq. Bioinformatics, 2018. 34(9): p. 1580-1582.

20. Sagri, E., et al., Housekeeping in Tephritid insects: the best gene choice for expression analyses in the medfly and the olive fly. Sci Rep, 2017. 7: p. 45634.

21. FlyBase--the Drosophila database. The FlyBase Consortium. Nucleic Acids Res, 1994. 22(17): p. 3456-8.

22. Tadros, W. and H.D. Lipshitz, The maternal-to-zygotic transition: a play in two acts. Development, 2009. 136(18): p. 3033.

23. Langley, A.R., et al., New insights into the maternal to zygotic transition. Development, 2014. 141(20): p. 3834-41.

24. Feng, J., et al., GFOLD: a generalized fold change for ranking differentially expressed genes from RNA-seq data. Bioinformatics, 2012. 28(21): p. 2782-8.

25. Fu, S., et al., Transcriptome analysis of sweet orange trees infected with 'Candidatus Liberibacter asiaticus' and two strains of Citrus Tristeza Virus. BMC Genomics, 2016. 17: p. 349.

26. Eisen, M.B., et al., Cluster analysis and display of genome-wide expression patterns. Proc Natl Acad Sci U S A, 1998. 95(25): p. 14863-8.

27. Walker, M.G., et al., Prediction of gene function by genome-scale expression analysis: prostate cancer-associated genes. Genome Res, 1999. 9(12): p. 1198-203.

28. McDowell, I.C., et al., Clustering gene expression time series data using an infinite Gaussian process mixture model. PLoS Comput Biol, 2018. 14(1): p. e1005896.

29. Penalva, L.O. and L. Sanchez, RNA binding protein sex-lethal (Sxl) and control of Drosophila sex determination and dosage compensation. Microbiol Mol Biol Rev, 2003. 67(3): p. 343-59, table of contents.

30. Lagos, D., et al., The transformer gene in Bactrocera oleae: the genetic switch that determines its sex fate. Insect Mol Biol, 2007. 16(2): p. 221-30.

31. Lagos, D., et al., Isolation and characterization of the Bactrocera oleae genes orthologous to the sex determining Sex-lethal and doublesex genes of Drosophila melanogaster. Gene, 2005. 348: p. 111-21.

32. Stenpani R. N. D., S.D., and Perondini A. L. P, Early developmental stages of Ceratitis capitata embryos, in Proceedings of the 6th International symposium on fruit flies of economic importance 6-10 May 2002, B.B. N., Editor. 2004, Isteg Scientific Publications, Irene, South Africa: Stellenbosch, South Africa. p. 55-58.

33. Papanicolaou, A., et al., The whole genome sequence of the Mediterranean fruit fly, Ceratitis capitata (Wiedemann), reveals insights into the biology and adaptive evolution of a highly invasive pest species. Genome Biology, 2016. 17(1): p. 192.

34. Pavlidi, N., et al., Analysis of the Olive Fruit Fly Bactrocera oleae Transcriptome and Phylogenetic Classification of the Major Detoxification Gene Families. PLoS One, 2013. 8(6): p. e66533.

35. De Renzis, S., et al., Unmasking activation of the zygotic genome using chromosomal deletions in the Drosophila embryo. PLoS Biol, 2007. 5(5): p. e117.

36. GENÇ, H., Embryonic development of the olive fruit fly, Bactrocera oleae Rossi (Diptera: Tephritidae), in vivo. Turkish Journal of Zoology, 2014. 38: p. 598-602.

37. Benoit, P., et al., PAP- and GLD-2-type poly(A) polymerases are required sequentially in cytoplasmic polyadenylation and oogenesis in Drosophila. Development, 2008. 135(11): p. 1969-79.

38. Cui, J., et al., Wispy, the Drosophila homolog of GLD-2, is required during oogenesis and egg activation. Genetics, 2008. 178(4): p. 2017-29.

39. Cui, J., et al., Cytoplasmic polyadenylation is a major mRNA regulator during oogenesis and egg activation in Drosophila. Dev Biol, 2013. 383(1): p. 121-31.

40. Handler, A.M., A current perspective on insect gene transformation. Insect Biochem Mol Biol, 2001. 31(2): p. 111-28.

41. Fuchs, S., T. Nolan, and A. Crisanti, Mosquito Transgenic Technologies to Reduce Plasmodium Transmission.

42. Kyrou, K., et al., A CRISPR-Cas9 gene drive targeting doublesex causes complete population suppression in caged Anopheles gambiae mosquitoes. Nat Biotechnol, 2018.

43. Riparbelli, M.G., G. Callaini, and R. Dallai, Primordial germ cell migration in the Ceratitis capitata embryo. Tissue Cell, 1996. 28(1): p. 99-105.

44. Salmela, L. and E. Rivals, LoRDEC: accurate and efficient long read error correction. Bioinformatics, 2014. 30(24): p. 3506-14.

45. Kirschneck, C., et al., Valid gene expression normalization by RT-qPCR in studies on hPDL fibroblasts with focus on orthodontic tooth movement and periodontitis. Sci Rep, 2017. 7(1): p. 14751.

