## Supplementary material for "Transcriptome landscape of the developing olive fruit fly embryo delineated by Oxford Nanopore long-read RNA-Seq"

### 1 Materials and methods

#### 1 Sample processing

##### 1.1 Olive fly breeding

The olive fly (*Bactrocera. oleae*) 'Demokritos' strain, that is considered in this study, is maintained in our laboratory for over 15 years and was originally sourced from the Nuclear Research Centre in Athens, Greece. No wild flies have been added since then, hence the strain has maintained a genetic uniformity. Olive flies were reared in appropriate holding cages at 25 ± 1 ◦C, 60 ± 10% relative humidity and 14 L: 10D cycles according to the conditions described in[1].

##### 1.2 Embryo collection, RNA extraction and quality control

In order to explore the transcriptome landscape of the developing embryo *B. oleae* individuals from an inbred isofemale line, were mated with males and then monitored to observe egg laying. Once the eggs were laid the eggs were incubated at room temperature for 1, 2, 3, 4, 5, and 6 hours, respectively followed by RNA extraction using Trizol method. Briefly, the eggs were counted (see

Supplementary Table **1** for number of eggs used per timepoint) and then homogenized in TRI Reagent® (Sigma-Aldrich) and allowed to stand for 5 minutes at room temperature for complete dissociation of nucleoprotein complexes. The samples were then centrifuged at 12,000x g for 15 minutes at 4 °C. The aqueous phase was transferred to a new tube and 0.1 mL of 1-bromo-3-chloropropane (BCP) or 0.2 mL of chloroform per mL of TRI Reagent® added. The sample tubes were covered and shaken vigorously for 15 seconds, and then allowed to stand for 2 – 15 minutes at room temperature.

The resulting mixture was centrifuged at 12,000x g for 15 minutes at 4°C to separate the mixture into 3 phases: a red organic phase (containing protein), an interphase (containing DNA), and a colorless upper aqueous phase (containing RNA). The aqueous phase was transferred to a fresh tube and 0.5 mL of 2-propanol per mL of TRI Reagent used in sample preparation (above) and mixed. Following a 5 – 10 minutes at room temperature incubation, samples were centrifuged at 12,000x g for 10 minutes at 4 °C to collect the RNA precipitate. The pellet was washed by adding a minimum of 1 ml of 75% ethanol per 1 ml of TRI Reagent® used in sample preparation followed by 5 – 10 minutes air-drying. The pellet was finally resuspended in TE buffer for storage.

The quantity of the extracted RNA was determined using a Qubit RNA HS Assay Kit (Thermo Fischer Scientific, Q32852). The quality of the isolated RNA was assessed using an Agilent TapeStation instrument and Agilent RNA ScreenTape kit as per manufacturer's instruction (see Supplementary protocol). The profile of all the total RNA samples showed a single peak at ~2 kb which contrasts with mammalian total RNA profiles where 2 peaks at ~2 kb and 6 kb representing 18S and 28S ribosomal RNA, respectively (Extended Figure 2). This however, is expected for most insects whose 28S rRNA contains a weak hydrogen bonds that easily denatures to release 2 similar sized fragments that run together with the 18S rRNA[2,3]. We therefore, did not consider the RIN of the RNA samples.

**1.3 ERCC Spikes and cDNA synthesis**

For each of the bulk samples, 300 ng of total RNA was used for the cDNA synthesis protocol. ERCC Spike In Mix 1 (Thermo Fischer Scientific, 4456740) were added to the cDNA synthesis master mix (See Supplementary protocol). Our customized and published cDNA synthesis protocol[4] is based on the highly sensitive Smart-seq2 protocol[5], which uses template switching and preamplification. It utilizes a combination of custom reagents and kits and is similar to the methodology tailored to long read sequencing we recently published[6] (See Supplementary protocol for the full step-by-step protocol).

Full-length cDNA was generated using the methods we reported recently[4] (also see Supplementary protocol), followed by sequencing on the ONT MinION sequencer using SQK-LSK108 kit. Due to known PCR biases that result from PCR amplification, we opted to use 12 cycles after performing cycle optimization and noticing that 12 cycles had negligible effect on cDNA profile (Extended Figure 3).

##### 1.4 ONT Library Synthesis and Sequencing

Sequencing libraries (1D) were prepared using ONT SQK-LSK108 protocol. Sequencing was performed using on the ONT MinION Mk1b sequencer and run by MinKNOW version (1.10.16). Flow cells used were R9.4, the library protocol used was 1D. For offline base-calling, Albacore version 2.0.2 was used. Quality analysis of the sequencing and base-called reads was performed using MinIONQC [7] and Pauvre (https://github.com/conchoecia/pauvre) (See Extended Figure 7 for QC results).

##### 1.5 Illumina library preparation and sequencing

The Illumina Trueseq stranded mRNA sample preparation protocol was used to generate the libraries. Briefly, one microgram of total RNA was used to extract poly adenylated transcripts using oligo dT Dynabeads (Invitrogen, USA). Purified RNA was fragmented. First and second strand cDNA synthesis was performed using SuperScrip II. Following adenylation of 3' ends, adapters were ligated and the fragments enriched by PCR using the following cycles: 98°C for 30 seconds, 15 cycles of (98°C for 10 seconds, 60°C for 30 seconds, 72°C for 30 seconds), and a final extension at 72°C for 5 minutes. PCR products were purified using AMPure XP beads (Beckman Coulter). The quality and concentration of cDNA libraries was checked using BioAnalyser DNA-1000 chips (Agilent, USA), and qPCR, respectively. The samples were sequenced on Illumina HiSeq2500 following a 100 bp paired-end sequencing protocol.

#### 2 Data processing

For all the tools used, the examples of parameters used and versions of tools are included in the Supplementary protocol appended at the end of this document.

##### 2.1 Illumina data analysis

Following sequencing, quality control metrics were generated using FastQC (https://www.bioinformatics.babraham.ac.uk/projects/fastqc/). Reads were trimmed using

Cutadapt[8] followed by Trimmomatic[9] processing. Using the NCBI published genome (GCA_001188975.2) and the associated NCBI annotation, alignments to the genome where performed using HISAT2[10] while quantification of gene expression was performed using RSEM[11]. The Transcripts Per Million (TPM) quantification of RSEM was used. ERCCs were used as described in the ONT data analysis to transform relative TPM quantification into absolute number of transcripts per embryo.

#### 2.2 ONT Data processing

The main data analysis software used, and commands run are included in the Supplementary protocol.

#### 2.3 Sequencing QC

The general data processing workflow is shown in Extended Figure 4. Between 3 – 5.5 million reads were generated per sample with the 1-hour timepoint (Bo.E.1H) having the least (3.6 million) and the 5-hour timepoint having the most (5.39 million, Supplementary Table 2). Albacore classifies reads as "Pass" or "Fail" depending on the phred score; greater or less than 7, respectively. In all timepoints 80 – 90 % of reads were classified as "Pass" and were used for further analysis (Extended Figure 5). The total number of bases ranged from 4.8 Gb – 5.8 Gb, of which > 90% belonged to reads classified as "Pass" (Extended Figure 6). Reads classified as "Fail", that is, having a phred score of <7 were not used in our analysis. Interestingly, the mean Q-score among reads equal to or above Q-score 7 decreased graduated over the 48-hour sequencing period while among reads with Q-score below 7, mean Q-score was either variable or constant except for 3- and 6-hour timepoints (Extended Figure 7 B). The mean read length was constant over the 48-hour sequencing period regardless of the read Q-score (Extended Figure 7 B).

#### 2.4 Genome guided *de novo* transcriptome assembly

##### 2.4.1 Comparison of long-read transcriptome assembly tools

We collected synchronously developing mixed sex *B. oleae* embryos at one-hour intervals for the first six hours of development. For each of the 6 timepoints we performed cDNA synthesis using an optimized and customized SMARTer protocol[4] aimed at capturing poly(A)+ RNA (See Supplementary protocol, Supplementary Table 2, Supplementary Table 3). cDNA sequencing libraries were generated following SQK-LSK108 protocol (ONT) and sequenced on the MinION using R9.4 flow cells. We also included cDNA libraries generated from adult male and female heads.

The de novo transcriptome assembly workflow is shown in Supplementary Figure 1. We generated 31 million reads of which 22 million reads (71 %) were error-corrected using Canu. We then focused only on full-length reads identified as those that possessed both 5' adapters and both poly(A) and 3' adapter. These were then pre-processed to orient the strands, trim adapters, and correct errors using short-reads. First, we used 3 million reads generated from 5-hour timepoint to compare 3 genome-guided de novo transcriptome assembly tools; Cupcake ToFU, TAMA, and TAPIS.

Several tools are currently available to enable genome guided *de novo* transcriptome assembly using reads generated from long-read sequencing technologies like PacBio and ONT. However, there's currently no single publication, to our knowledge, that has applied and evaluated these tools using ONT cDNA seq. We therefore, sought to evaluate 3 tools; Cupcake ToFU (https://github.com/Magdoll/cDNA_Cupcake/wiki/Cupcake-ToFU%3A-supporting-scripts-for-Iso-Seq-after-clustering-step), TAMA (https://github.com/GenomeRIK/tama), and TAPIS[12] on ONT *de novo* transcriptome assembly and compare the assemblies to a short-read assembly generated using Cufflinks[13]. We used two tools currently available to analyse long-read de novo transcriptomes; PRAPI[14] and SQANTI[15]. The transcriptome assemblies generated from each tool were evaluated using SQANTI, and their respective annotation files were compared to the NCBI annotation file using Cuffcompare[13]. We used 3.07 million reads (3.29 Gb) taken from the *B. oleae* embryo sample collected at 5 hours post oviposition for which we have both long-read and short-read data. We also used the NCBI genome assembly (GeneBank accession GCA_001188975.2) and NCBI *Bactrocera oleae* Annotation Release 100 as references.

Regarding computational resources we compared ToFU and TAMA directly since they involve similar operations. Cupcake ToFU was more computationally efficient requiring 199.87 minutes wall time (199.81 CPU time) and 12.9 Gb at peak memory compared to TAMA's 6854.9 minutes wall time (6853.5 CPU time) and 226.3 Gb at peak memory to assemble the 3.07 million long-reads. TAPIS required 1046.3 minutes wall time (1097.5 CPU time) and 8.3 Gb at peak memory although TAPIS was run with an option to plot figures which extended run time.

Comparison of annotation files generated by each tool to the NCBI annotation revealed that TAMA had the highest sensitivity (87.5 %, capturing the most among reference transcripts) followed by ToFU at 82.9 %. TAMA also had the lowest precision suggestive of novel features although these can also be artifacts. We filtered ToFU transcripts to retain only transcripts with at least 2 supporting reads, an option that is not available for the other tools tested which might explain the higher precision. The high computational burden of TAMA and low precision of TAPIS discouraged us from continuing with them for generating the *B. oleae* whole transcriptome. Besides, ToFU allows to filter isoforms based on number of supporting reads and has good computational efficiency. We therefore used ToFU for our final assembly.

##### 2.4.2 Comparison of genome guided *de novo* transcriptome assemblies form Illumina short-read versus ONT long-read cDNA-Seq

We sought to compare ToFU *de novo* assembly to an assembly generated from short reads. To control for differences in number of reads and bases generated between short-read and long-read technologies we plotted rare fraction curves to determine the number of reads and bases required to detect the same number of genes between ONT and Illumina. From the rare fraction curves, we estimated that for identification of the same number of genes about 40 times more reads and 8 times more bases were required from Illumina short-read cDNA-Seq compared to ONT long-read cDNA-Seq (Extended Figure 9). We thus used 3.07 million ONT reads (3.29 Gb) and 95.4 million Illumina reads (18.6 Gb).

The Tuxedo genome guided protocol[13] which uses Cufflinks was used to derive the short-read assembly. The Cufflinks and ToFU (long-read) assemblies were compared to the NCBI assembly (

Supplementary **Table 7**). Both assemblies showed comparable sensitivity at base level (83.3 % versus 82.9 %, respectively). However, the long-read assembly showed slightly higher sensitivity at all other levels evaluated. The long-read assembly also showed higher precision at base level (74.9 % versus 65 %), although the short-read assembly had higher precision at other levels evaluated except at locus level. We also analysed the transcriptomes generated using SQANTI. ToFU generated a richer transcriptome containing 43,676 transcripts compared to Cufflinks' 21,840. However, Cufflinks transcripts overlapped 8239 annotated genes compared to ToFU's 6776. Cufflinks transcriptome contain 8712 novel genes, a number which seemed exaggerated in our experience. ToFU transcriptome contained 2060 novel genes which was reasonable to us. ToFU transcriptome however, contained a higher number (14436) and percentage (33%) of transcripts that fully matched an annotated transcript compared to Cufflinks (5962, 27 %, respectively). Interestingly, 80% of genes in the Cufflinks transcriptome had one isoform compared to only 25 % in the ToFU assembly. This shows the power of long-read transcriptome assembly in identifying the full range of splicing patterns and the weakness of short-read RNA-seq in assembling transcriptomes as has been noted by others in the field. Noteworthy, although we used 3.07 million ONT reads, ToFU ignored 1.08 million reads due to low identity, coverage or being unmapped.

##### 2.4.3 *Bactrocera oleae* genome guided long-read transcriptome assembly

Although the olive fly has been annotated by NCBI (*Bactrocera oleae* Annotation Release 100) the annotation was based mainly on computational predictions and support from short-read data. We therefore, sought to provide an assembly based on long reads which would confirm the NCBI predictions, improve on the annotations, and add missing genes, and particularly missing isoforms. We included reads generated from adult female and male heads and sequenced following the same protocol described above. All reads were provided to Canu[16] to perform consensus error correction. Of the 25.7 million total reads generated across 6 timepoints 18.5 million reads (72 %)

were error-corrected which improved alignment identity from ~87% to ~96% (Extended Figure 14). Because we added different adapters at each end of each original cDNA molecule, and because the current MinION performs sequencing randomly starting from either 3' end of each molecule we used a customized version of Mandalorion to return the correct original strand of each molecule based on detection of the 5' and 3' adapters. Only full-length reads were used in transcriptome assembly (9.8 million reads). Full-length reads where identified as those that possessed the both 5' adapters and both poly(A) and 3' adapter. Using this filtering, about 50 % of the reads were classified as full-length (Extended Figure 8). The adapters were trimmed using Porechop. Since we had generated short-read Illumina reads for the 5- and 6-hour timepoints, we used Lordec to perform hybrid error correction which further improved the alignment identity from ~96% to ~98%. Another customized version of Mandalorion was used to perform a final round of adapter trimming. Reads that had not been error corrected using Canu were taken through a similar pre-processing described above and combined with the error-corrected reads. The pre-processed reads were aligned to the genome using GMAP.

We used Cupcake ToFU for transcriptome assembly. ToFU provided adequate user options and reasonable running speed. ToFU was used to collapse the transcripts into a non-redundant set of transcripts comprising the genes and their associated isoforms. SQANTI[15] was then used to analyse the transcripts, identify novel genes, and perform open reading frame prediction using the GeneMarkS algorithm. Because ToFU was set to consider reads that were aligned at least 99 % in length and with at least 95 % identity, out of the total 14.7 million reads, 3.9 million were used to derive the transcriptome. See Extended Figure 11 for a summary of the SQANTI output.

The ToFU collapsed transcripts contained a total of 11,883 genes and 79,810 isoforms. Of the genes, 8330 matched the NCBI annotated genes while 3553 genes were novel. Over 50% of novel genes were mono-exon compared to annotated genes where over 80% of the genes were multi-exon (Extended Figure 11). Novel genes also showed lower expression compared to annotated genes (Extended Figure 11). Structurally, SQANTI categorises the transcripts into 9 groups depending on their splice junction and genomic coordinate including; full splice match (FSM), incomplete splice match (ISM), novel in catalogue (NIC), novel not in catalogue (NNC), genic genomic, antisense, fusion, intergenic, genic intron. Most of ToFU collapsed transcripts were a perfect splice match to the annotated transcripts (32.2 %), followed by transcripts containing alternatively spliced junctions (22.7 %, Extended Figure 11). The length distribution of transcripts across the different structural categories was largely similar (Extended Figure 11). There was however, difference in expression across the structural categories; ISM, NIC, NNC, and fusion transcripts had the lowest expression as measured with short-read RNA-Seq (Extended Figure 11). Regarding splice junctions across structural categories, the antisense transcripts had the highest number of non-canonical splice junctions (Extended Figure 11).

About 1.9% of the reads that passed assembly pre-processing either did not align to the NCBI genome or aligned with less than 51% of their length. These reads were aligned to an inhouse *B. oleae* assembly that is more contiguous. 32% of these reads aligned either with at least 50% coverage. Cupcake ToFU and SQANTI were also run on these aligned reads to further identify genes missing from the NCBI assembly. We identified another 228 genes missing in the NCBI genome but found within our inhouse assembly.

All NCBI predicted proteins and the novel identified ORFs were blasted against Uniprot *Drosophila melanogaster* Swiss-prot and Trembl databases to identify *D. melanogaster* homologues. We updated the NCBI *B. oleae* annotation to include novel genes and thus created a new gene transfer file (GTF) termed Annotation v2. We also added the novel genes that resulted from our inhouse *B. oleae* genome assembly to the NCBI assembly and created a new assembly termed NCBI v2. The Annotation v2 and assembly NCBI v2 were used in gene expression quantification and expression profiling. The full transcriptome generated was also analysed using PRAPI[14].

##### 2.5 Identification of Cricket paralysis virus as a major threat to our fly colony

Out of the 274435 reads (1.9%) that did not align to the NCBI *B. oleae* genome assembly, 187073 reads (68%) also failed to align to our inhouse assembly. We therefore, performed blastx against the Uniprot Swiss-prot database. 37% of these un-aligned reads returned a hit (evalue <= 1e-4). Interestingly, 86% of the hits were to Viruses and among Viruses, the genus Cripavirus of the order Picornvirales accounted for 87% of the hits. Further digging into this category revealed that the blastx hits to the Cripavirus were only to 3 proteins; Structural polyprotein (65%, Uniprot ID

P13418), Replicase polyprotein, ORF1 (24%, Uniprot ID Q9IJX4), and Replicase polyprotein, ORF1 (10%, Uniprot ID O36966).

Cricket paralysis virus (CrPV) has been shown to infect and replicate in adult olive flies[17]. Feeding insects on solution containing CrPV resulted in 80% of the flies dying within 12 days. Identifying a high number of reads corresponding the structural polyprotein which is the precursor of all viral capsid proteins suggests that virus was replicating within the flies we used. Further, Replicase polyprotein, ORF1 (CrPV-1A) is a known suppressor of RNA-mediated gene silencing, an antiviral defense mechanism of insect cells. CrPV-1A obstructs the initial target searching of Ago2-RISC by binding to Argonaute protein which is the core of the RNA-induced silencing complex (RISC) in insects and thus suppress its target cleavage reaction[18].

##### 2.6 Read alignment

The NCBI *B. oleae* assembly (accession code GCA_001188975.2) was used for the alignment. However, since we included ERCC in our cDNA the ERCC sequences were included in the NCBI assembly prior to alignment. Reads were preprocessed to make them stranded and remove adapters and poly(A) tail. All the reads were also supplied to Canu to perform consensus error correction (see Supplementary protocol for run parameters, Supplementary Table 4). Alignment of reads to the reference genome and transcriptome was performed using 2 splice-aware and long-read enabled aligners; GMAP[19] and Minimap2[20] (see Supplementary Table 4 for alignment statistics).

###### 2.6.1 Comparison of GMAP and Minimap2 splice-ware long-read aligners

Alignment of reads generated from third generation long-read sequencing technologies provides unique challenges due to the length of the reads and the relatively high raw read error rates. Alignment of discontinuous reads for example cDNA further complicates the alignment. Currently, GMAP and Minimap2 are the most widely used long-read splice-aware aligners although others like GraphMap[21] support transcript alignment albeit through an annotation file which is used to reconstruct the transcriptome. Recently, Krizanovic et al.,[22] evaluated long-read RNA-seq aligners and found GMAP to be the best aligner for long-read cDNA-Seq. However, Minimap2 lacked splice-aware alignment at the time and thus was not evaluated. We have used RNAseqEval[22] developed by Krizanovic et al., to directly compare GMAP and Minimap2.

Using our Cluster computing system; CentOS6, Linux x86-64 with 32 Gb of memory and 24 cores we aligned 1 million reads subsampled from the 5-hour timepoint to the *B. oleae* NCBI genome (471,863,126 bases, including ERCC sequences) using GMAP and Minimap2 with their default settings except for restriction of secondary alignments and setting the number of threads. Both tools involve an indexing step of the genome although this is optional for Minimap2. Regarding genome indexing, Minimap2 showed exceptional speed taking only 0.52 minutes (wall time) compared to GMAP's 7.06 minutes. Minimap2 indexing also showed better memory usage taking up only 345 Mb of memory and returning a single 1.3 gigabytes index whereas GMAP used 1.3 gigabytes of memory returning an index of ~ 2.5 gibabytes distributed in 15 files. Regarding read alignment Minimap2 showed exceptional speed (for example aligning the 1 million reads in only 4.2 minutes (wall time) with 24 threads) compared to GMAP's 63.5 minutes (Extended Figure 10). Both tools however, showed comparable scaling with the number of processors (threads) used (Extended Figure 10). Running with just 1 thread during alignment, we noticed that GMAP used 1.4 gigabytes at peak memory compared to Minimap2's 3.3 gigabytes. Minimap2 however, includes a '-I' option to adjust the number of reads loaded into memory and this could probably reduce memory usage. Indeed, Chu et al.,[23] who evaluated several genomic DNA mapping tools observed that Minimap (a less advanced version of Minimap2) was the most computationally efficient, specific and sensitive method on ONT datasets tested.

Minimap2 was therefore our preferred method for evaluating alignment statistics due to its speed and higher accuracy. For genome guided *de novo* transcriptome assembly we used GMAP because the assembly tools required GMAP and because GMAP includes options to control alignments; for example --max-intronlength-ends which helps prevent spurious alignments. To determine the alignment statistics, all reads regardless of their adapter content were aligned to the reference using Minimap2. Both custom scripts and AlignQC[24] were used to compute alignment statistics. The read alignment rates reached 98%. Alignment of reads to the *B. oleae* NCBI predicted gene models showed 91% alignment rate. Addition of novel genes identified following our de novo assembly increased alignment rates to gene models from 91% to 97%. However, after filtering for good alignments the alignment rates across timepoints were ~90% (Supplementary Table 4).

We found AlignQC to provide extensive exploration of alignment statistics. Majority of reads aligned were single alignments whereas gapped and chimeric alignment were 0.11 % and 0.76 %, respectively (Extended Figure 12**Error! Reference source not found.**). The median percentage of bases aligned (84 %) was slightly lower than reads aligned. Exons accounted for most of the base alignments (68.5 %) followed by introns, genome, and intergenic at 17 %, 12.4 %, and 6.5 %, respectively. The genome and intergenic fraction probably indicating novel genes and/or DNA contamination in our RNA samples.

As noted elsewhere, long-read sequencing technologies exhibit relatively high error rates compared to short-read sequencing technologies. We observe a median of 16.8 % error rates in aligned segments of reads with deletions contributing the most (7.5%), followed by insertions (4.7 %), and mismatches (4.5 %). Canu correction reduced the error rates from ~16% to ~8% (Supplementary Table 4).

##### 2.7 Relative quantification of gene expression

Relative quantification of gene expression was performed using customized Mandalorion pipeline[25] with the NCBI annotations updated with novel genes (Annotation v2) and assembly NCBI v2 as references. Mandalorion counts the number of reads overlapping exon features of a gene and normalizes for sequencing depth and calculates the relative abundance as Reads Per Gene Per 10,000 aligned reads (RPG10K). There was high correlation between sequenced ERCC internal standards and the expected molecules (Extended Figure 13 A). This was however, expected as we and others have shown that ONT cDNA-Seq shows highly accurate quantification of gene expression[6,24]. We noted that although we added ERCC standards at a constant ratio per embryo across sample the relative normalization showed varying levels at different timepoints (Supplementary Figure 3), perhaps reflecting the varying amount of poly(A) RNA in the embryos across timepoints. We thus opted to perform absolute normalization.

#### 2.8 Direct Absolute normalization of gene expression

The method and justification for absolute normalization have been previously reported by Owens et al.[26]. The method relies on the use of known transcript copy numbers for each ERCC standard and their corresponding relative expressions to derive a conversion factor. The conversion factor is derived from a generalized linear model with a dispersed Poisson likelihood using R statistical software as follows;

$$\text{glm(formula} = r_{qj} \sim \text{offset(log(Sq)), family} = \text{poisson(link} = \text{log))}$$

where:

$r_{qj}$ is the relative abundance (RPG10K) of standard q in sample j

Sq is the known abundance (number of molecules /transcripts) of standard q

The intercept coefficient from the above function is the conversion factor used to convert RPG10K to absolute quantification using the following formula;

$$m_{ij} = \rho_{ij} e^{-\beta j}$$

Where:

$m_{ij}$ is the absolute abundance (number of molecules /transcripts) for gene i in sample j

$\rho_{ij}$ is the relative abundance (RPG10K) of gene i in sample j

$\beta_j$ is the conversion factor

The absolute gene abundances were normalized to the number of embryos used per timepoint to obtain the absolute transcripts per embryo (TPE). Indeed, absolute quantification showed a more representative expression profile than relative quantification when compared to total amount of cDNA generated per embryo (Supplementary Figure 4). The absolute normalized ERCC profiles showed more constant levels across timepoints than relative normalized profiles (Supplementary

Figure 3). This was expected since an equal amount of ERCC standards were added per embryo to each sample. Absolute gene expression was also highly correlated between Illumina short-read cDNA-Seq and ONT long-read cDNA-Seq both for ERCC and genes; Spearman r=0.94 and r=0.9, respectively (Extended Figure 13).

##### 2.9 Detection limits and mRNA content of the embryo

Detection limits were calculated as guided by the ERCC manufacturer (http://tools.thermofisher.com/content/sfs/manuals/cms_086340.pdf). For ONT long-read RNA-Seq we define the detection limit as the number of transcripts per embryo required to produce ~1-2 reads. For Illumina short-read RNA-Seq the detection limit is defined by the number of transcripts per embryo required to produce 10 reads.

#### References


1. Tzanakakis ME, Economopoulos AP, Tsitsipis JA (1967) The importance of conditions during the adult stage in evaluating an artificial food larvae of Dacus oleae (Gmelin) (Diptera: Tephritidae). Zeitschrift für Angewandte Entomologie 59 (1-4):127-130. doi:10.1111/j.1439-0418.1967.tb03846.x
2. Winnebeck EC, Millar CD, Warman GR (2010) Why Does Insect RNA Look Degraded? Journal of Insect Science 10:159. doi:10.1673/031.010.14119
3. Macharia RW, Ombura FL, Aroko EO (2015) Insects' RNA Profiling Reveals Absence of "Hidden Break" in 28S Ribosomal RNA Molecule of Onion Thrips, Thrips tabaci. Journal of nucleic acids 2015:965294. doi:10.1155/2015/965294
4. Bayega A, Wang YC, Oikonomopoulos S, Djambazian H, Fahiminiya S, Ragoussis J (2018) Transcript Profiling Using Long-Read Sequencing Technologies. Methods in molecular biology (Clifton, NJ) 1783:121-147. doi:10.1007/978-1-4939-7834-2_6
5. Picelli S, Bjorklund AK, Faridani OR, Sagasser S, Winberg G, Sandberg R (2013) Smart-seq2 for sensitive full-length transcriptome profiling in single cells. Nature methods 10 (11):1096-1098. doi:10.1038/nmeth.2639
6. Oikonomopoulos S, Wang YC, Djambazian H, Badescu D, Ragoussis J (2016) Benchmarking of the Oxford Nanopore MinION sequencing for quantitative and qualitative assessment of cDNA populations. Scientific reports 6:31602. doi:10.1038/srep31602
7. Lanfear R, Schalamun M, Kainer D, Wang W, Schwessinger B (2018) MinIONQC: fast and simple quality control for MinION sequencing data. Bioinformatics (Oxford, England). doi:10.1093/bioinformatics/bty654
8. Martin M (2011) Cutadapt removes adapter sequences from high-throughput sequencing reads. EMBnetjournal; Vol 17, No 1: Next Generation Sequencing Data Analysis
9. Bolger AM, Lohse M, Usadel B (2014) Trimmomatic: a flexible trimmer for Illumina sequence data. Bioinformatics (Oxford, England) 30 (15):2114-2120. doi:10.1093/bioinformatics/btu170

10. Kim D, Langmead B, Salzberg SL (2015) HISAT: a fast spliced aligner with low memory requirements. Nature methods 12 (4):357-360. doi:10.1038/nmeth.3317
11. Li B, Dewey CN (2011) RSEM: accurate transcript quantification from RNA-Seq data with or without a reference genome. BMC bioinformatics 12:323. doi:10.1186/1471-2105-12-323
12. Abdel-Ghany SE, Hamilton M, Jacobi JL, Ngam P, Devitt N, Schilkey F, Ben-Hur A, Reddy AS (2016) A survey of the sorghum transcriptome using single-molecule long reads. Nature communications 7:11706. doi:10.1038/ncomms11706
13. Trapnell C, Roberts A, Goff L, Pertea G, Kim D, Kelley DR, Pimentel H, Salzberg SL, Rinn JL, Pachter L (2012) Differential gene and transcript expression analysis of RNA-seq experiments with TopHat and Cufflinks. Nature Protocols 7 (3):562-578. doi:10.1038/nprot.2012.016
14. Gao Y, Wang H, Zhang H, Wang Y, Chen J, Gu L (2018) PRAPI: post-transcriptional regulation analysis pipeline for Iso-Seq. Bioinformatics (Oxford, England) 34 (9):1580-1582. doi:10.1093/bioinformatics/btx830
15. Tardaguila M, de la Fuente L, Marti C, Pereira C, Pardo-Palacios FJ, Del Risco H, Ferrell M, Mellado M, Macchietto M, Verheggen K, Edelmann M, Ezkurdia I, Vazquez J, Tress M, Mortazavi A, Martens L, Rodriguez-Navarro S, Moreno-Manzano V, Conesa A (2018) SQANTI: extensive characterization of long-read transcript sequences for quality control in full-length transcriptome identification and quantification. Genome research. doi:10.1101/gr.222976.117
16. Koren S, Walenz BP, Berlin K, Miller JR, Bergman NH, Phillippy AM (2017) Canu: scalable and accurate long-read assembly via adaptive k-mer weighting and repeat separation. Genome research 27 (5):722-736. doi:10.1101/gr.215087.116
17. Manousis T, Moore NF (1987) Cricket Paralysis Virus, a Potential Control Agent for the Olive Fruit Fly, Dacus oleae Gmel. Applied and environmental microbiology 53 (1):142-148
18. Watanabe M, Iwakawa HO, Tadakuma H, Tomari Y (2017) Biochemical and single-molecule analyses of the RNA silencing suppressing activity of CrPV-1A. Nucleic acids research 45 (18):10837-10844. doi:10.1093/nar/gkx748
19. Wu TD, Watanabe CK (2005) GMAP: a genomic mapping and alignment program for mRNA and EST sequences. Bioinformatics (Oxford, England) 21 (9):1859-1875. doi:10.1093/bioinformatics/bti310
20. Li H (2018) Minimap2: pairwise alignment for nucleotide sequences. Bioinformatics (Oxford, England). doi:10.1093/bioinformatics/bty191
21. Sovic I, Sikic M, Wilm A, Fenlon SN, Chen S, Nagarajan N (2016) Fast and sensitive mapping of nanopore sequencing reads with GraphMap. Nature communications 7:11307. doi:10.1038/ncomms11307
22. Krizanovic K, Echchiki A, Roux J, Sikic M (2018) Evaluation of tools for long read RNA-seq splice-aware alignment. Bioinformatics (Oxford, England) 34 (5):748-754. doi:10.1093/bioinformatics/btx668
23. Chu J, Mohamadi H, Warren RL, Yang C, Birol I (2017) Innovations and challenges in detecting long read overlaps: an evaluation of the state-of-the-art. Bioinformatics (Oxford, England) 33 (8):1261-1270. doi:10.1093/bioinformatics/btw811
24. Weirather JL, de Cesare M, Wang Y, Piazza P, Sebastiano V, Wang XJ, Buck D, Au KF (2017) Comprehensive comparison of Pacific Biosciences and Oxford Nanopore Technologies and their applications to transcriptome analysis. F1000Research 6:100. doi:10.12688/f1000research.10571.2
25. Byrne A, Beaudin AE, Olsen HE, Jain M, Cole C, Palmer T, DuBois RM, Forsberg EC, Akeson M, Vollmers C (2017) Nanopore long-read RNAseq reveals widespread transcriptional variation among the surface receptors of individual B cells. Nature communications 8:16027. doi:10.1038/ncomms16027
26. Owens NDL, Blitz IL, Lane MA, Patrushev I, Overton JD, Gilchrist MJ, Cho KWY, Khokha MK (2016) Measuring Absolute RNA Copy Numbers at High Temporal Resolution Reveals Transcriptome Kinetics in Development. Cell reports 14 (3):632-647. doi:10.1016/j.celrep.2015.12.050

27. Sagri E, Koskinioti P, Gregoriou ME, Tsoumani KT, Bassiakos YC, Mathiopoulos KD (2017) Housekeeping in Tephritid insects: the best gene choice for expression analyses in the medfly and the olive fly. Scientific reports 7:45634. doi:10.1038/srep45634

#### Tables

**Supplementary Table 1.** Number of embryos used at the hourly intervals of the early development in *B. oleae* and the corresponding total RNA per embryo.

| Hours post oviposition (hpo) | Organism | ID | No. of embryos | Total RNA/embryo (ng) |
|---|---|---|---|---|
| 1 | *Bactrocera oleae* | Bo.E.1H | 300 | 33.3 |
| 2 | *Bactrocera oleae* | Bo.E.2H | 144 | 33.3 |
| 3 | *Bactrocera oleae* | Bo.E.3H | 108 | 63.9 |
| 4 | *Bactrocera oleae* | Bo.E.4H | 100 | 50 |
| 5 | *Bactrocera oleae* | Bo.E.5H | 110 | 56.4 |
| 6 | *Bactrocera oleae* | Bo.E.6H | 109 | 53.2 |
| Adult female | *Bactrocera oleae* | Bo.Head.female | - | |
| Adult male | *Bactrocera oleae* | Bo.Head.Male | - | |

**Supplementary Table 2.** ONT MinION sequencing statistics for reads generated from the 6 *B. oleae* experiments. The number of reads from the fail and pass categories as well as their sum are presented.

| | **Total reads** | **Fail reads** | **Pass reads** | **mean length** | **median length** | **Maxlength** |
|---|---|---|---|---|---|---|
| Bo_1H | 3590509 | 454769 | 3135740 | 1432 | 1187 | 28385 |
| Bo_2H | 4285167 | 456721 | 3828446 | 1089 | 912 | 120093 |
| Bo_3H | 3720378 | 524711 | 3195667 | 1142 | 940 | 63619 |
| Bo_4H | 4821248 | 504368 | 4316880 | 1271 | 1050 | 200175 |
| Bo_5H | 5333014 | 832579 | 4500435 | 1062 | 884 | 155372 |
| Bo_6H | 4102977 | 693981 | 3408996 | 1060 | 874 | 54207 |

**Supplementary Table 3.** ONT MinION sequencing statistics for bases generated from the 6 *B. olea* experiments. The number of bases of the fail and pass reads as well as their sum are presented.

| | Pass bases | Fail bases | Total bases |
|---|---|---|---|
| Bo_1H | 4,492,856,391 | 295,887,224 | 4,788,743,615 |
| Bo_2H | 4,170,815,792 | 247,454,600 | 4,418,270,392 |
| Bo_3H | 3,650,043,547 | 331,774,650 | 3,981,818,197 |
| Bo_4H | 5,486,943,583 | 289,284,771 | 5,776,228,354 |
| Bo_5H | 4,783,466,942 | 421,877,054 | 5,205,343,996 |
| Bo_6H | 3,614,582,591 | 411,162,173 | 4,025,744,764 |

**Supplementary Table 4. Alignment statistics**. Alignments were performed using Minimap2. The alignments were analysed using AlignQC to derive the statistics in the table.

| | No. pass reads (x $10^6$) | Percentage of reads aligned | Percentage of bases aligned | % reads to ERCC | Error rate (%) | Error rate after Canu (%) |
|---|---|---|---|---|---|---|
| **Bo.E.1H** | **3.1** | **94.2** | **89.3** | **8.6** | **16.6** | **8.0** |
| **Bo.E.2H** | **3.8** | **92.6** | **89** | **16.3** | **16.3** | **7.9** |
| **Bo.E.3H** | **1.3** | **90.4** | **92.9** | **7.5** | **15.9** | **7.9** |
| **Bo.E.4H** | **2.3** | **90.0** | **93.7** | **8.2** | **15.4** | **7.0** |
| **Bo.E.5H** | **3.7** | **89.1** | **93.4** | **8.1** | **14.4** | **7.5** |
| **Bo.E.6H** | **3.4** | **89.3** | **85.0** | **11.3** | **17.0** | **8.8** |

**Supplementary Table 5. Structural categories of the transcriptome.** The terms are adapted from Tardaguila et al.[16]

| Category | Explanation |
|---|---|
| **Transcripts matching already annotated (reference) genes** | |
| **Full splice match (FSM)** | Transcripts matching all reference slice junctions |
| **Incomplete splice match (ISM)** | Transcripts matching consecutive, but not all, reference splice junctions |
| **Categories of Novel transcript found in annotated genes** | |
| **Novel in catalogue (NIC)** | Transcripts with new combinations of splice junctions already in the reference |
| **Novel not in catalogue (NNC)** | Transcripts with donor and/or acceptor combinations not found in the reference |
| **Categories of Novel transcript not found in annotated genes** | |
| **Genic Genomic** | Transcripts with partial overlap with partial overlap in the reference exon |
| **Genic intron** | Transcripts entirely lying within a reference intron |
| **Intergenic** | Transcripts without overlap with any reference genes |
| **Antisense** | Any poly(A) containing transcript overlapping the complementary strand of an annotated reference gene |
| **Fusion** | Transcripts the overlapping two or more annotated genes |

**Supplementary Table 6. Miss-annotated genes.** The *de novo* transcriptome assembly was analysed using PRAPI which led to the identification of 63 genes previously miss-annotated. The gene-pairs that were previously annotated as separate groups but found to be isoforms of the same gene are shown together separated by a hyphen in the table.

| | | | |
|---|---|---|---|
| gene4676_gene4675 | gene10059_gene10058 | gene6700_gene6701 | gene2078_gene2079 |
| gene11616_gene11615 | gene11756_gene11755 | gene7160_gene7161 | gene3823_gene3822 |
| gene3016_gene3017 | gene13068_gene13069 | gene8421_gene8422 | gene4812_gene4813 |
| gene3697_gene3696 | gene196_gene197 | gene8719_gene8718_gene8717 | gene6360_gene6358_gene6359 |
| gene4832_gene4831 | gene2106_gene2105 | gene8837_gene8838 | gene6966_gene6967 |
| gene4869_gene4866_gene4867_gene4868 | gene235_gene234_gene236 | gene12204_gene12203 | gene8604_gene8605 |
| gene7548_gene7547 | gene4803_gene4802 | gene11572_gene11573 | Gene9199_gene9200 |
| Gene285_gene286 | | | |

**Supplementary Table 7. Comparison of short-read and long-read genome guided *de novo* transcriptome assemblies.** Cufflinks[13] and Cupcake ToFU were used to generate the short-read and long-read transcriptome assemblies, respectively. Cuffcompare from Cufflinks was used to compare the gene models to the NCBI assembly.

| | Sensitivity (%) | | Precision (%) | |
|---|---|---|---|---|
| | Short-read | Long-read | Short-read | Long-read |
| Base level | 83.3 | 82.9 | 65 | 74.9 |
| Exon level | 61.4 | 65.4 | 54.9 | 46.1 |
| Intron level | 74.9 | 75.3 | 85 | 69.9 |
| Intron chain level | 41.1 | 49.3 | 44.1 | 25.9 |
| Transcript level | 42 | 50.5 | 26.4 | 19.9 |
| Locus level | 62 | 74.4 | 29.4 | 60.2 |

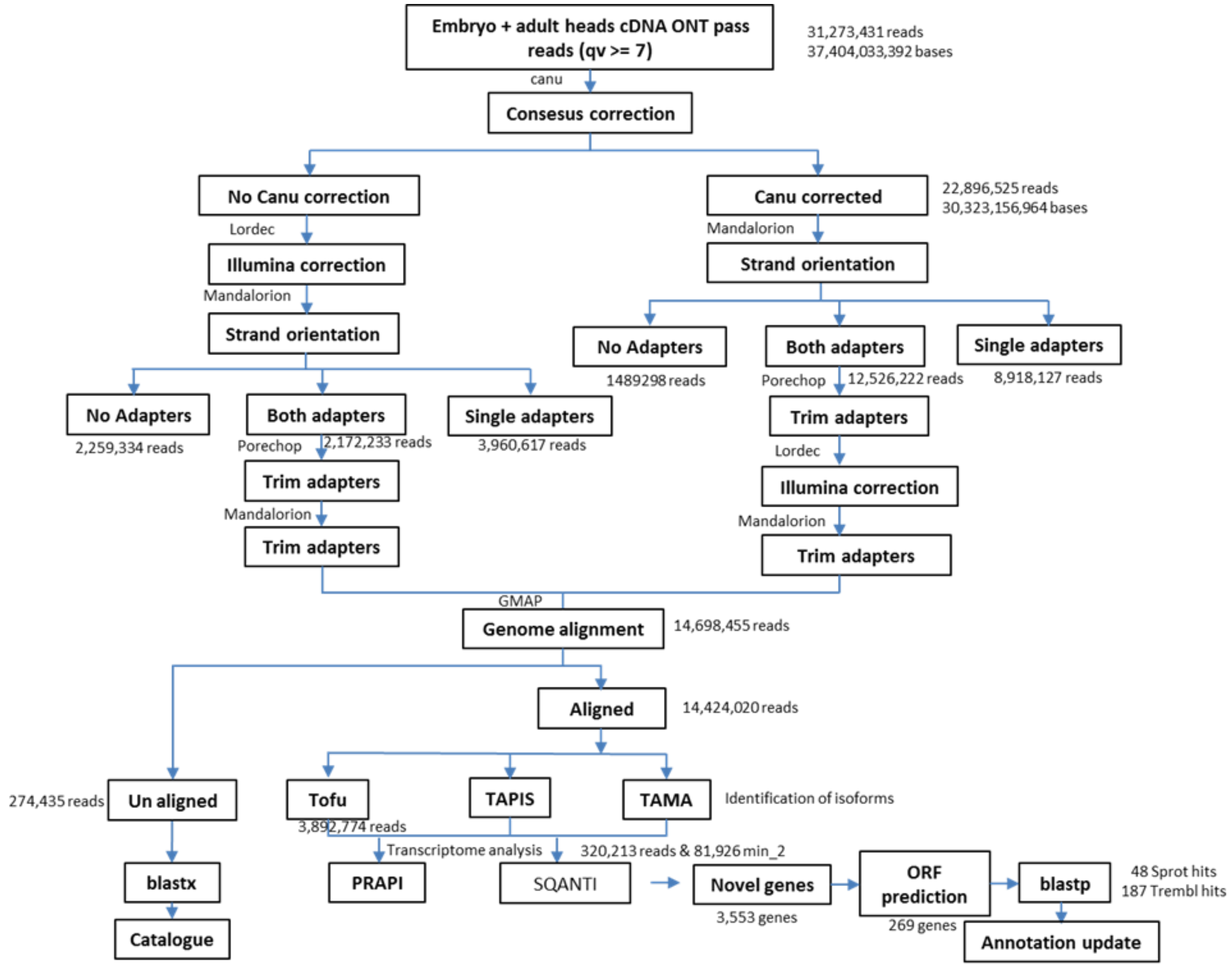


**Supplementary Figure 1.** Transcriptome assembly workflow. All reads were provided to Canu to perform consensus error correction. A customized version of Mandalorion was used to return the correct original strand of each molecule based on detection of the 5’ and 3’ adapters. Only reads with both 5’ and 3’ adapters detected were used in transcriptome assembly. The adapters were trimmed using Porechop. Short-read Illumina reads were used to perform hybrid error correction with Lordec. Another customized version of Mandalorion was used to perform a final round of adapter trimming. Reads that had not been error corrected using Canu were taken through a similar pre-processing described above and combined with the error-corrected reads. The pre-processed reads were aligned to the genome using GMAP. ToFU was used develop a non-redundant trascriptome followed by transcriptome analysis using SQANTI and PRAPI. TAMA, and TAPIS were also evaluated for transcriptome assembly.

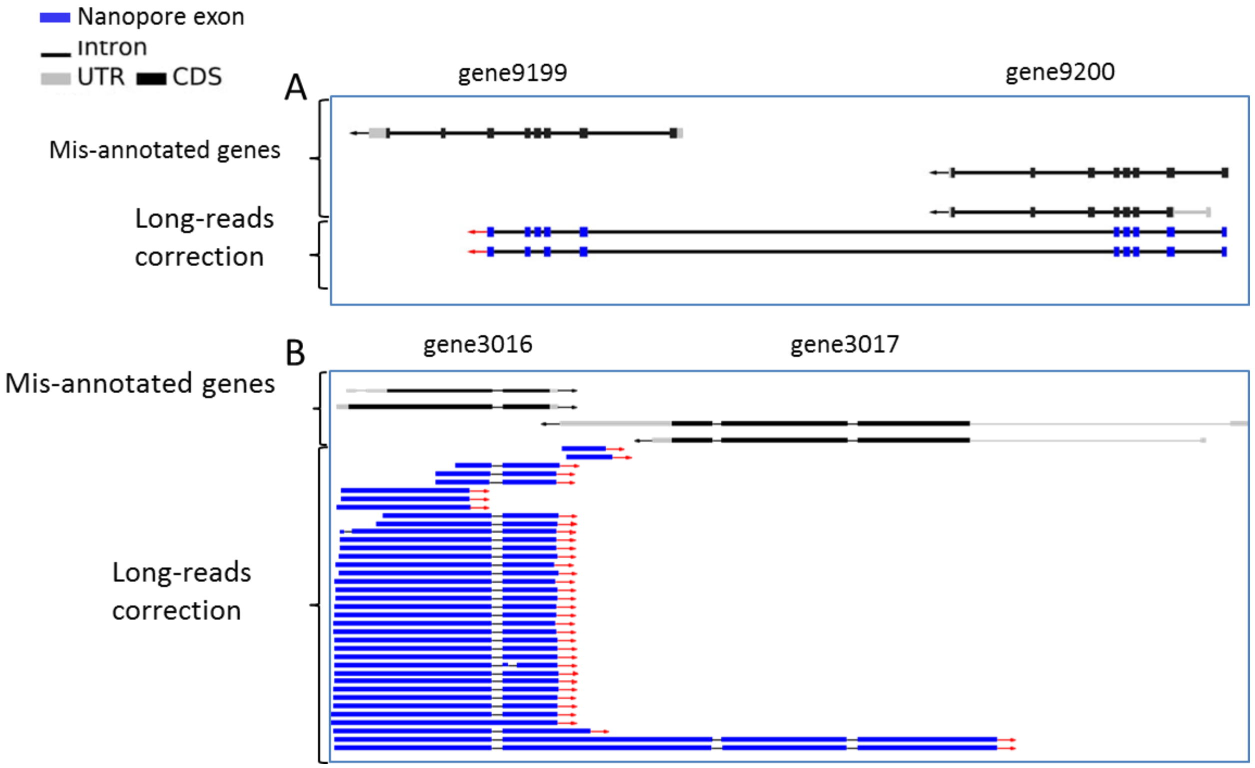


**Supplementary Figure 2.** Identification of miss-annotated genes. PRAPI was used to align transcripts to the predicted genome models and identify genes previously annotated as separate genes but with long-read evidence that they were isoforms of the same gene. Two examples of such genes are shown here (A and B). The miss-annotated isoforms are shown on top while the long-read alignments are shown at the bottom of each panel.

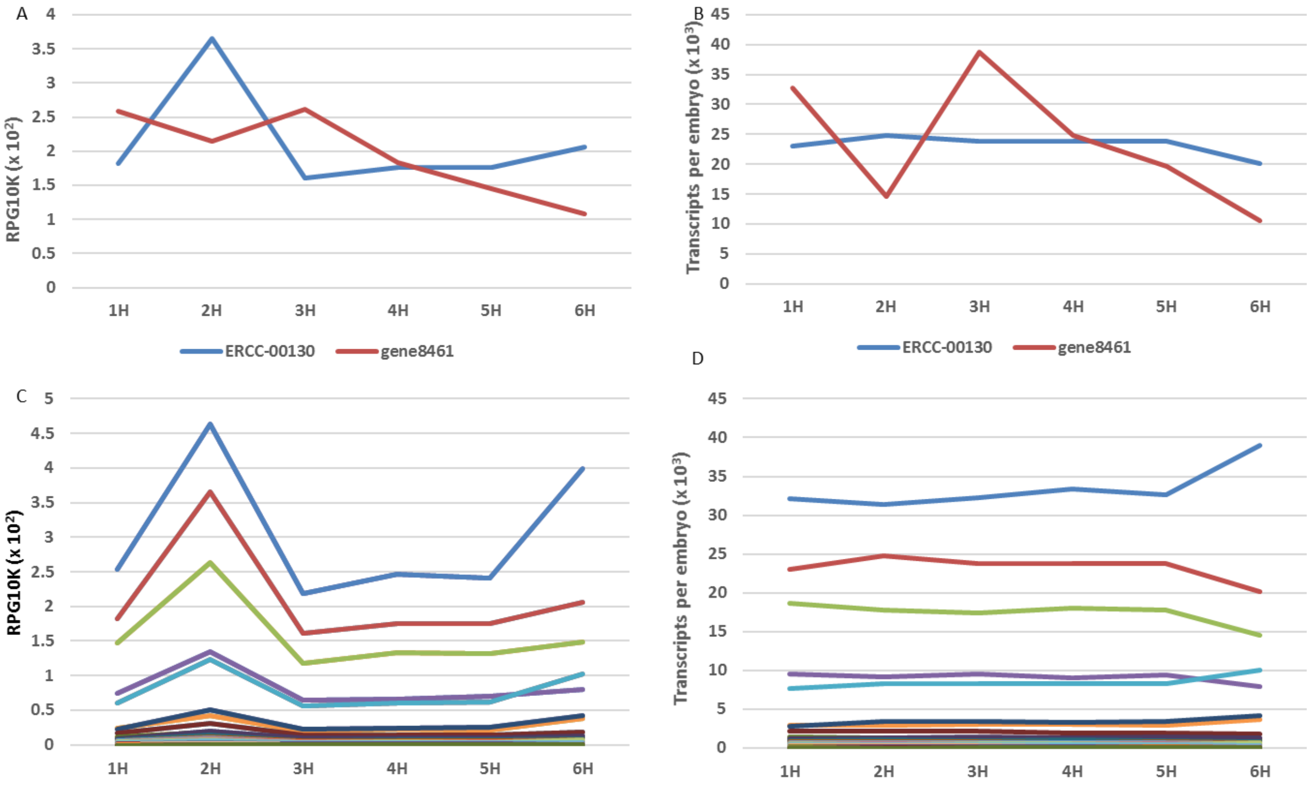


**Supplementary Figure 3.. Comparison of relative normalization and absolute normalization of ERCC internal standards.** A) Relative normalization of the most abundant ERCC (ERCC00130, blue) and a randomly picked gene (gene8461, red). Abundances of the RNA standard varied with time most likely due to changes in the amount of poly(A) RNA in the embryo. B) Same as A but showing absolute normalization of the most abundant ERCC (ERCC00130, blue) and a randomly picked gene (gene8461, red). Here, the abundance of the ERCC is stabilized across timepoints. C) Same as A but including all ERCC internal controls and excluding gene8461. D) Same as C but showing absolute normalization.

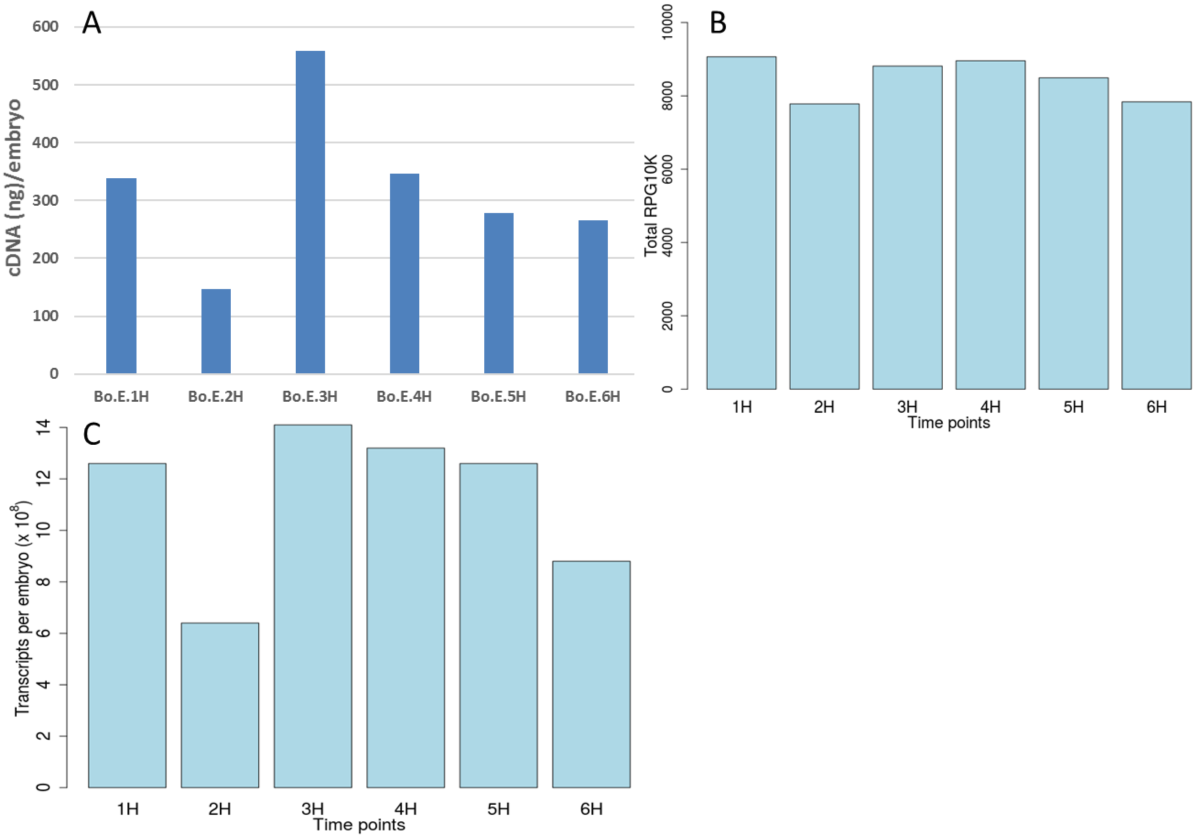


**Supplementary Figure 4. Comparison of relative normalization and absolute normalization of genes.** (A) Amount of synthesized cDNA per embryo. Equal amount of total RNA was used during cDNA synthesis. The amplified cDNA generated was purified and normalized to the number of embryos. B) Summed expression values for all genes across timepoints. The relative method of quantifying gene expression was used. Here, read counts aligning to a gene are normalized by the total reads aligned to all other genes and further normalized to 10000 reads (RPG10K). This profile does not closely resemble the total cDNA profile in A. C) Same as B but using the absolute normalization and normalizing for the number of embryos. This profile shows close resemblance to the cDNA profile, demonstrating the advantage of absolute normalization.

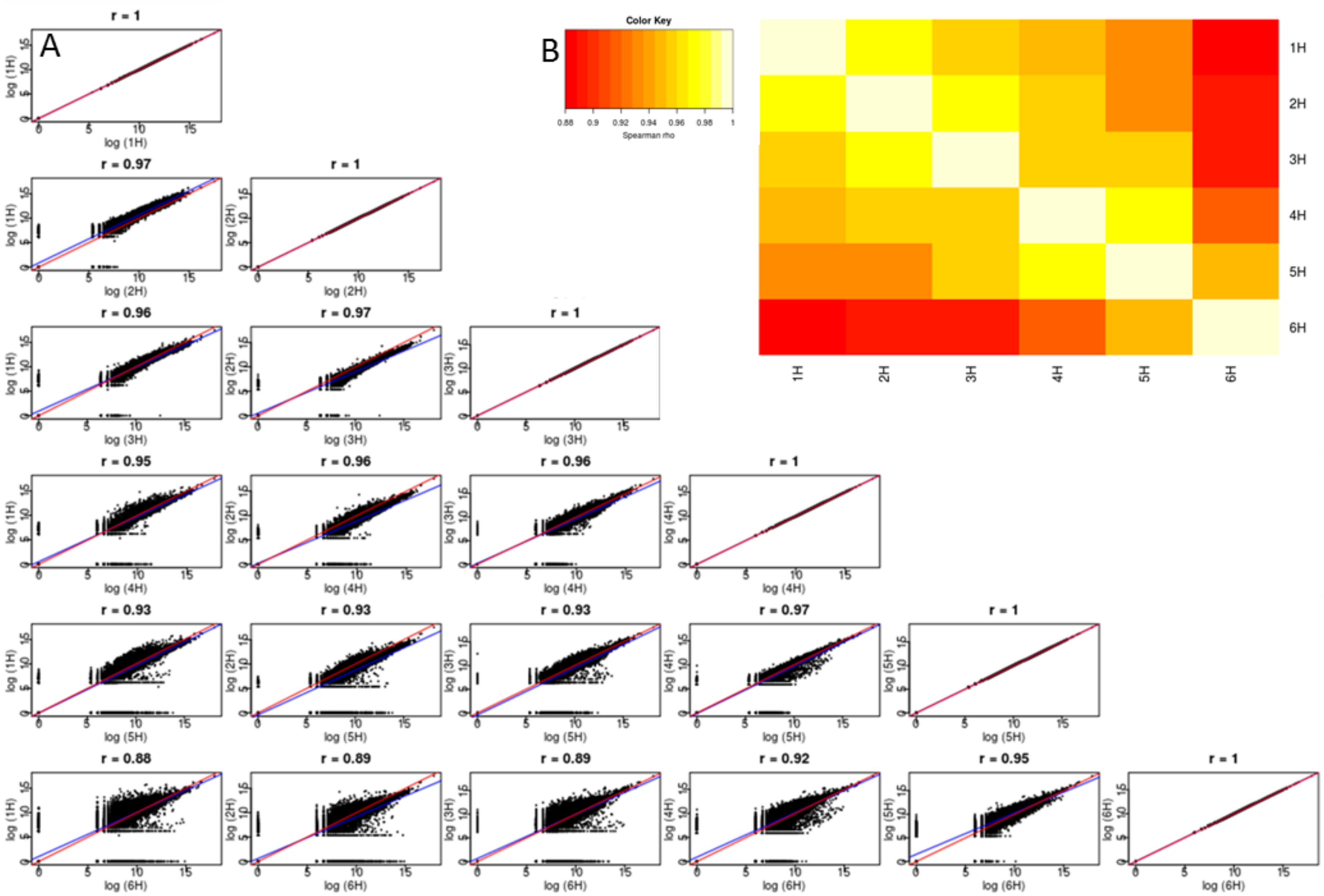

**Supplementary Figure 5.** Correlation of gene expression across time-points. A) The gene expression of each timepoint was compared to all other timepoints and the Spearman rho (r) correlation determined. The plots are fitted with linear model (blue) and arbitrary line with intercept set at 0 and slope of 1 (red). B) Heatmap of the Spearman correlations from (A).

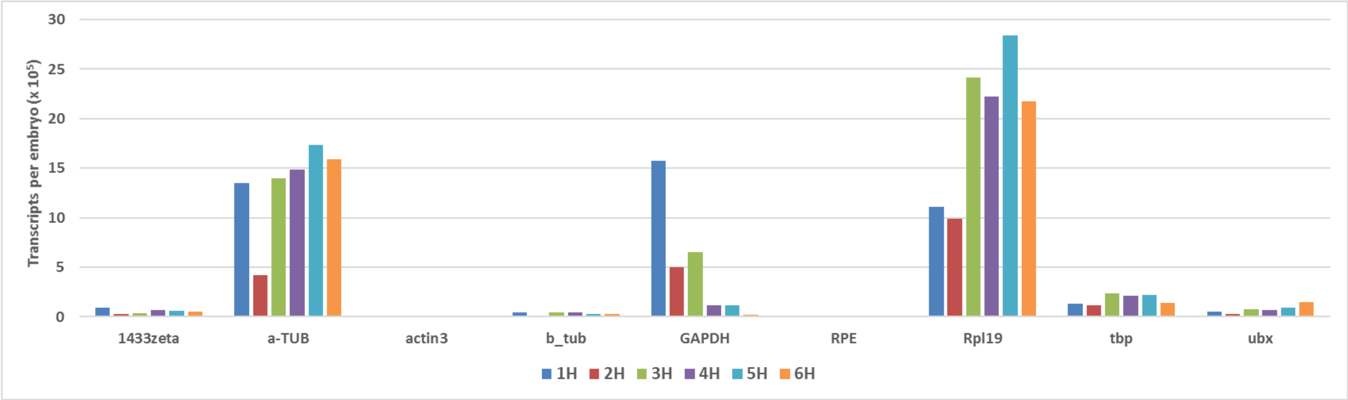


**Supplementary Figure 6.** Comparison of absolute gene expression patterns of 9 genes routinely used as reference genes in qPCR normalization [27].

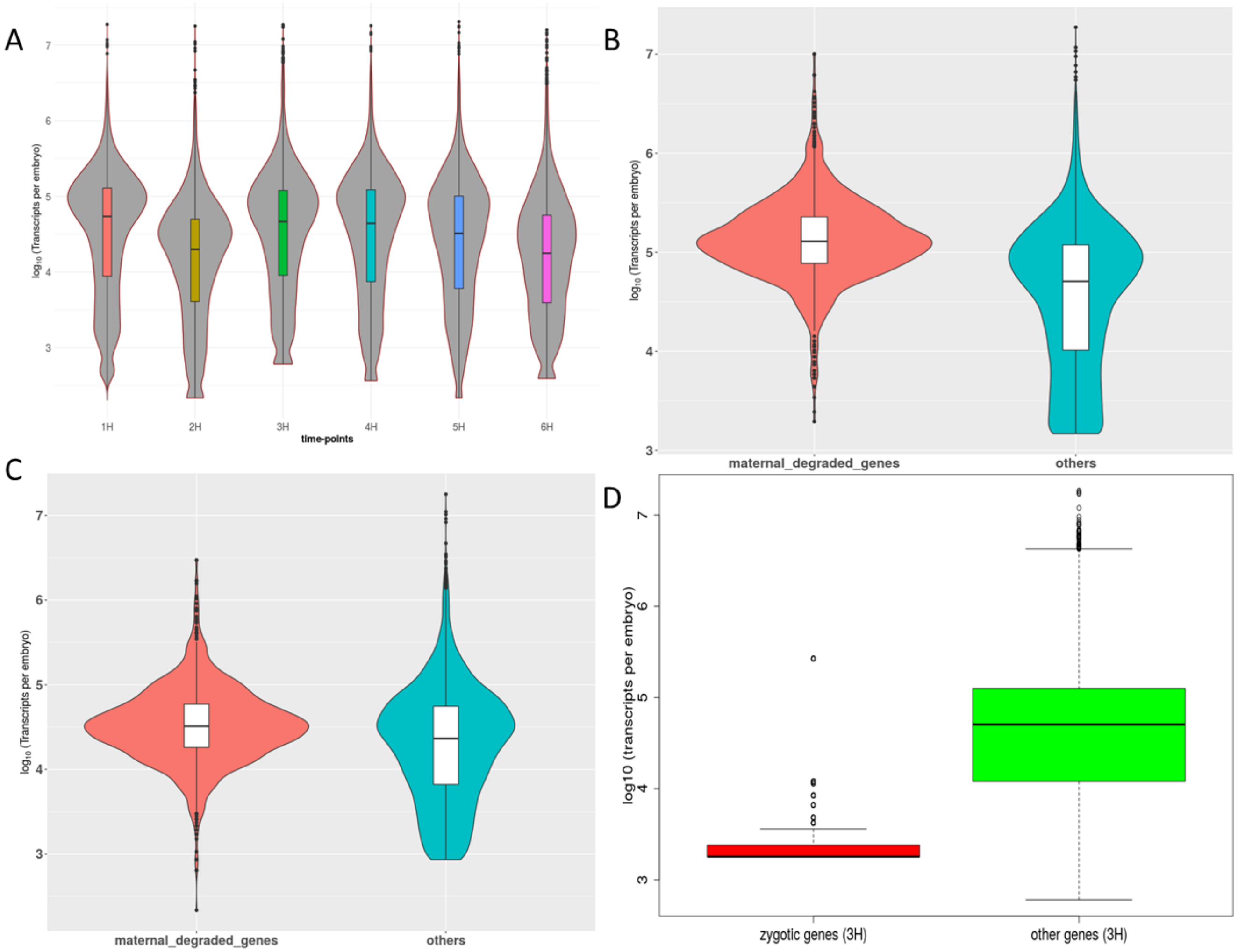


**Supplementary Figure 7.** Reorganization of maternal to zygotic transcripts. A) Violin plot showing the variation in gene expression across timepoints. B) Violin plot of comparing gene expression between maternal-degraded genes and the rest of the genes at 1 hpo. C) Same as B but for 3 hpo. D) Boxplot comparing gene expression pattern of zygotic and all the other genes at 3 hpo. Maternal-degraded genes are defined as genes with a Gfold >0.5 between 1 hour post oviposition (hpo) and 2 hpo. Zygotic genes are genes whose expression is not detected at 1 hpo but detected thereafter suggesting they were transcribed from the zygotic genome as opposed to being maternally derived.

#### Extended material

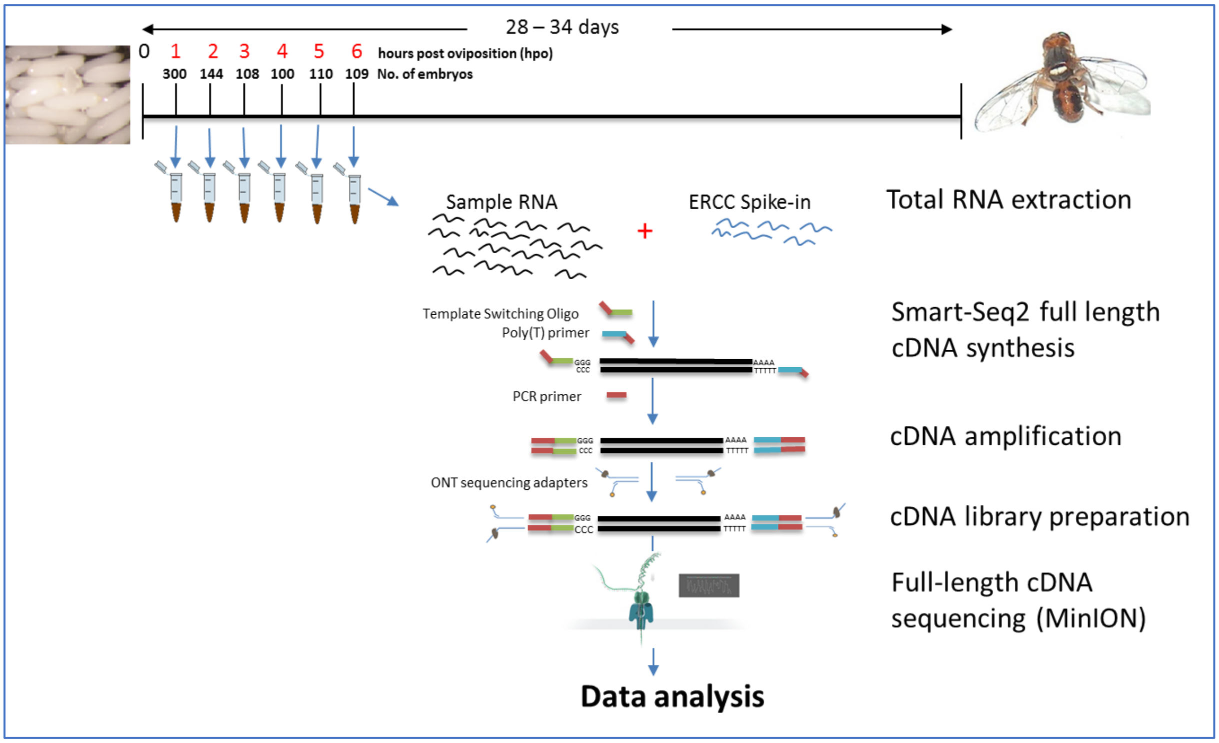


**Extended Figure 1.** Schematic of cDNA library generation and sequencing. Embryos were collected at hourly point post oviposition (hpo), counted and total RNA extracted using the Trizol method. At cDNA synthesis step, external RNA standards (ERCC) were added to each sample commensurate to the number of embryos that were used. The Smart-Seq2 protocol was used to generate full length cDNA, followed by PCR amplification of the cDNA. The Oxford Nanopore Technologies (ONT) SQK-LSK108 protocol for library preparation was then followed, albeit with some custom changes. The library was then sequenced on the ONT MinION, followed by basecalling using ONT Albacore basecaller.

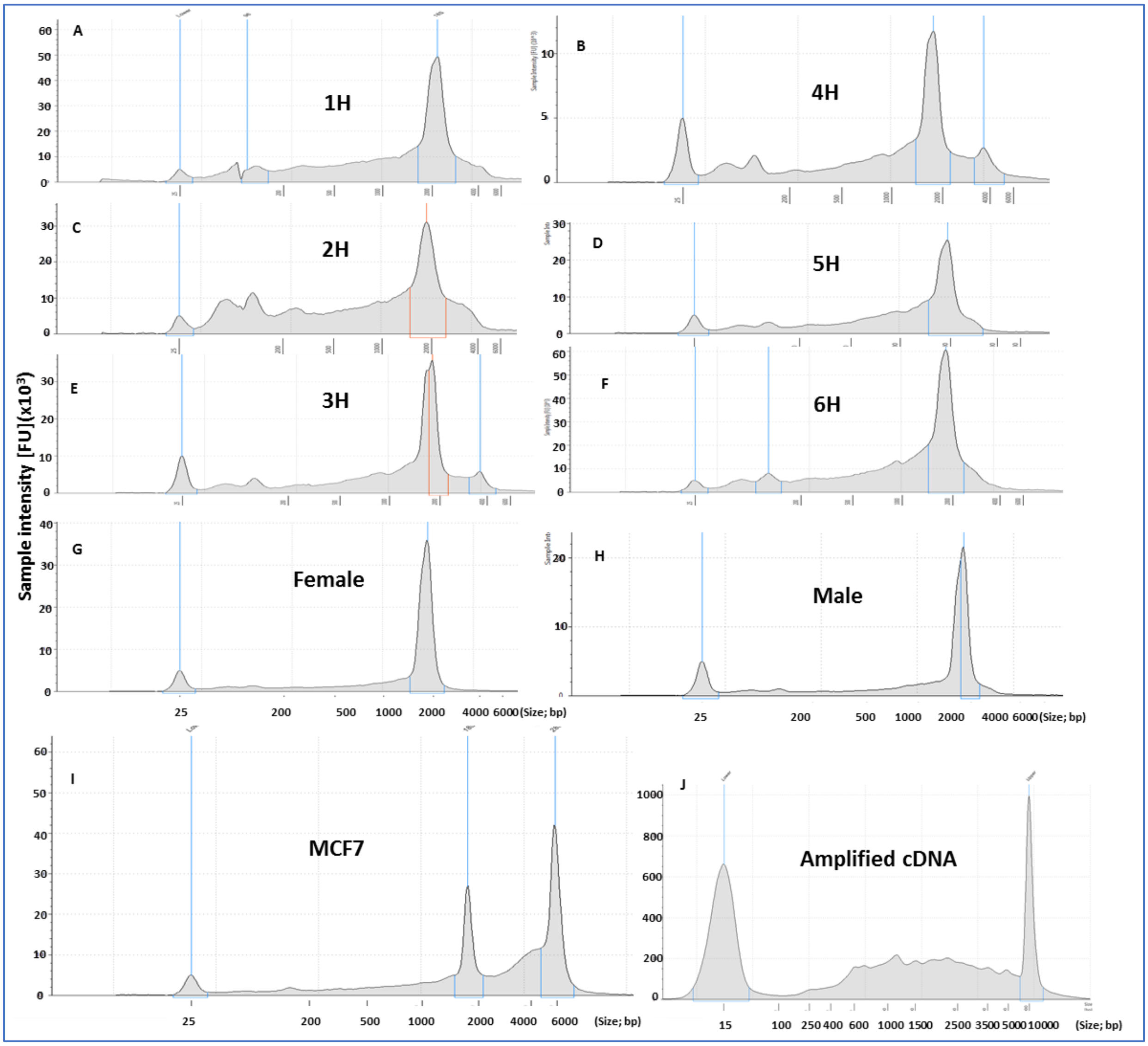


**Extended Figure 2.** Electropherogram showing the profile of total RNA of the samples used in this experiment. A-F) Show the profile of embryo samples at the indicated timepoints post oviposition. G-H) Show profile of mature insects. I) MCF-7 total RNA is added to show the difference in profile between mammalian and *B. oleae.* J) Example profile of amplified cDNA generated from the samples.

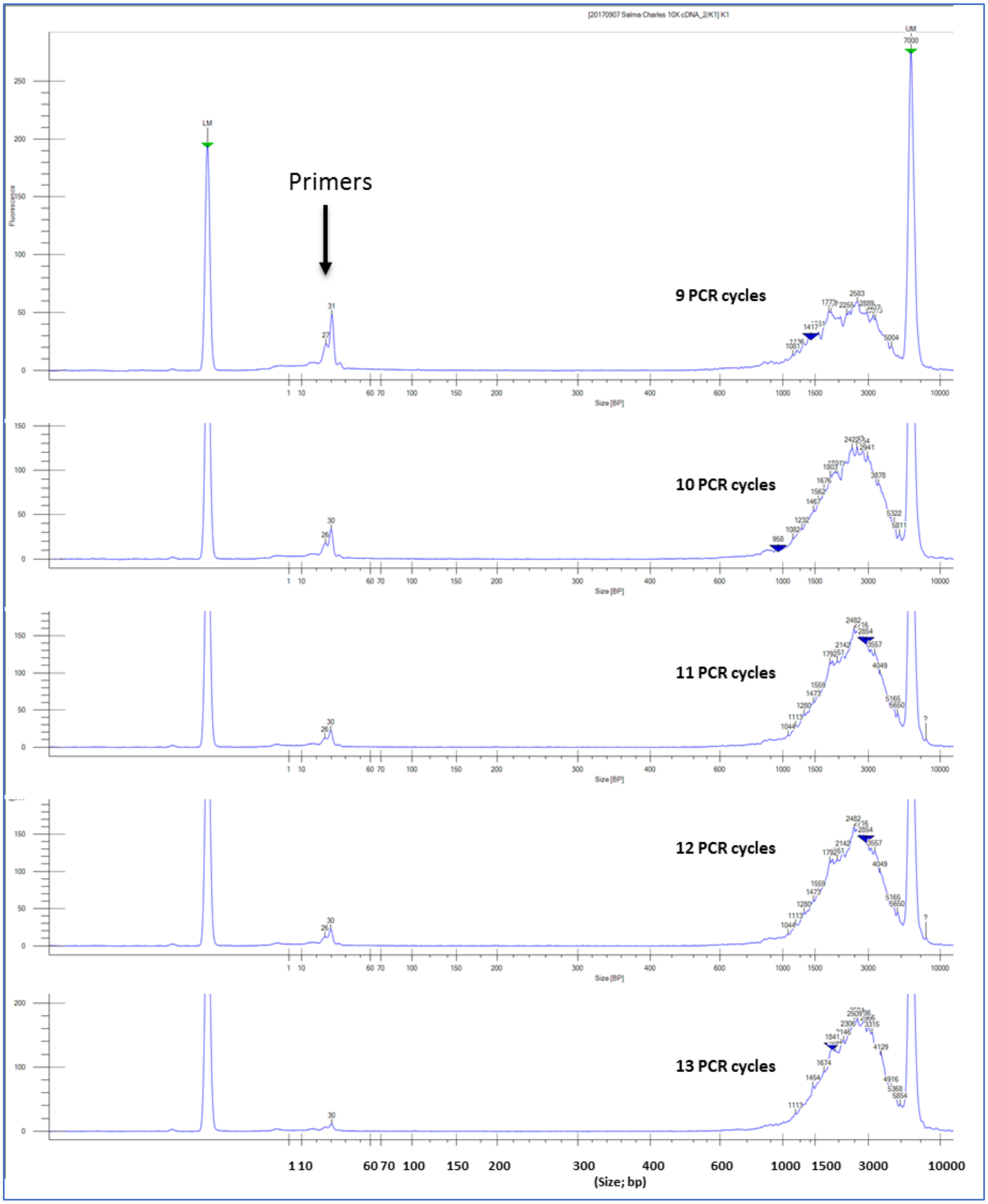


**Extended Figure 3.** Optimization of the number of cycles needed to amplify the cDNA in order to obtain enough material without skewing the distribution. We selected 12 cycles which gave enough material (~2 µg) without dramatically skewing cDNA profile.

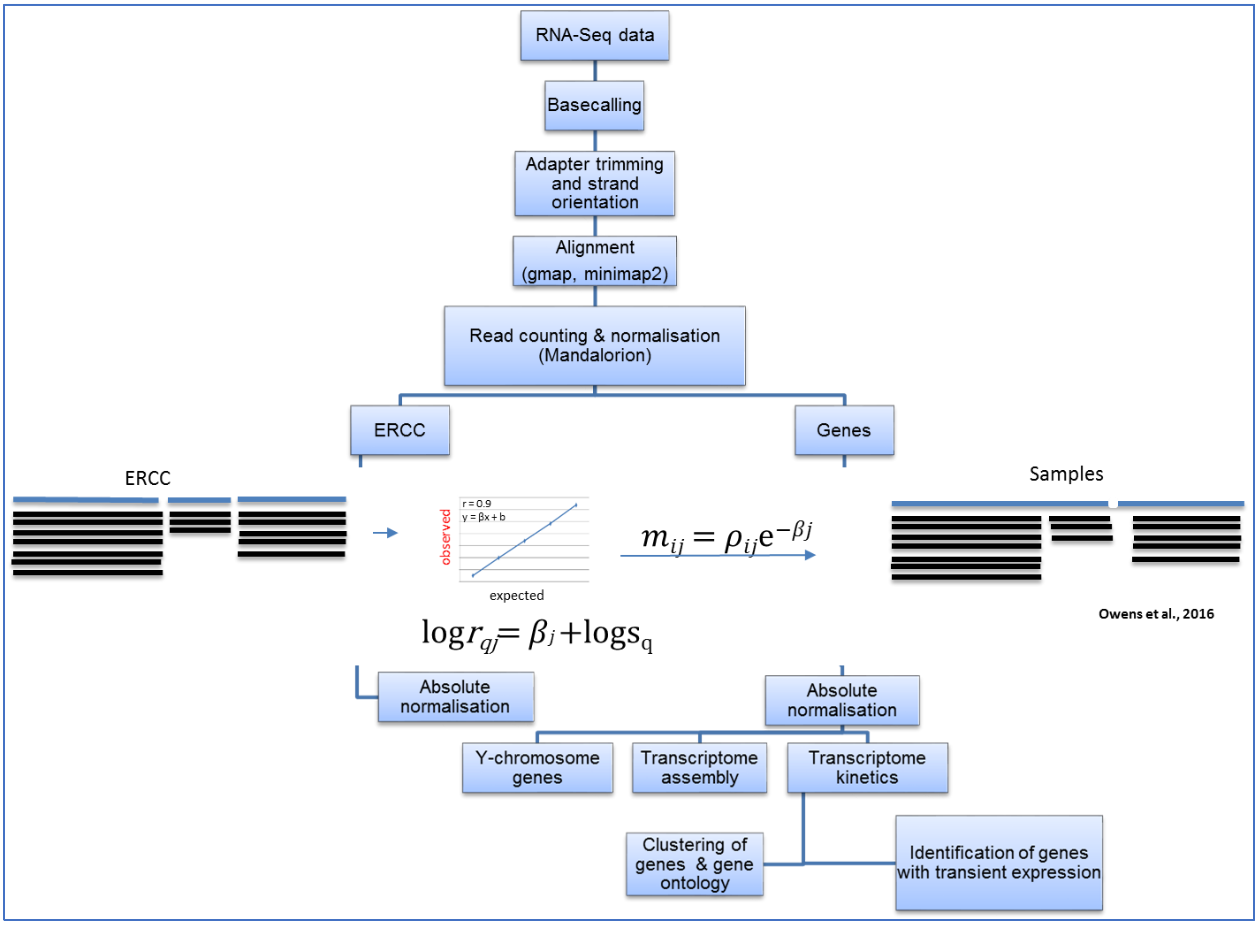


**Extended Figure 4.** Data analysis workflow. Following sequencing, reads were basecalled using Albacore (ONT, version 2.0.2). A customized version of Mandalorion was used to perform adapter trimming and strand orientation. Trimmed and stranded reads were aligned to the genome using GMAP followed by relative quantification of expression using Mandalorion. Relative expression were converted to absolute quantification using ERCC standards, followed by downstream data analyses

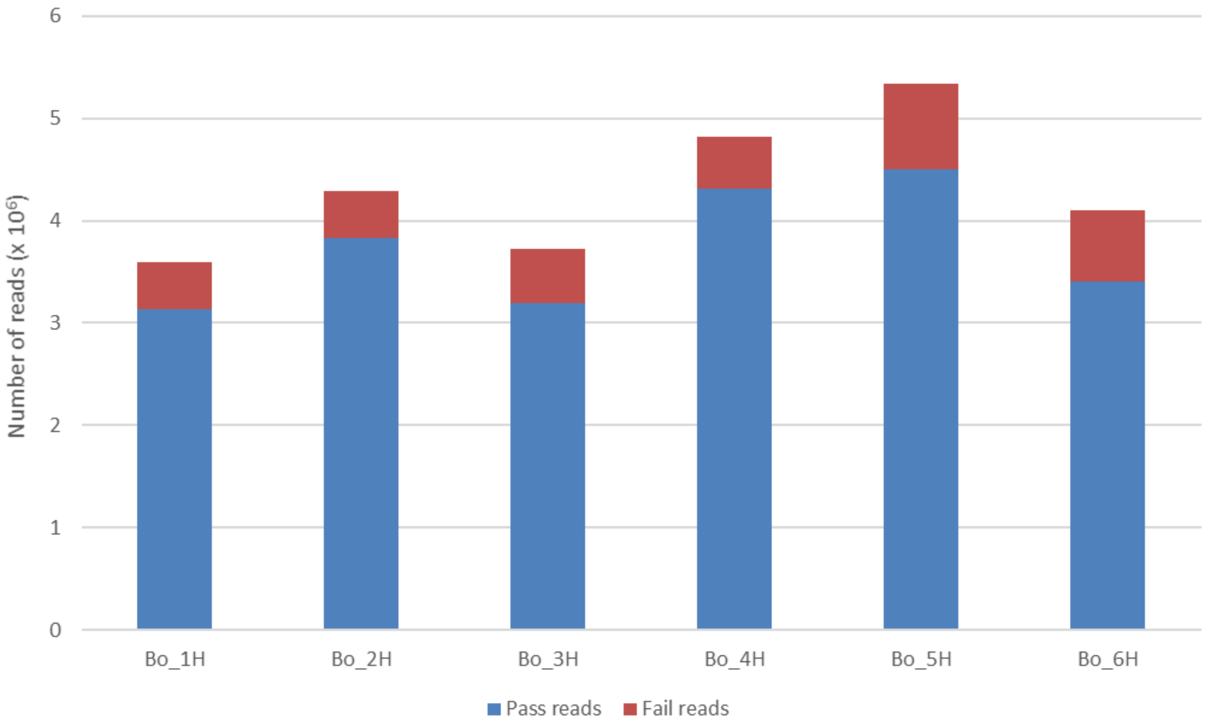


**Extended Figure 5. Summary read stats from ONT sequencing of *B. oleae* embryo samples**

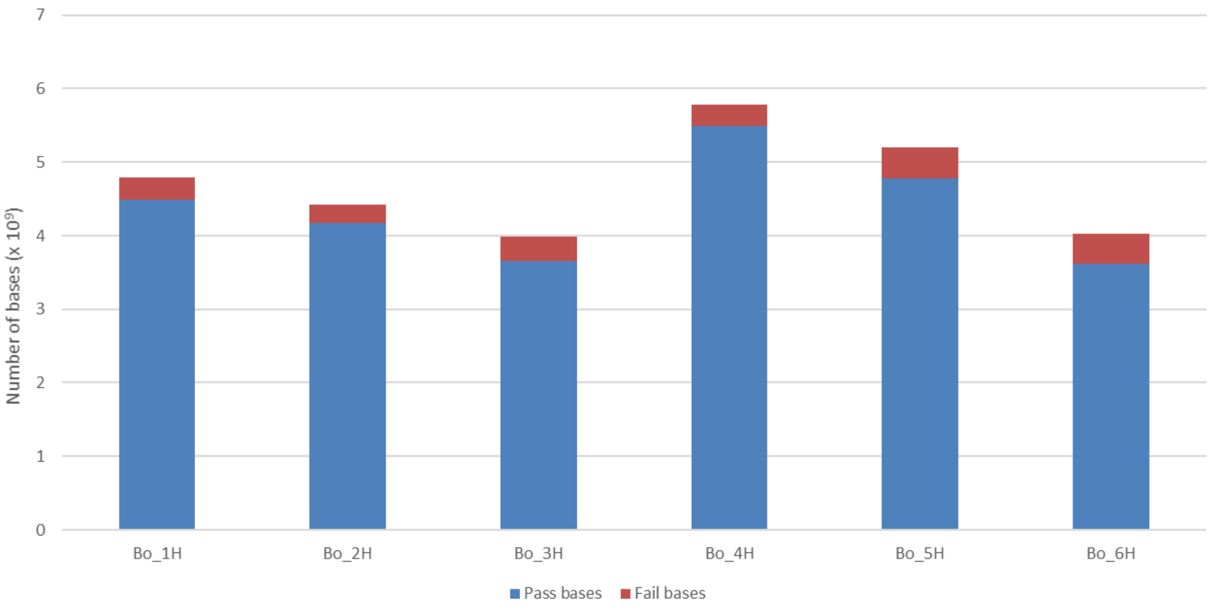


**Extended Figure 6.** Summary bases stats from ONT sequencing of *B. oleae* embryo samples

**Extended Figure 7: Quality control profiles of Oxford Nanopore sequencing runs**

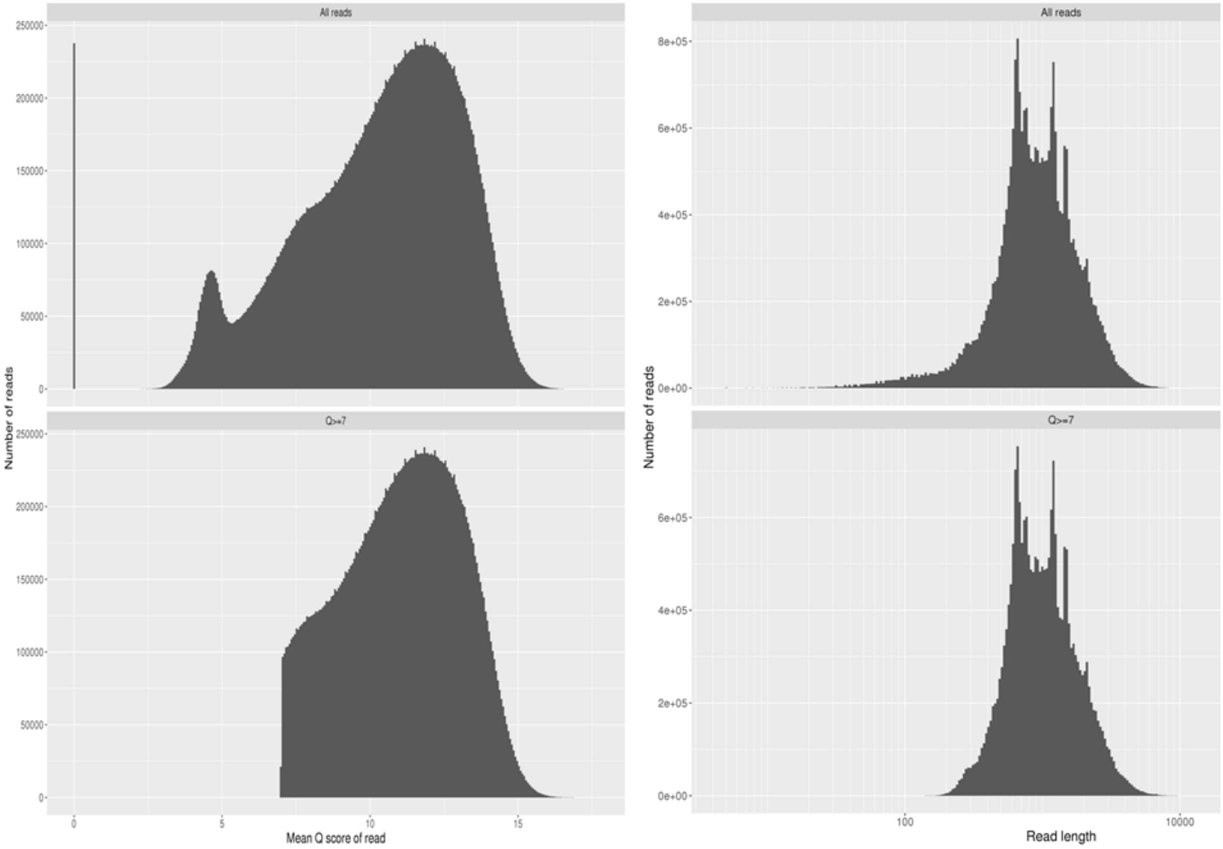


**Extended Figure 7 A**

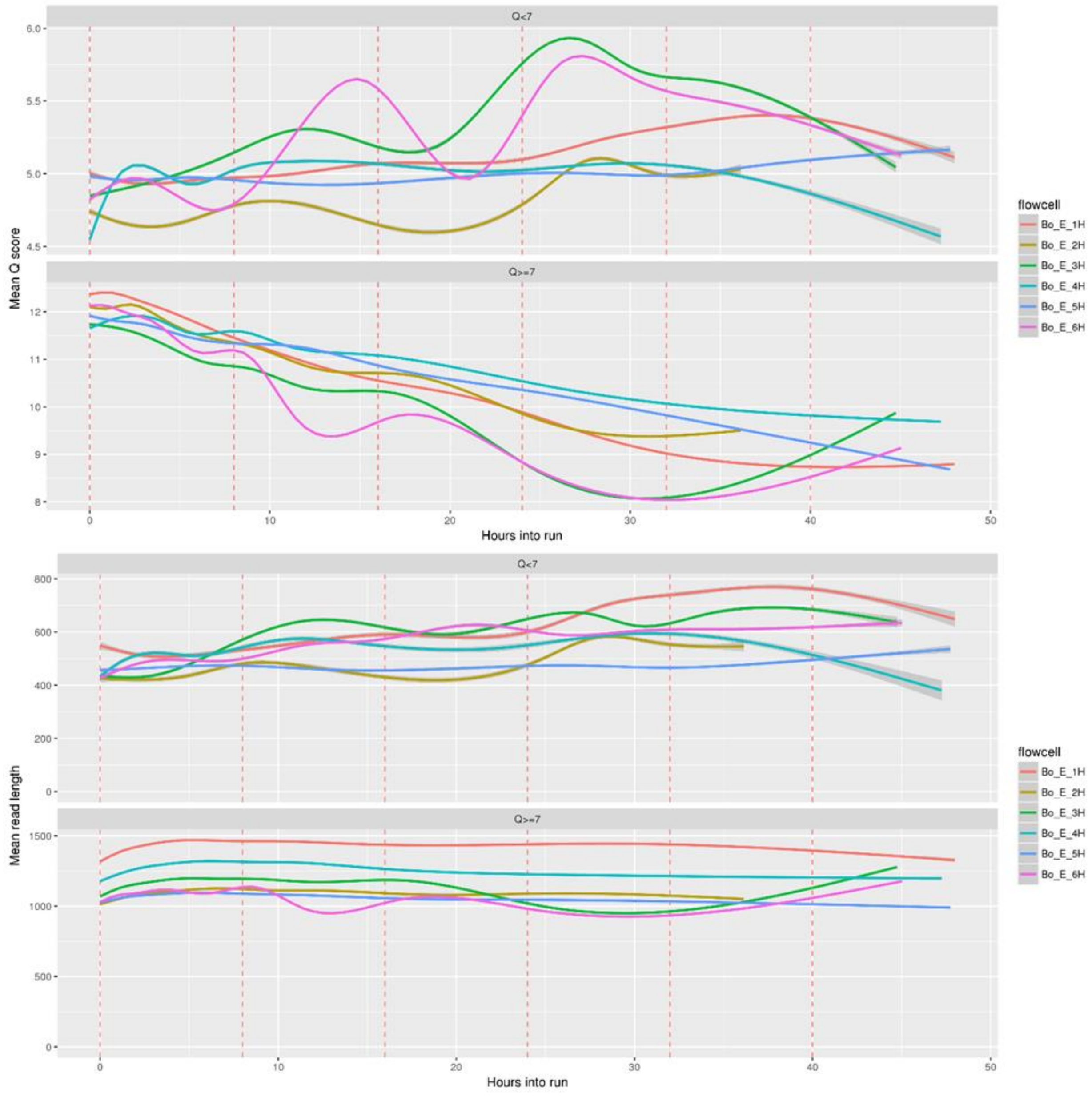


**Extended Figure 7 B**

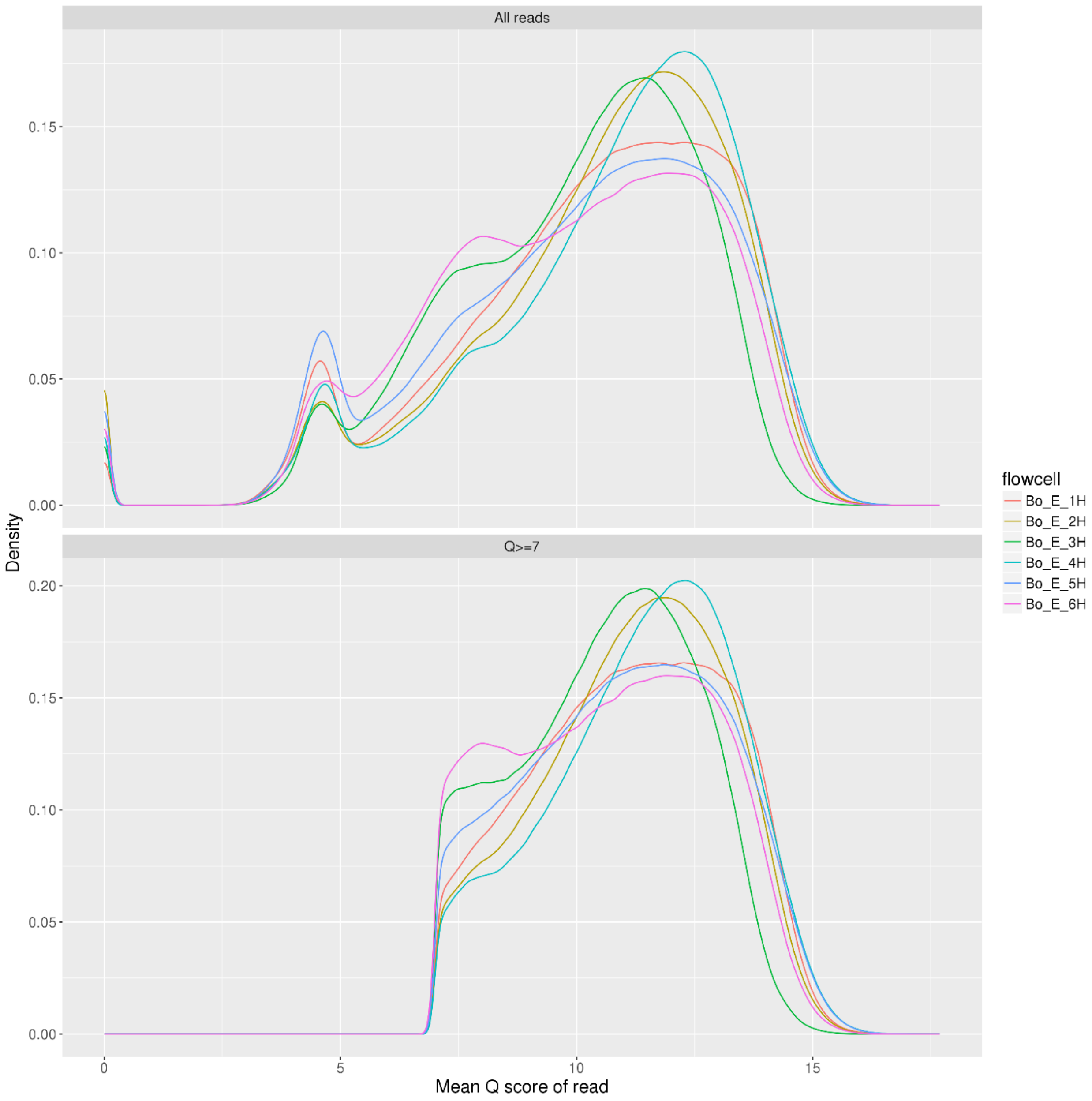


**Extended Figure 7 C**

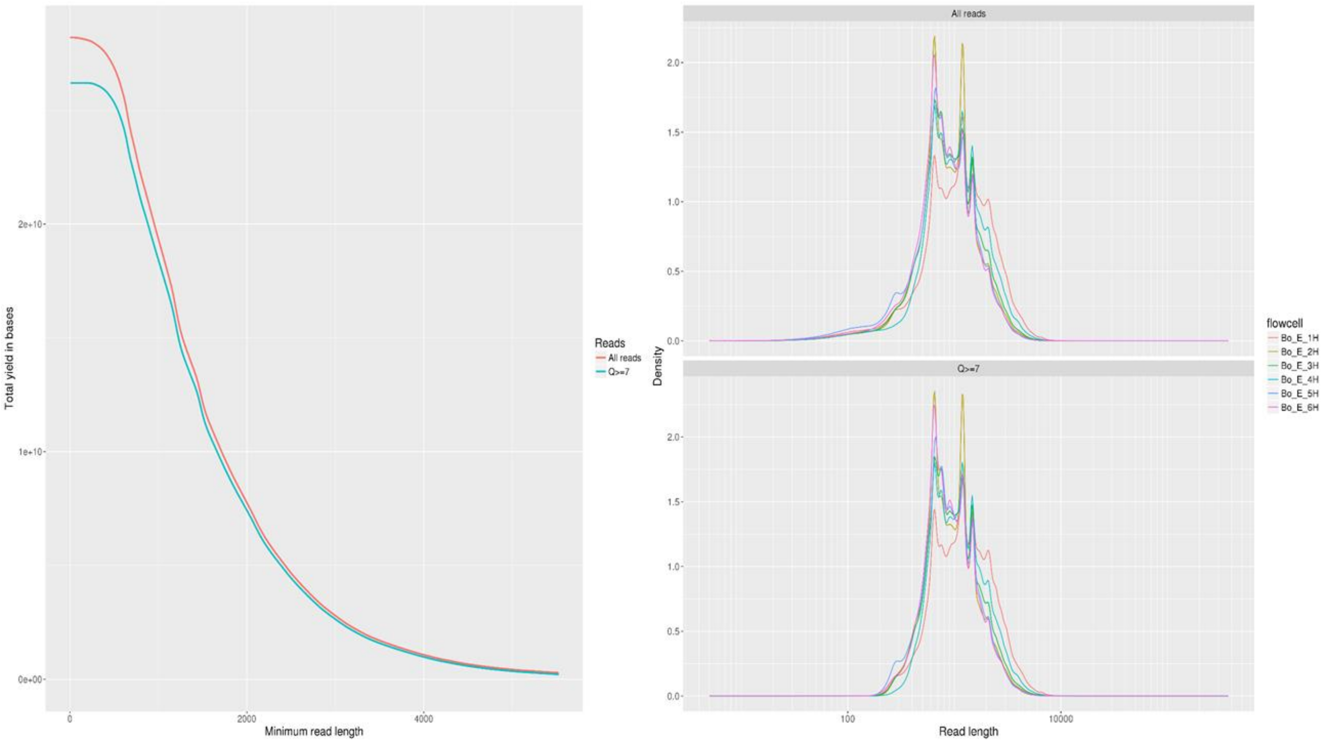


**Extended Figure 7 D**

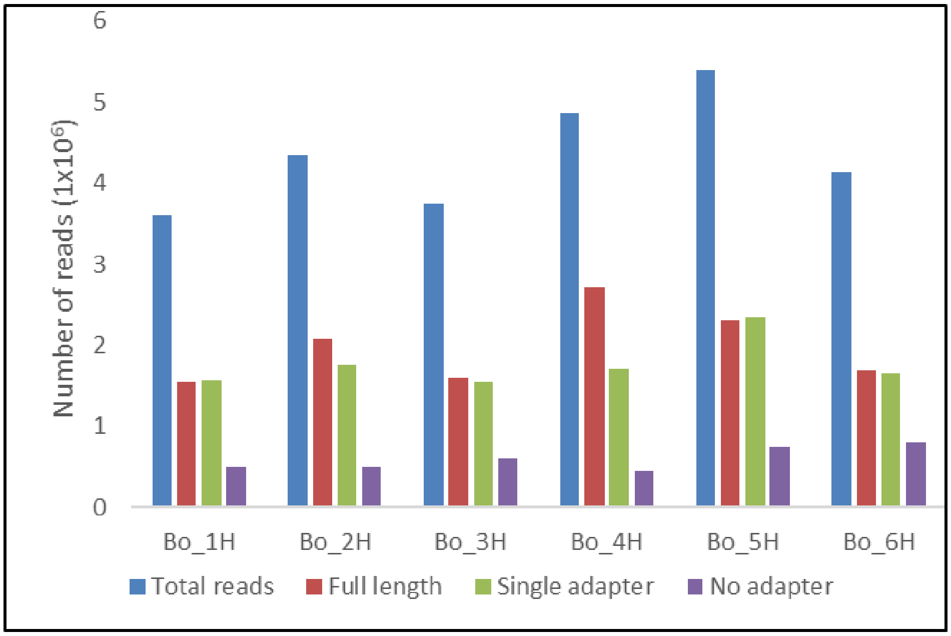


**Extended Figure 8.** Number of reads identified as full-length according to detection of 5' and 3' adapters and those with a single adapter or no adapter at all.

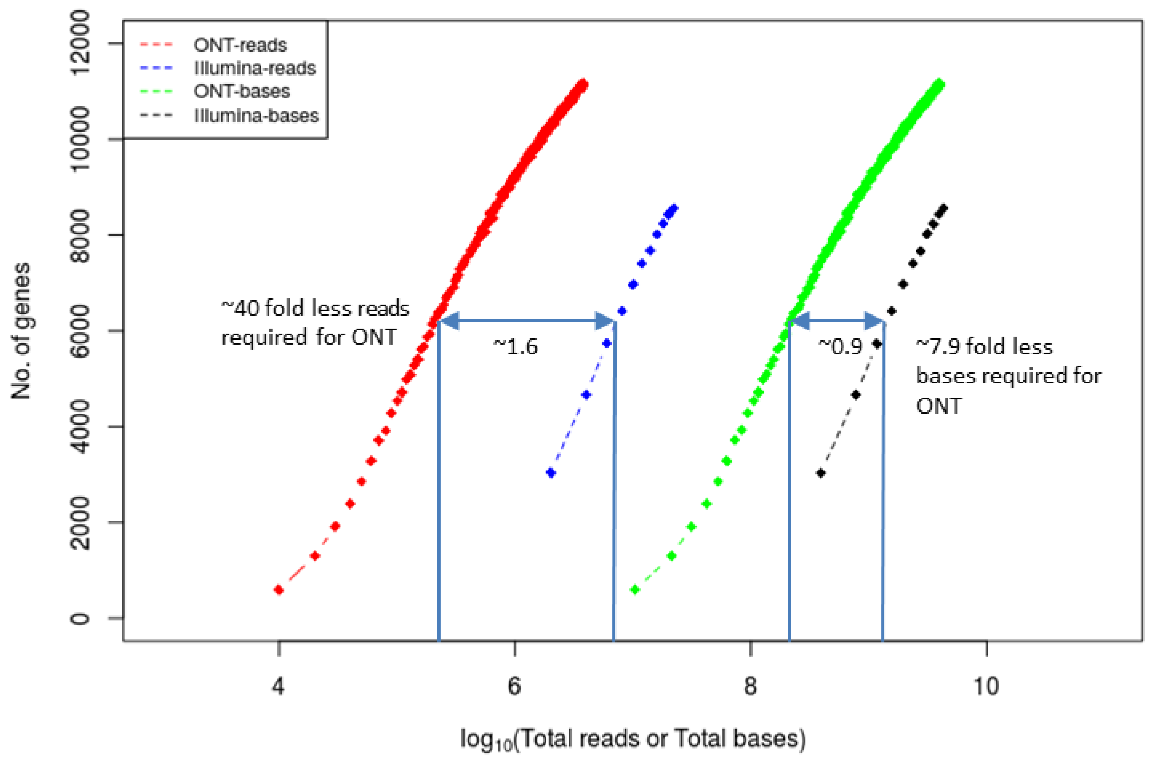


**Extended Figure 9. Rare fraction curve comparing number of reads and bases required to observe the same number of genes between long-reads (Oxford Nanopore) and short-reads (Illumina).**

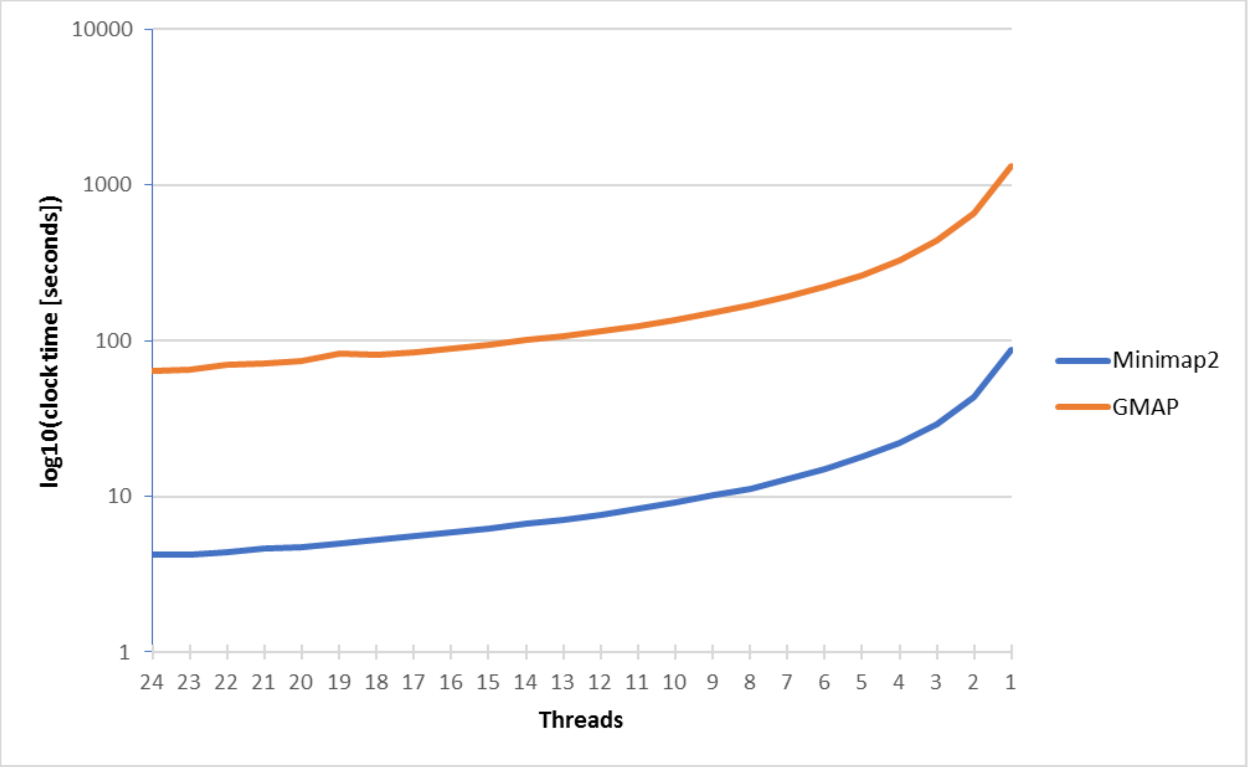


**Extended Figure 10.** Comparison of computational efficiency of GMAP and Minimap2. One million Nanopore long cDNA reads were aligned to the reference genome using GMAP or Minimap2 setting different processors (threads) and the amount of clock-time require to complete the time recorded. Both tools showed similar scaling with number of threads although Minimap2 showed exceptional speed in completing the jobs.

Extended Figure 11. In-depth characterization of B. oleae de novo transcriptome assembly with reference to the Bactrocera oleae Annotation release 100 from NCBI. Statistics derived using SQANTI.

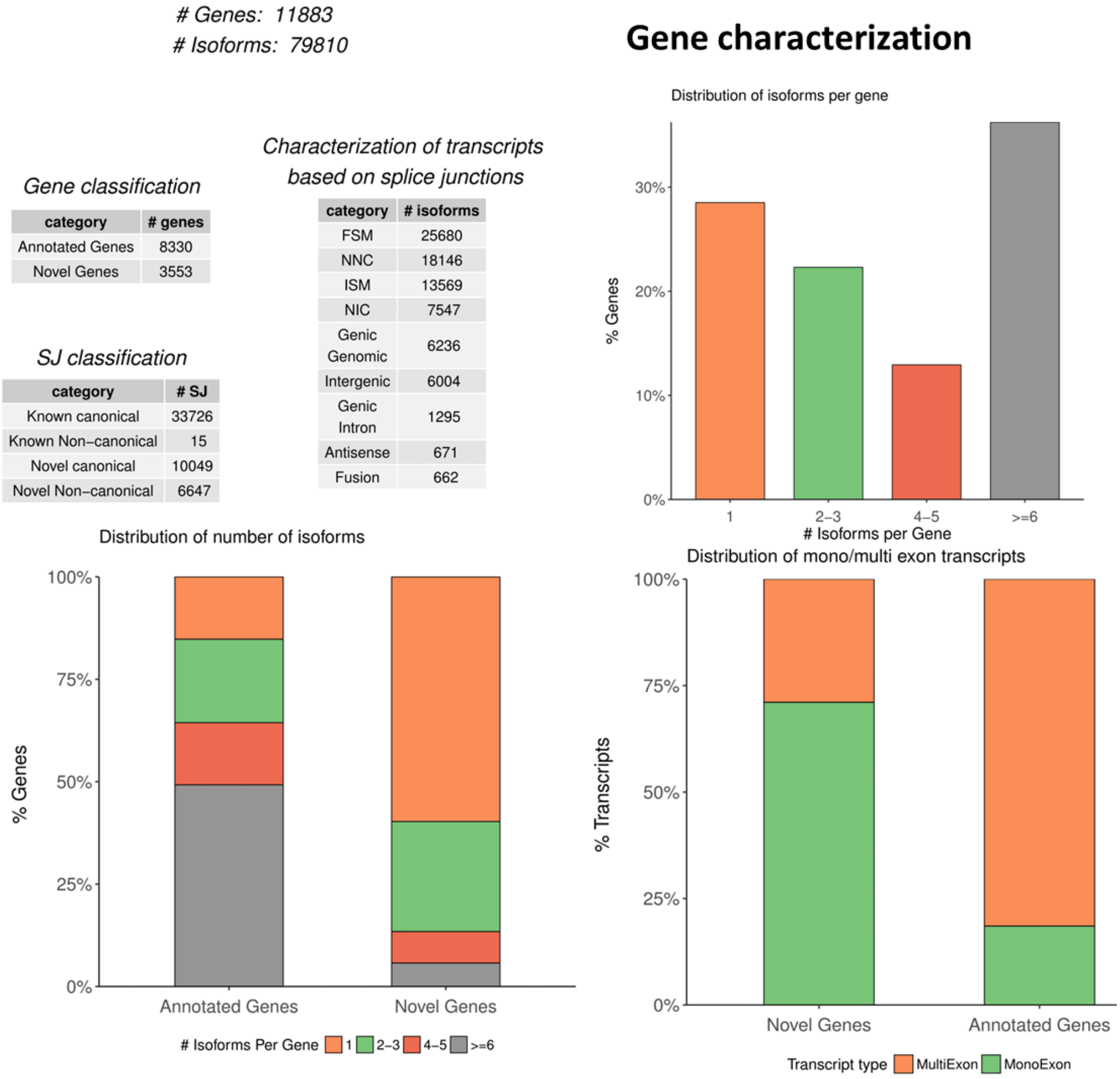


**Extended Figure 11 A**

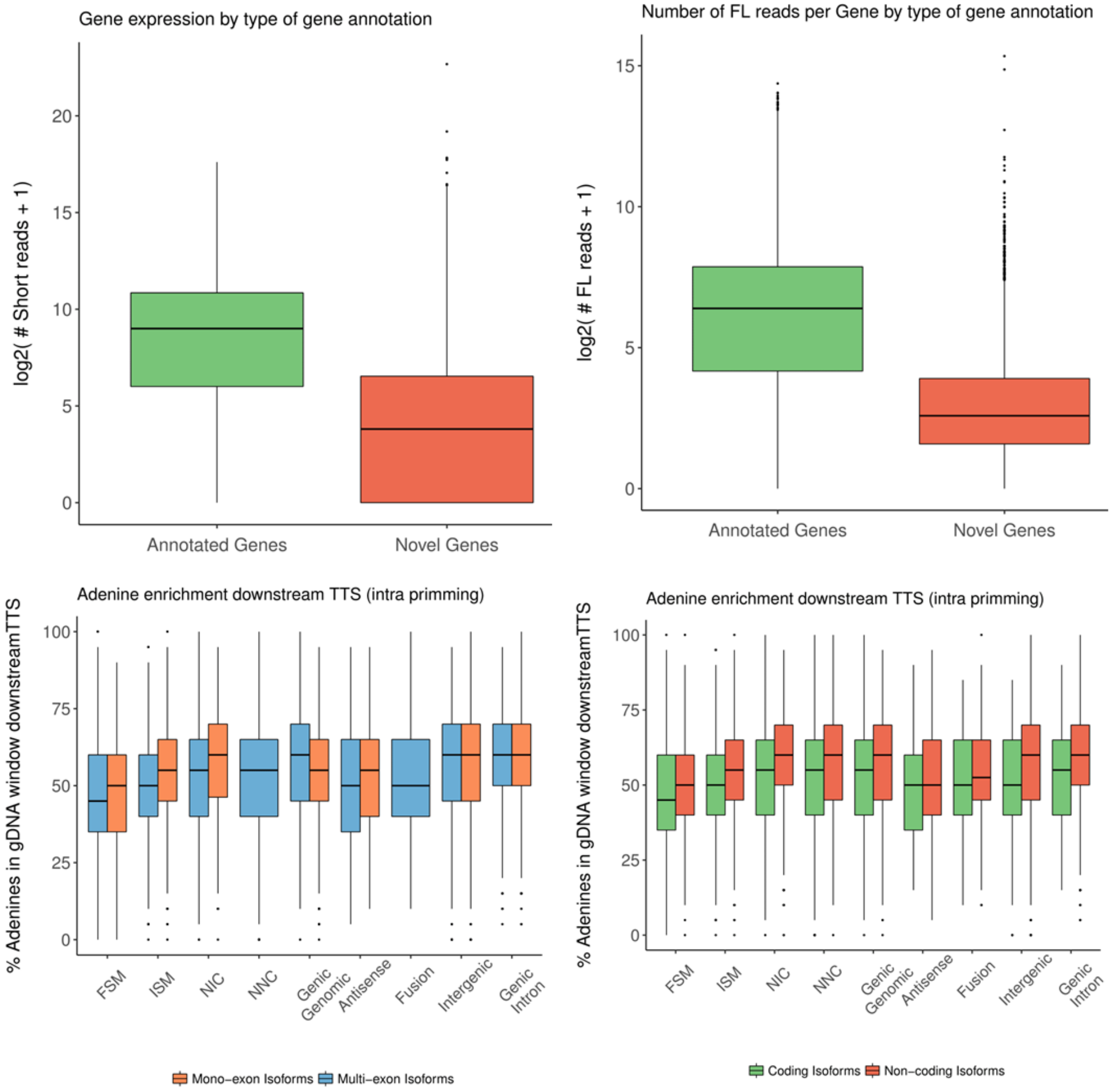


**Extended Figure 11 B**

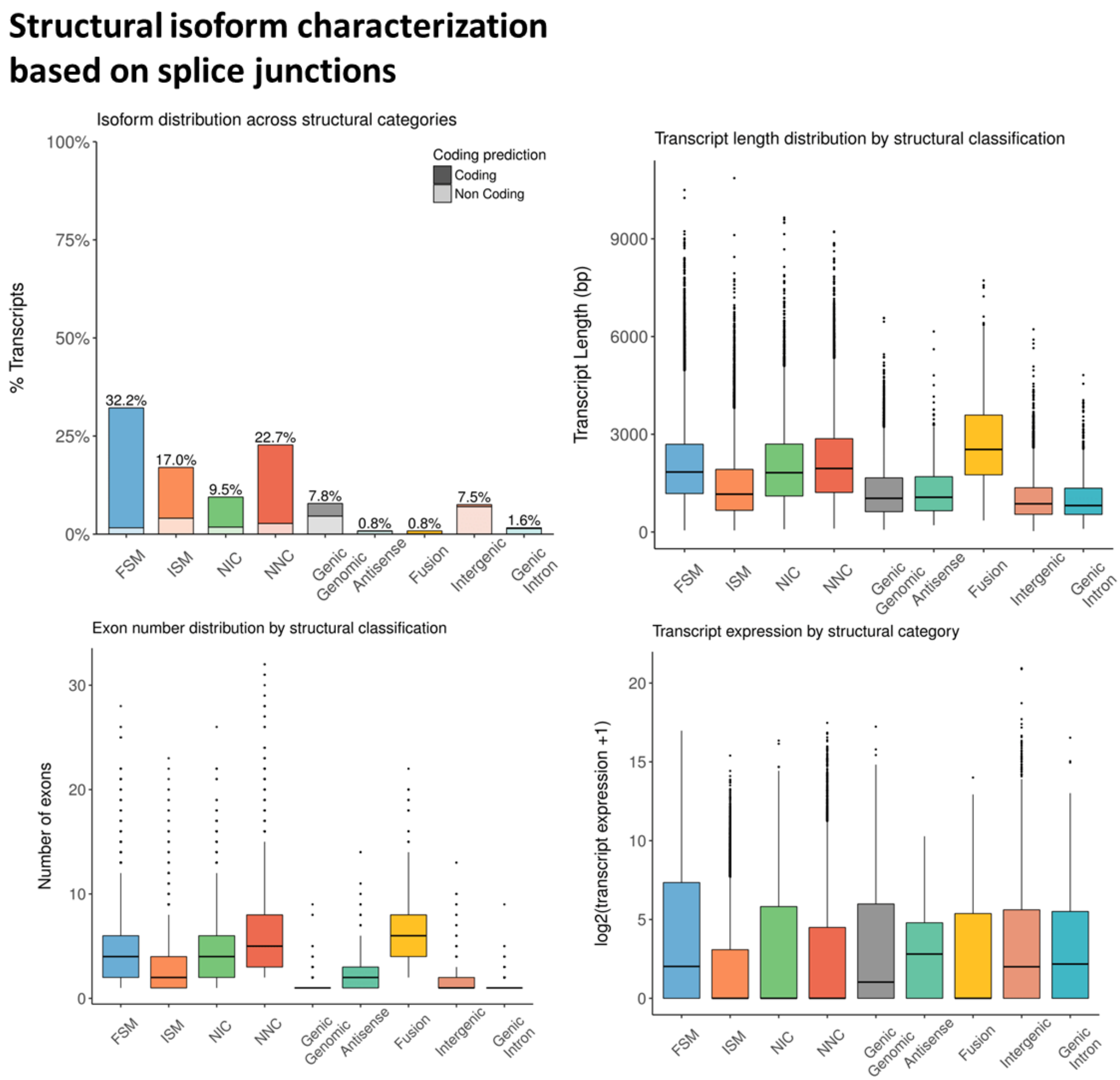


Extended Figure 11 C

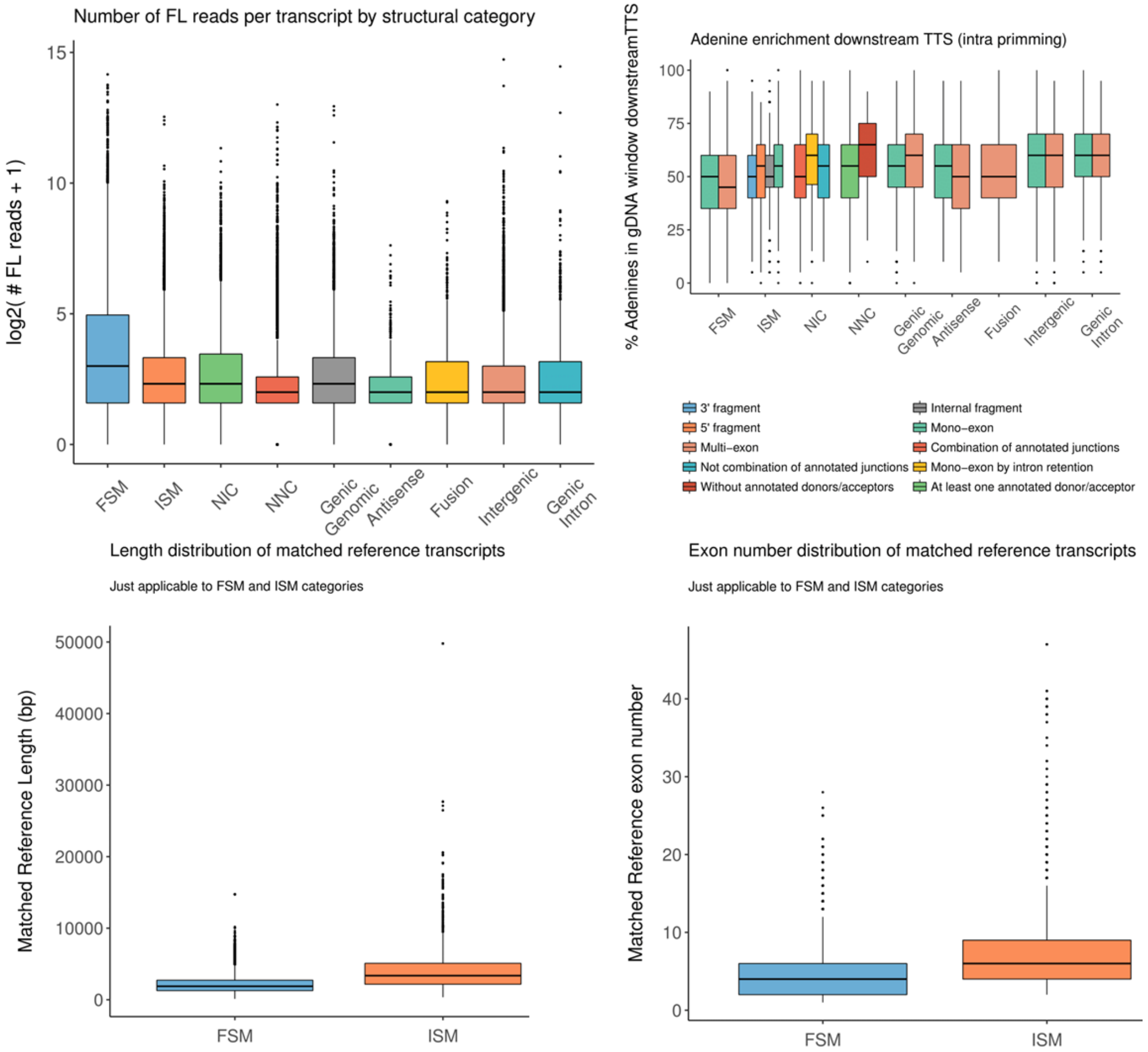


**Extended Figure 11 D**

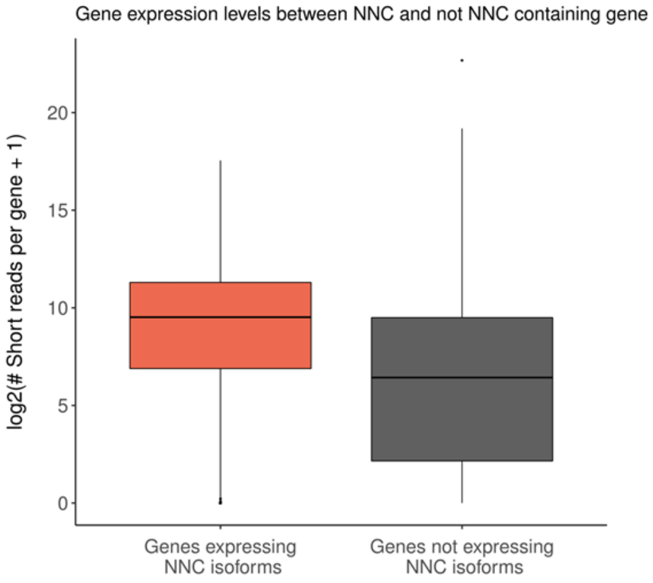


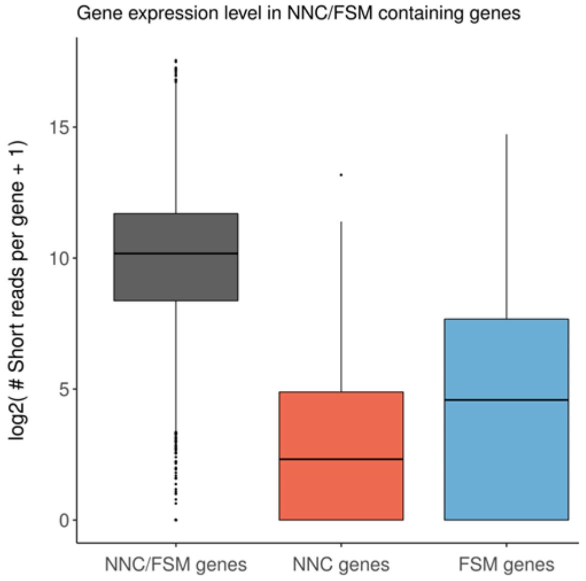


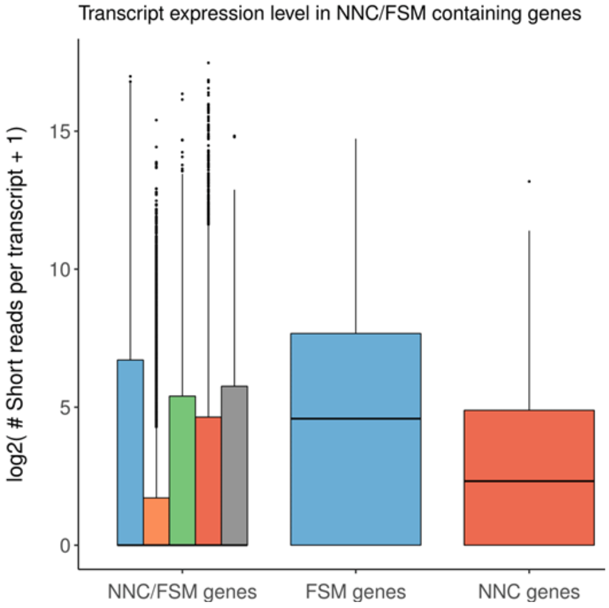


**Extended Figure 11 E**

### Splice junction characterization

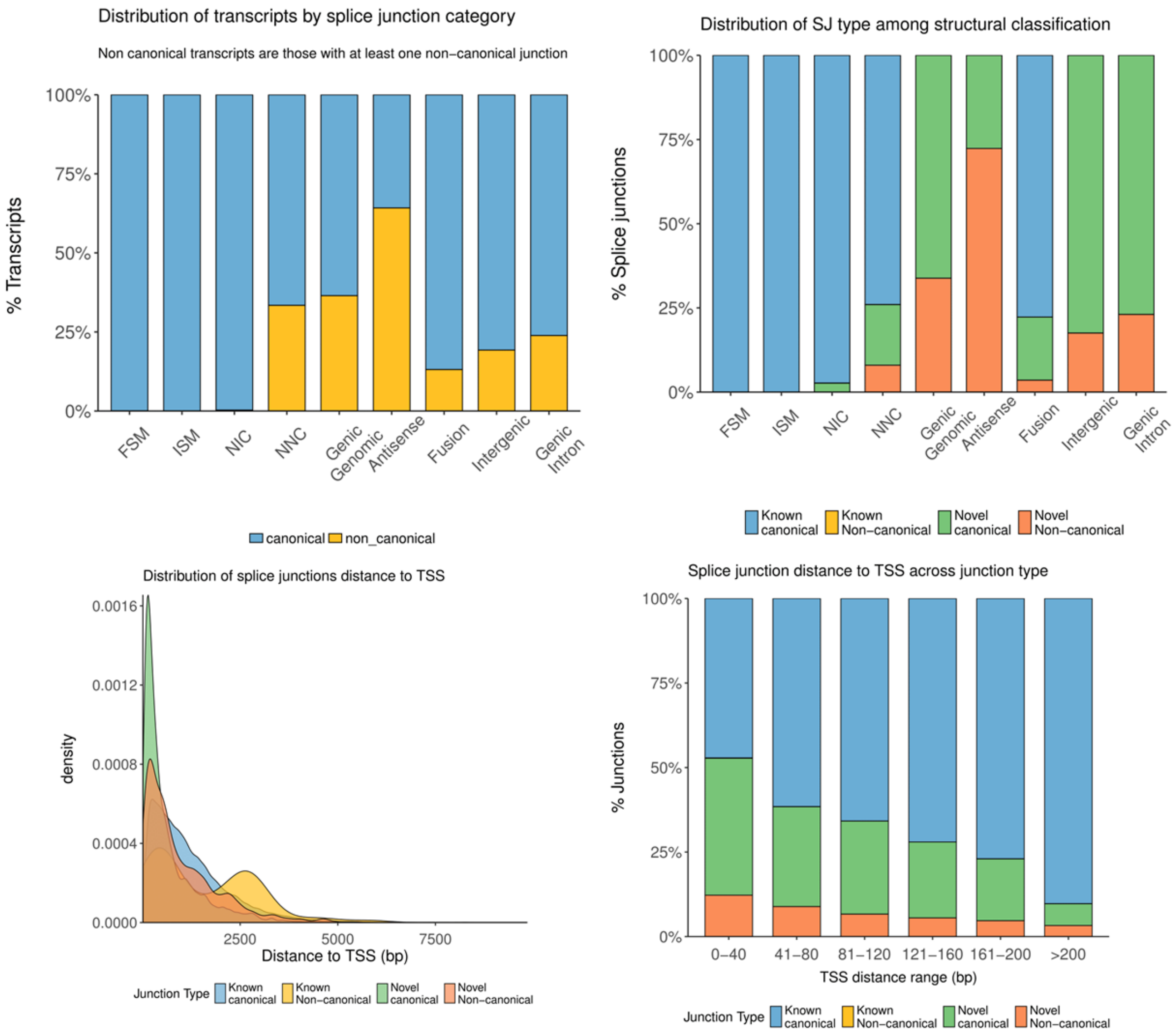


**Extended Figure 11 F**

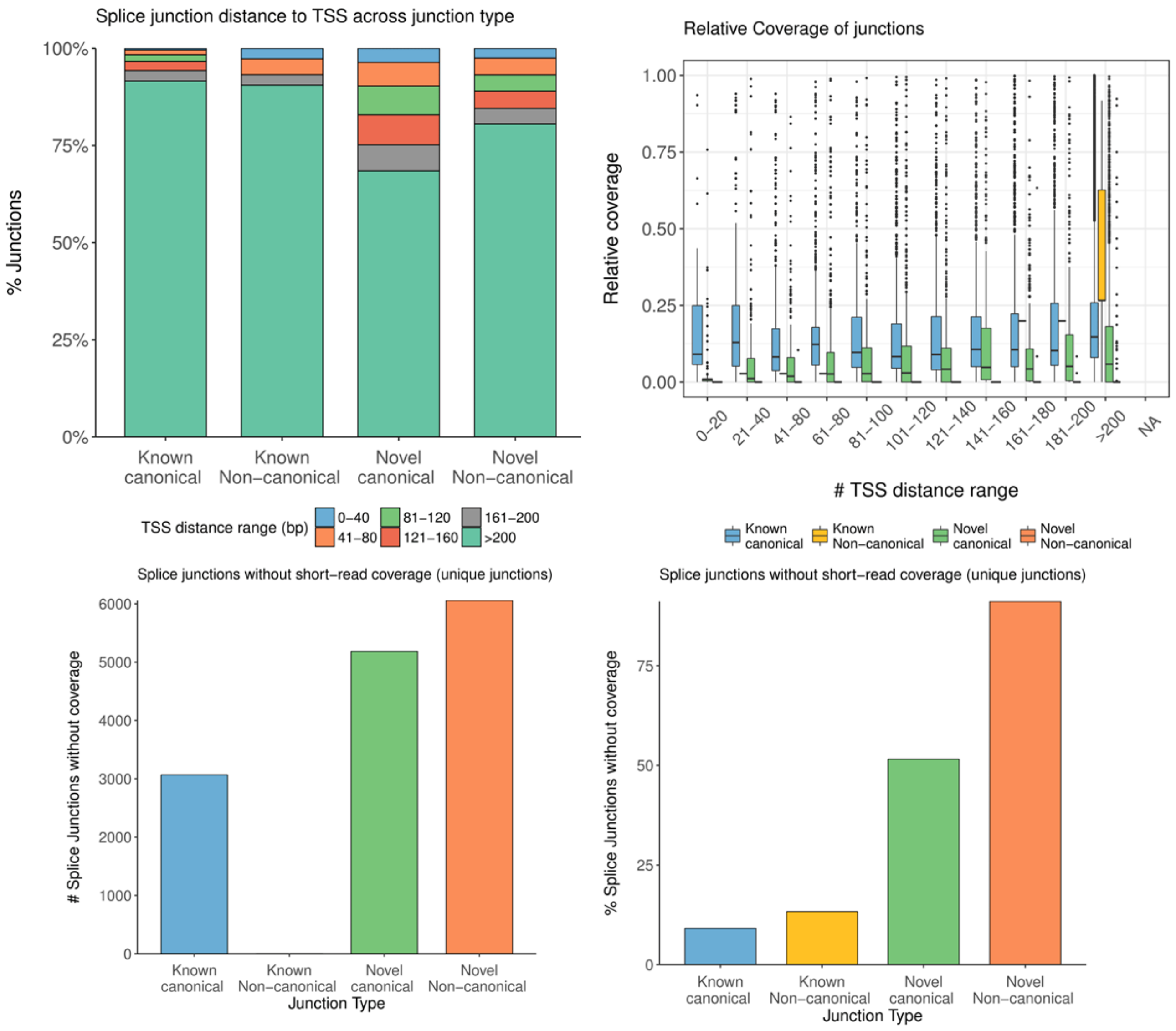


**Extended Figure 11 G**

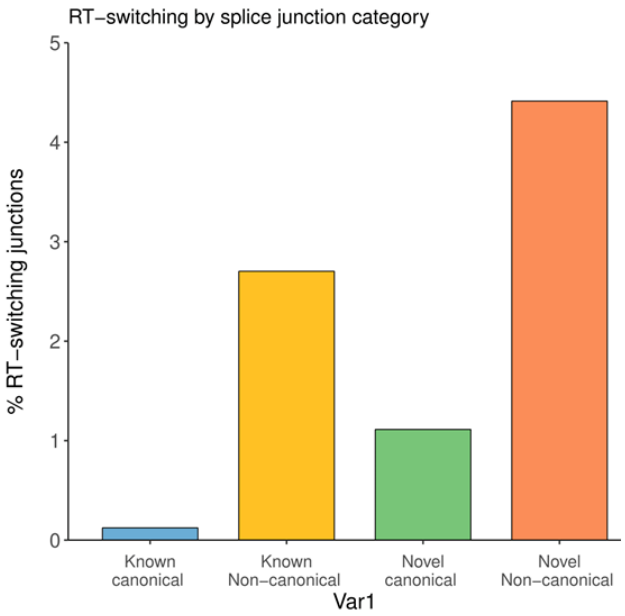


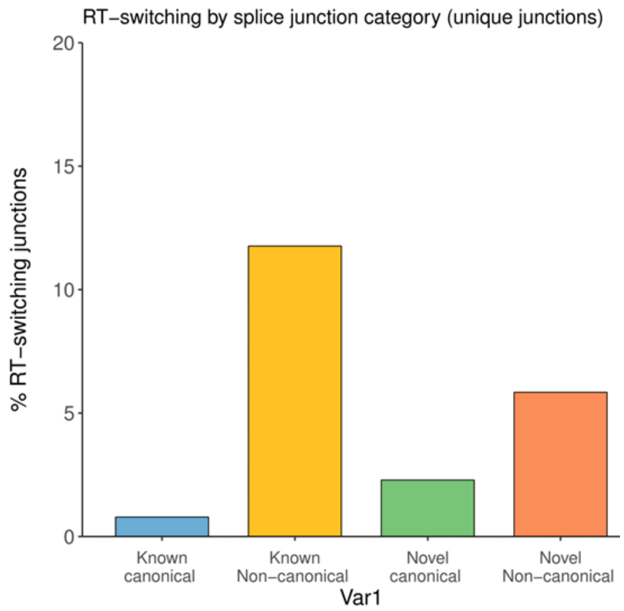


Full-lengthness characterization of isoforms

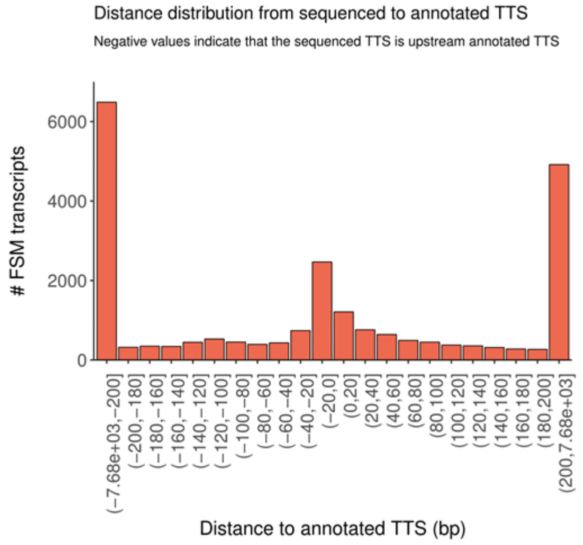


**Extended Figure 11 H**

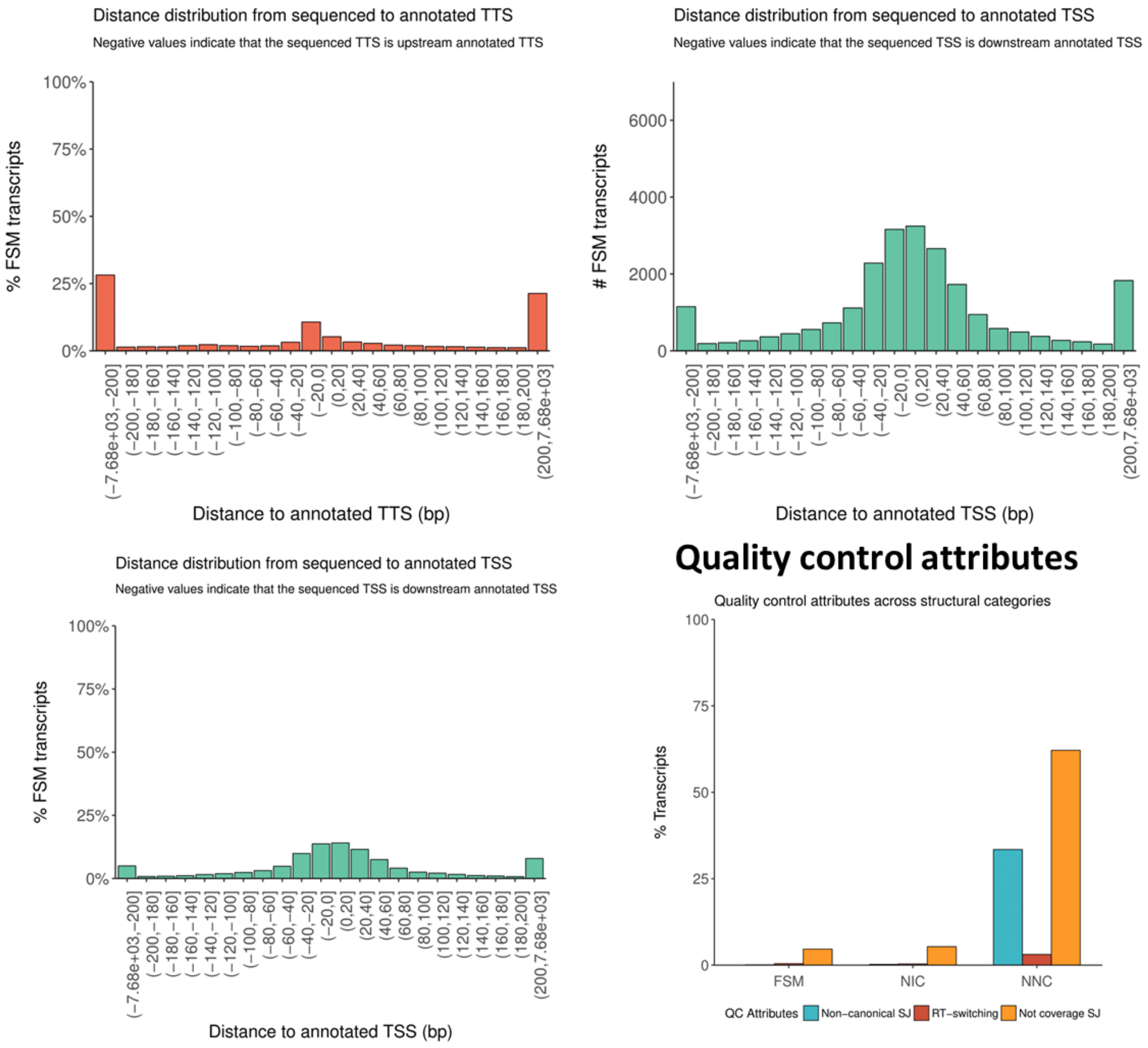


**Extended Figure 11 I**

Extended Figure 12. Alignment statistics of one of the samples. Results were generally similar across samples. Statistics generated using AlignQC

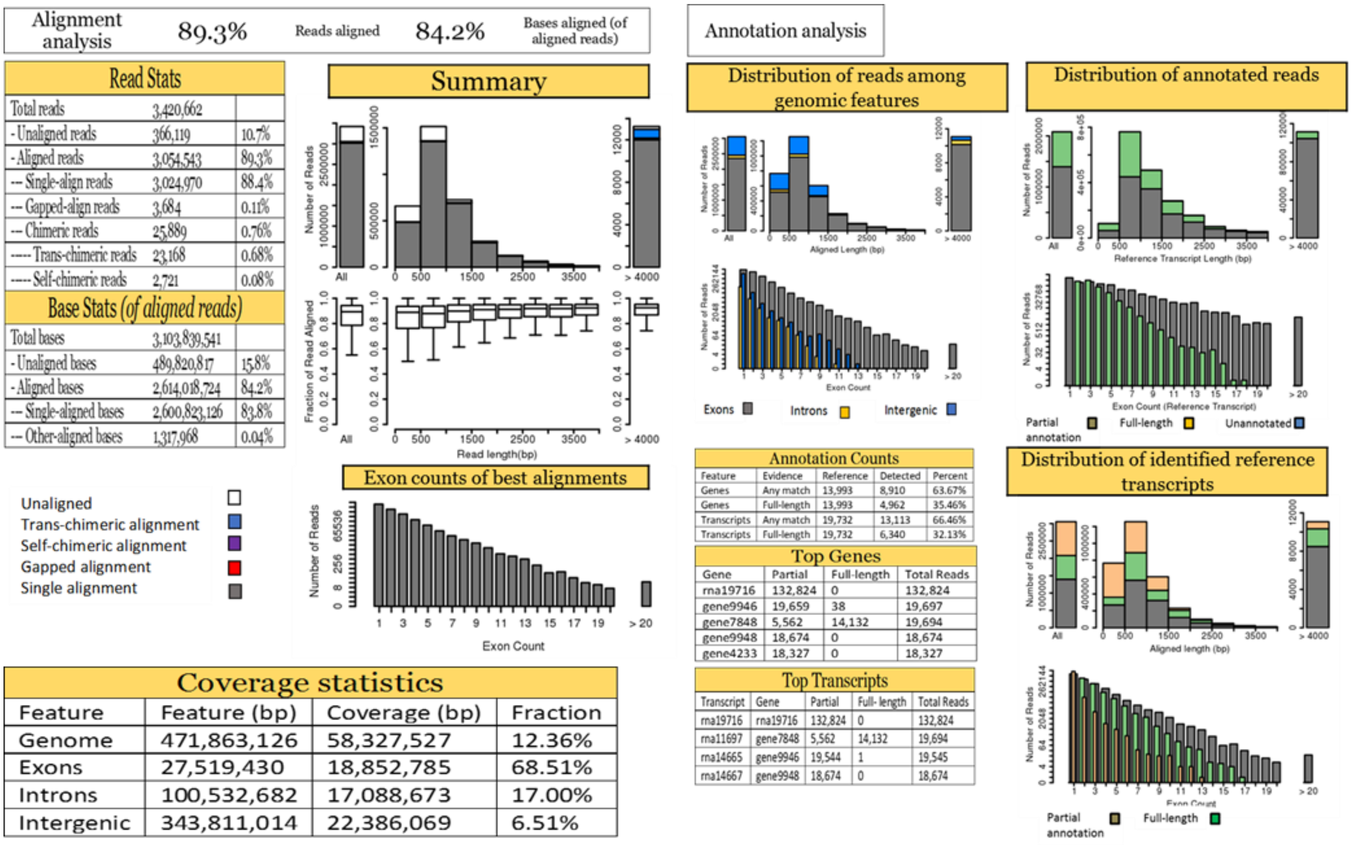


**Extended Figure 12 A. Alignment QC (Using the 6-hour timepoint as example)**

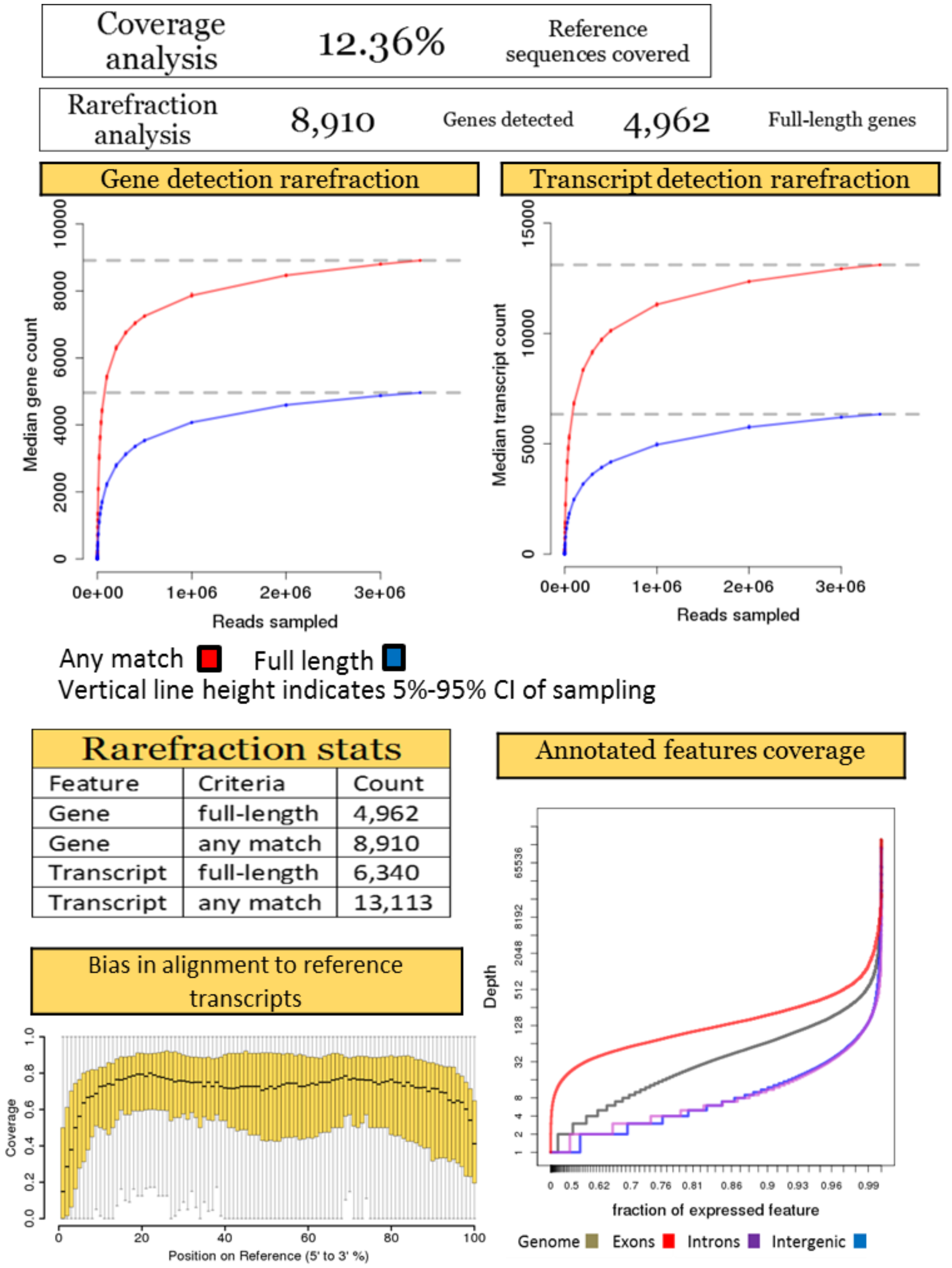


**Extended Figure 12 B**

Error pattern analysis 16.835% Error rate
based on aligned segments

**Error rates, given a target sequence**

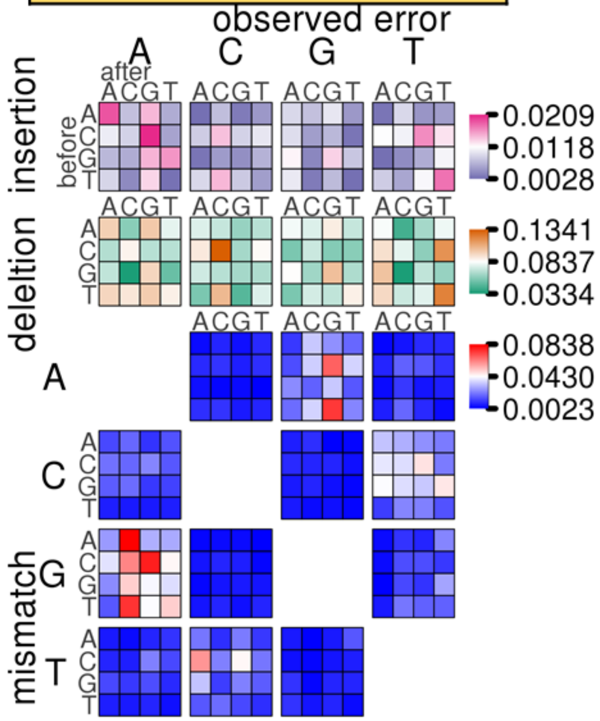


**Alignment-based error rates**

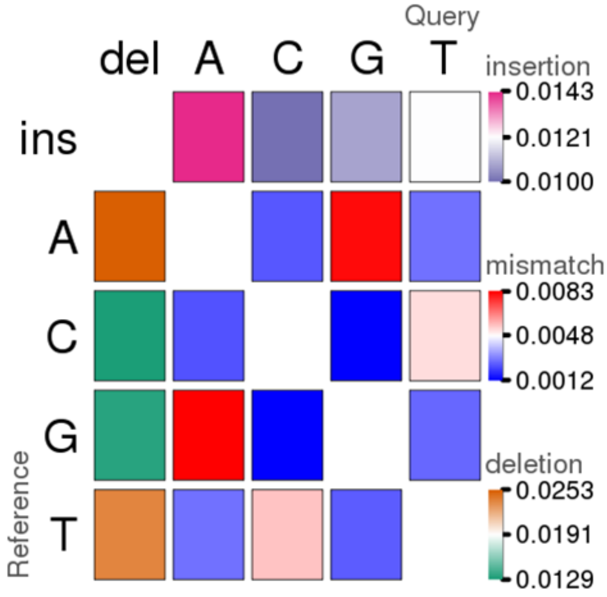


| Alignment stats | | |
|---|---|---|
| Best alignments sampled | 1101 | |
| **Base stats** | | |
| Bases analyzed | 1,060,762 | |
| - Correctly aligned bases | 882,179 | 83.2% |
| - Total error bases | 178,583 | 16.835% |
| --- Mismatched bases | 48,409 | 4.564% |
| --- Deletion bases | 80,111 | 7.552% |
| ----- Complete deletion bases | 61,366 | 5.785% |
| ----- Homopolymer deletion bases | 18,745 | 1.767% |
| --- Insertion bases | 50,063 | 4.720% |
| ----- Complete insertion bases | 32,343 | 3.049% |
| ----- Homopolymer insertion bases | 17,720 | 1.670% |

**Distance of observed junctions from reference junctions**

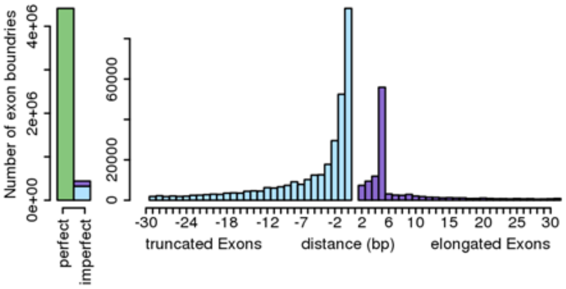


**Extended Figure 12 C**

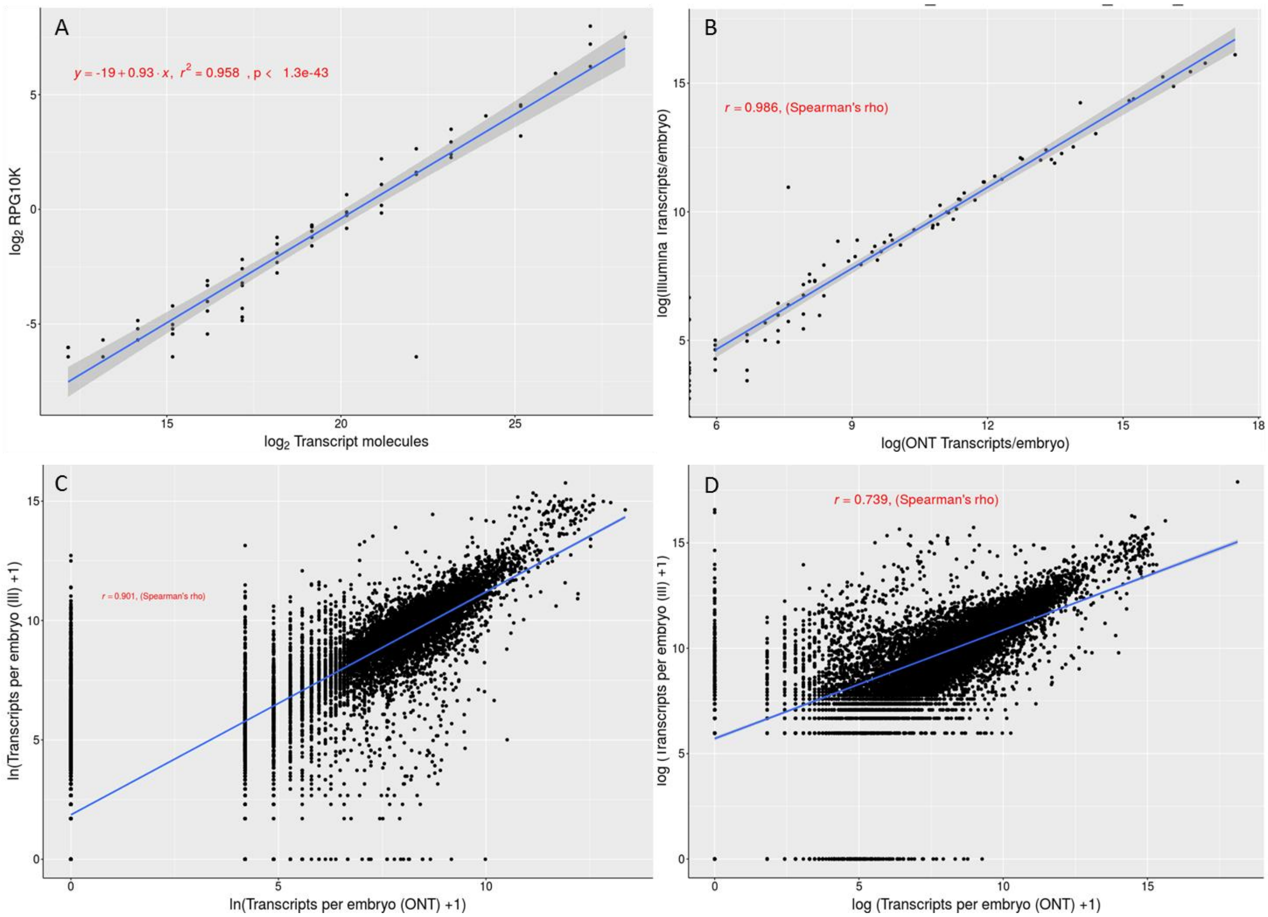


**Extended Figure 13.** E) Correlation of Illumina short-read and ONT long-read quantification of RNA standards (ERCC). F) Correlation of Illumina short-read and ONT long-read quantification of annotate genes.

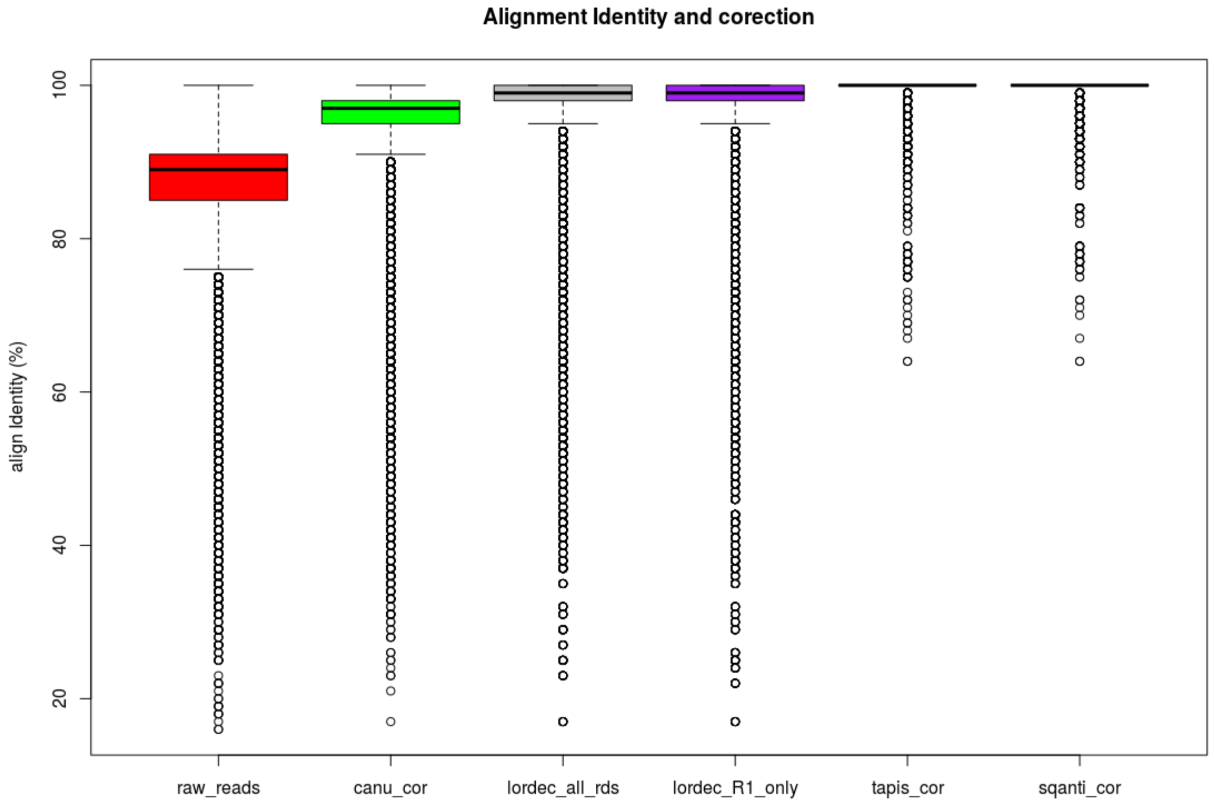


**Extended Figure 14.** Comparison of alignment identity following error correction. Raw_reads refers to the raw Nanopore cDNA reads. canu_cor refers to raw reads after one round of Canu correction. lordec_all_rds refers to Canu-corrected reads after one round of Lordec correction using read1 and read2 of Illumina short reads. Lordec_R1_only refers to Canu-corrected reads after one round of Lordec correction using only read1 of Illumina short reads. tapis_cor and sqanti_cor refer to reads after correction with TAPIS and SQANTI. TAPIS and SQANTI performed 3 rounds of genome guided error correction.

Extended Figure 16. Temporal clustering of gene expression using DGGP

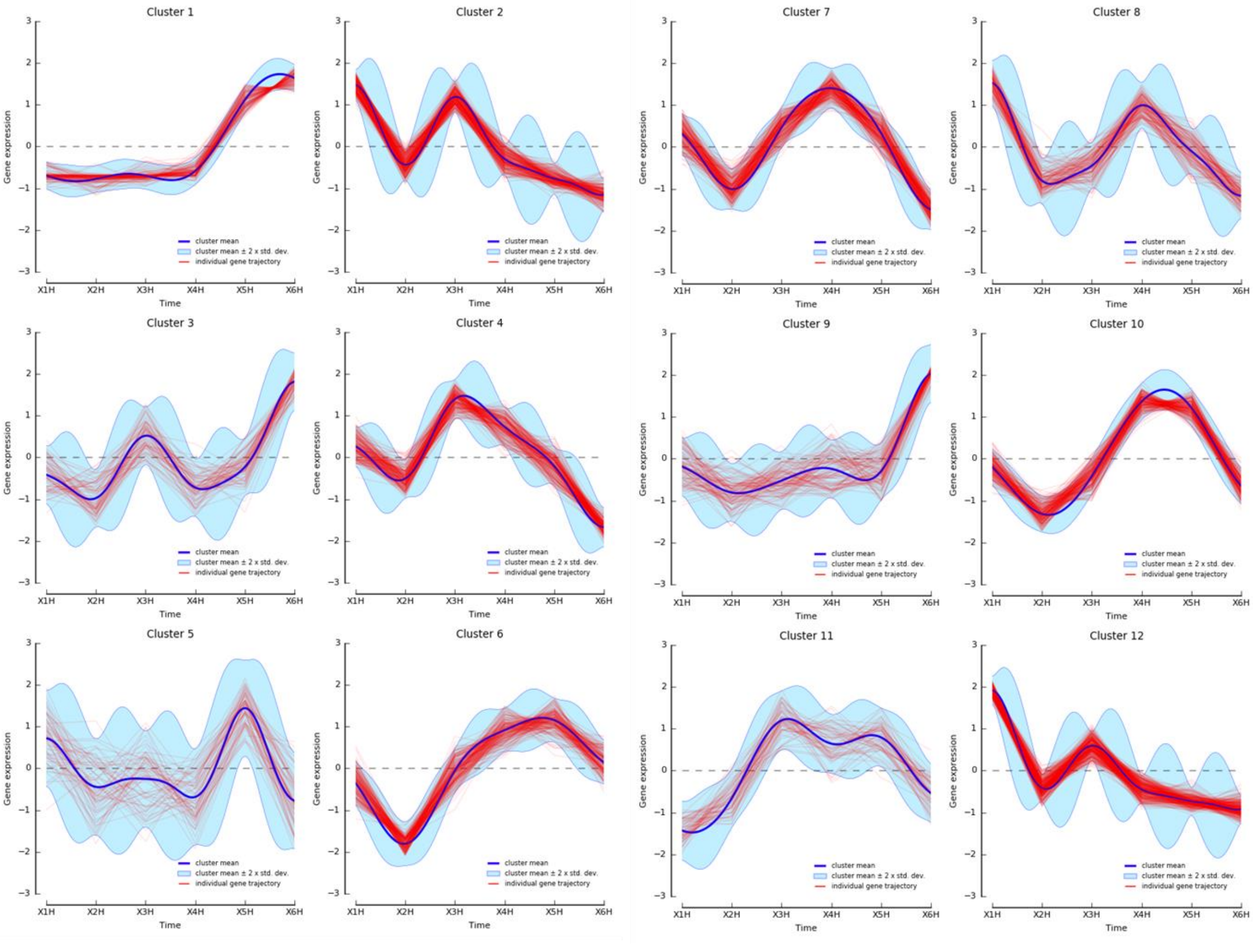


**Extended Figure 15 A. Clustered trajectories of expressed genes across the early embryonic development of *B. oleae.***

(picture continuous in the next page)

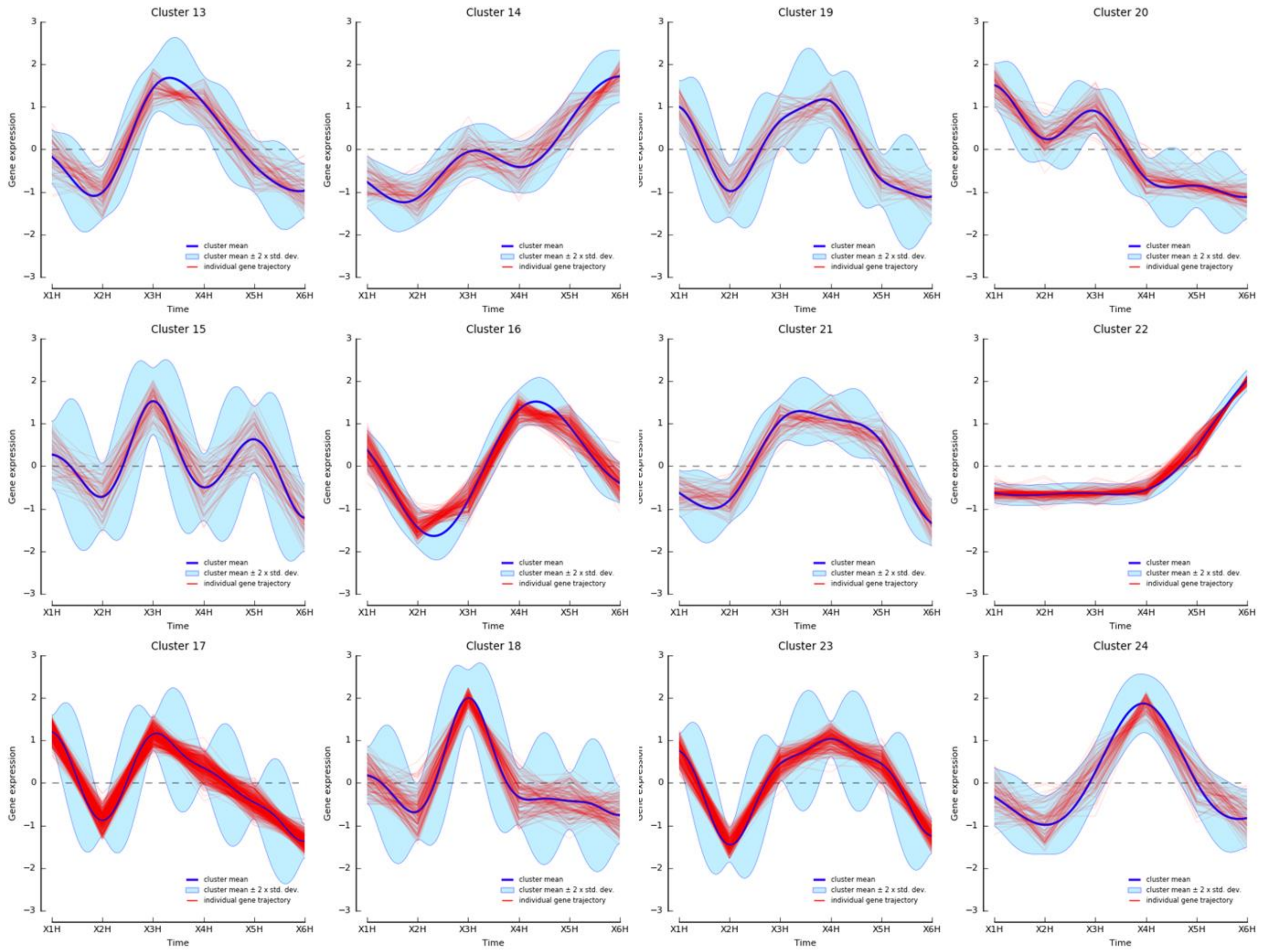


**Extended Figure 16 B. Clustered trajectories of expressed genes across the early embryonic development of *B. oleae*.**

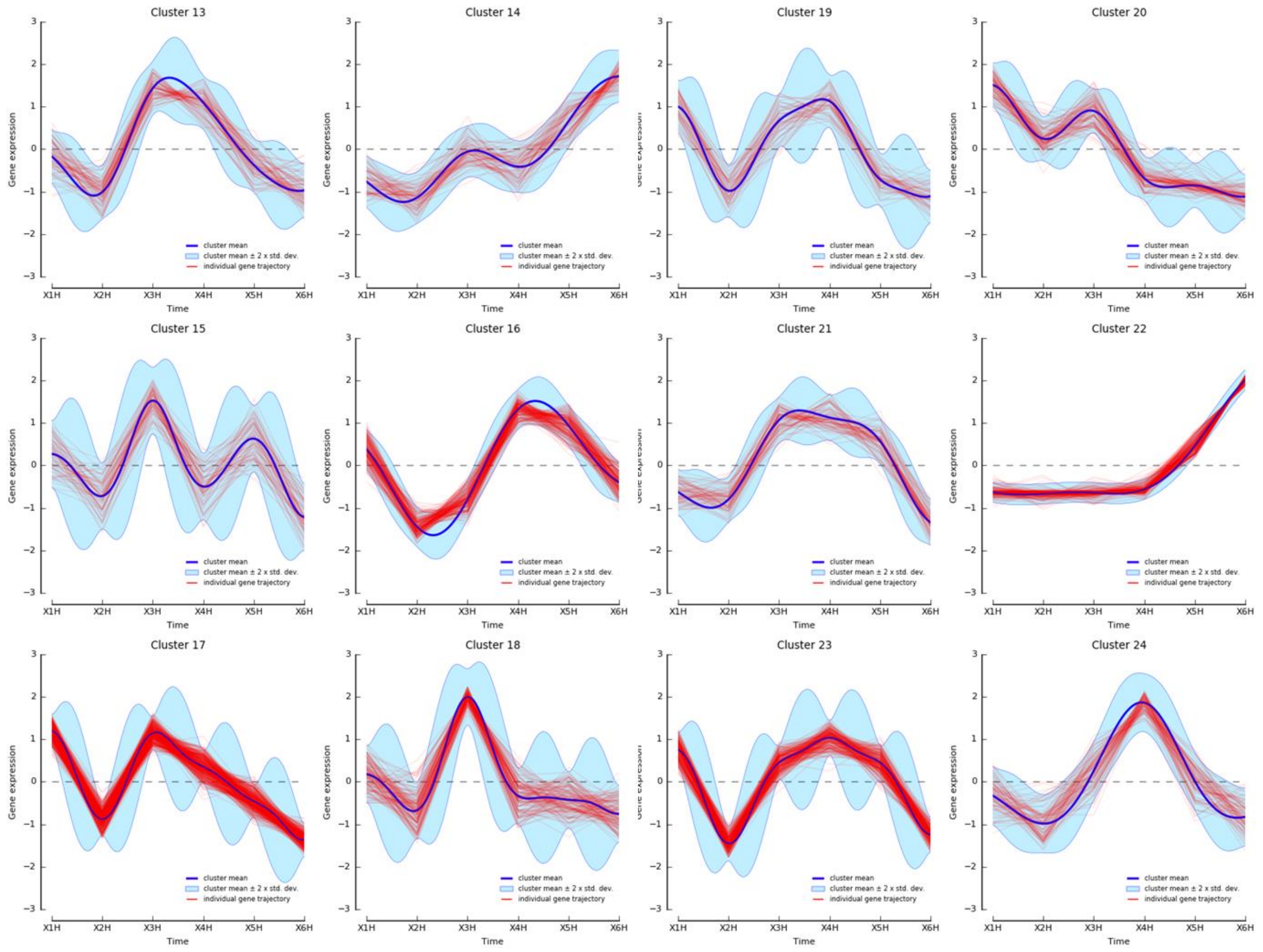


**Extended Figure 16 C. Clustered trajectories of expressed genes across the early embryonic development of *B. oleae*.**

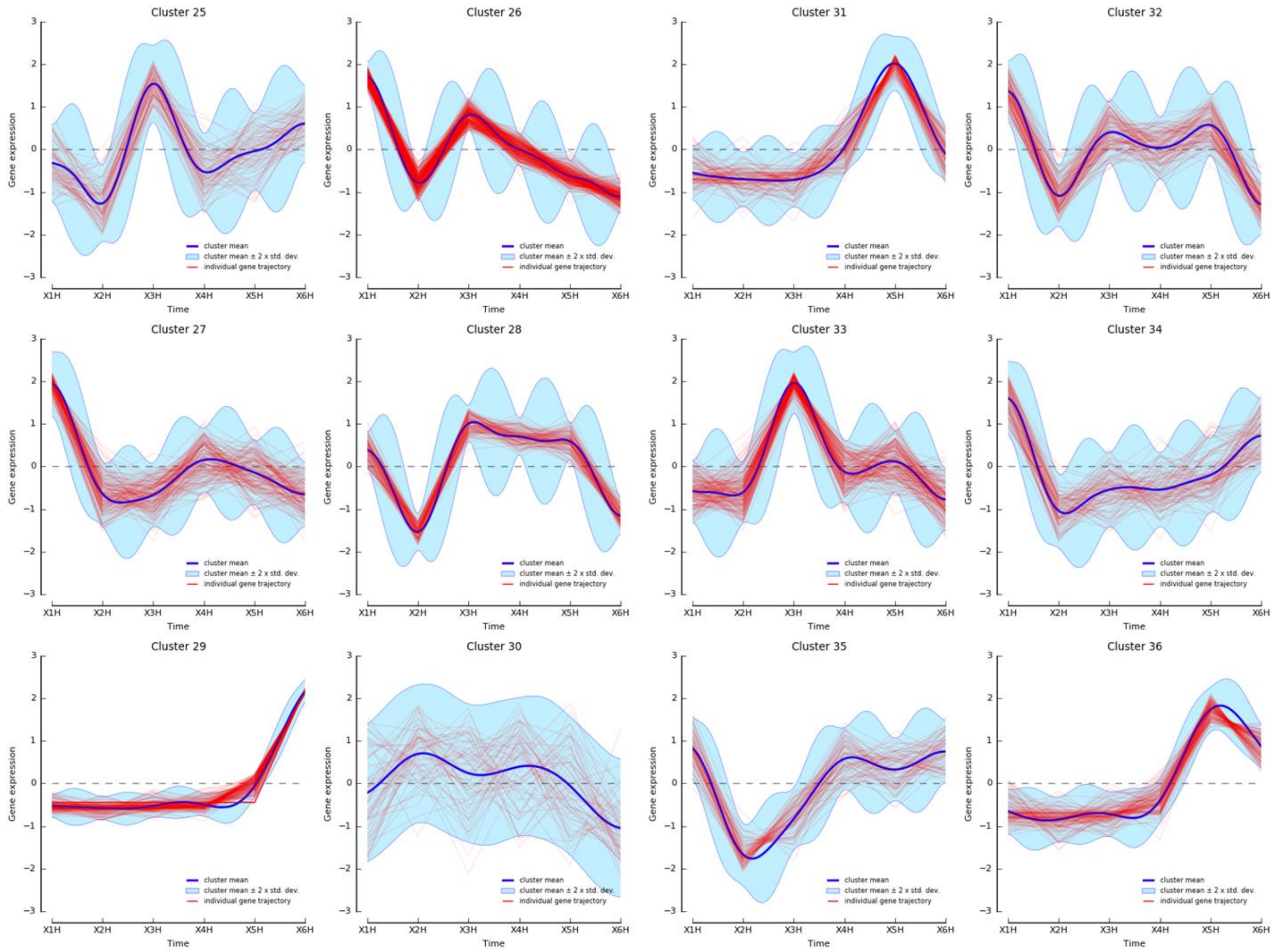


**Extended Figure 16 D. Clustered trajectories of expressed genes across the early embryonic development of *B. oleae*.**

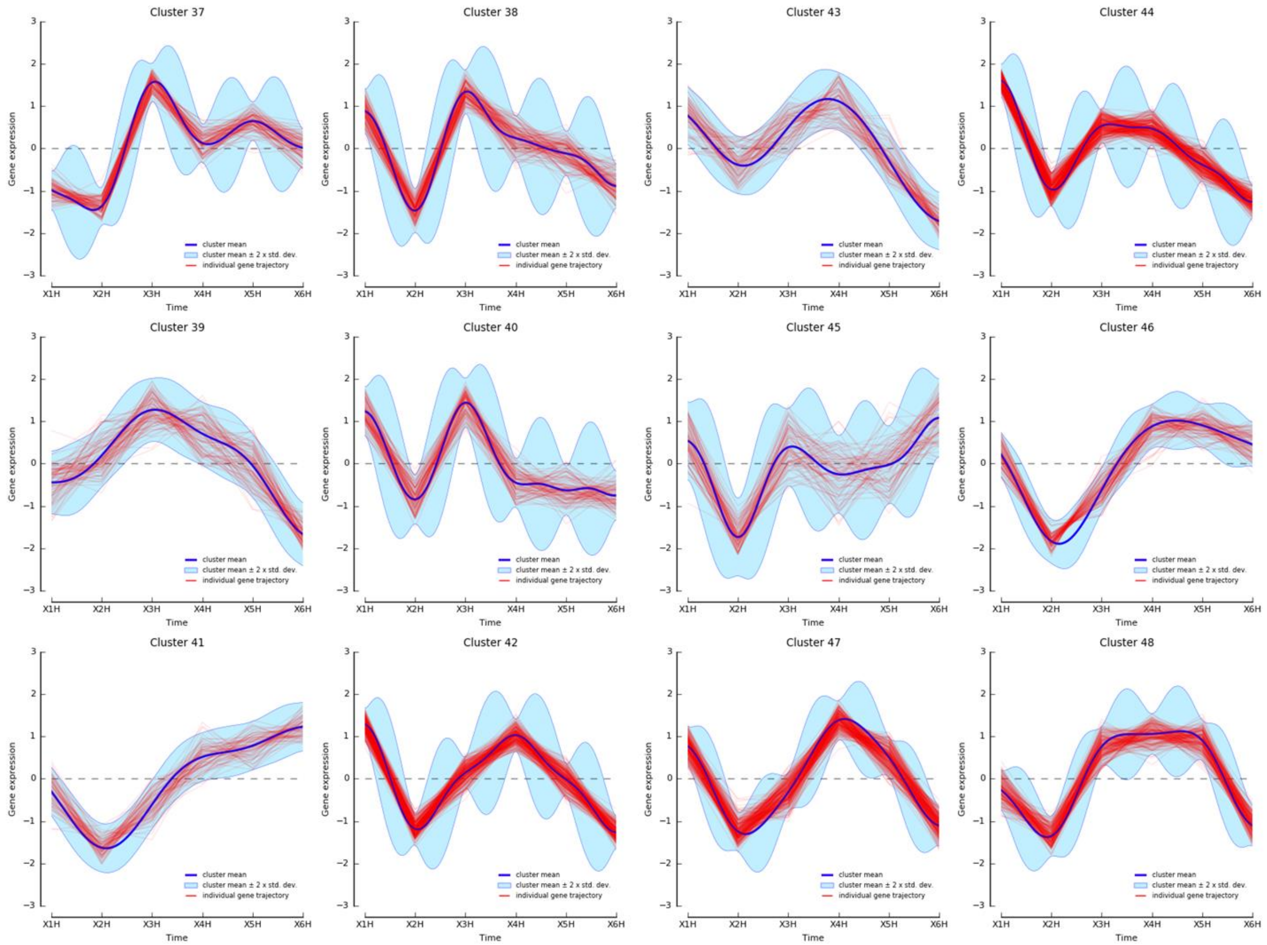


**Extended Figure 16 E. Clustered trajectories of expressed genes across the early embryonic development of *B. oleae*.**

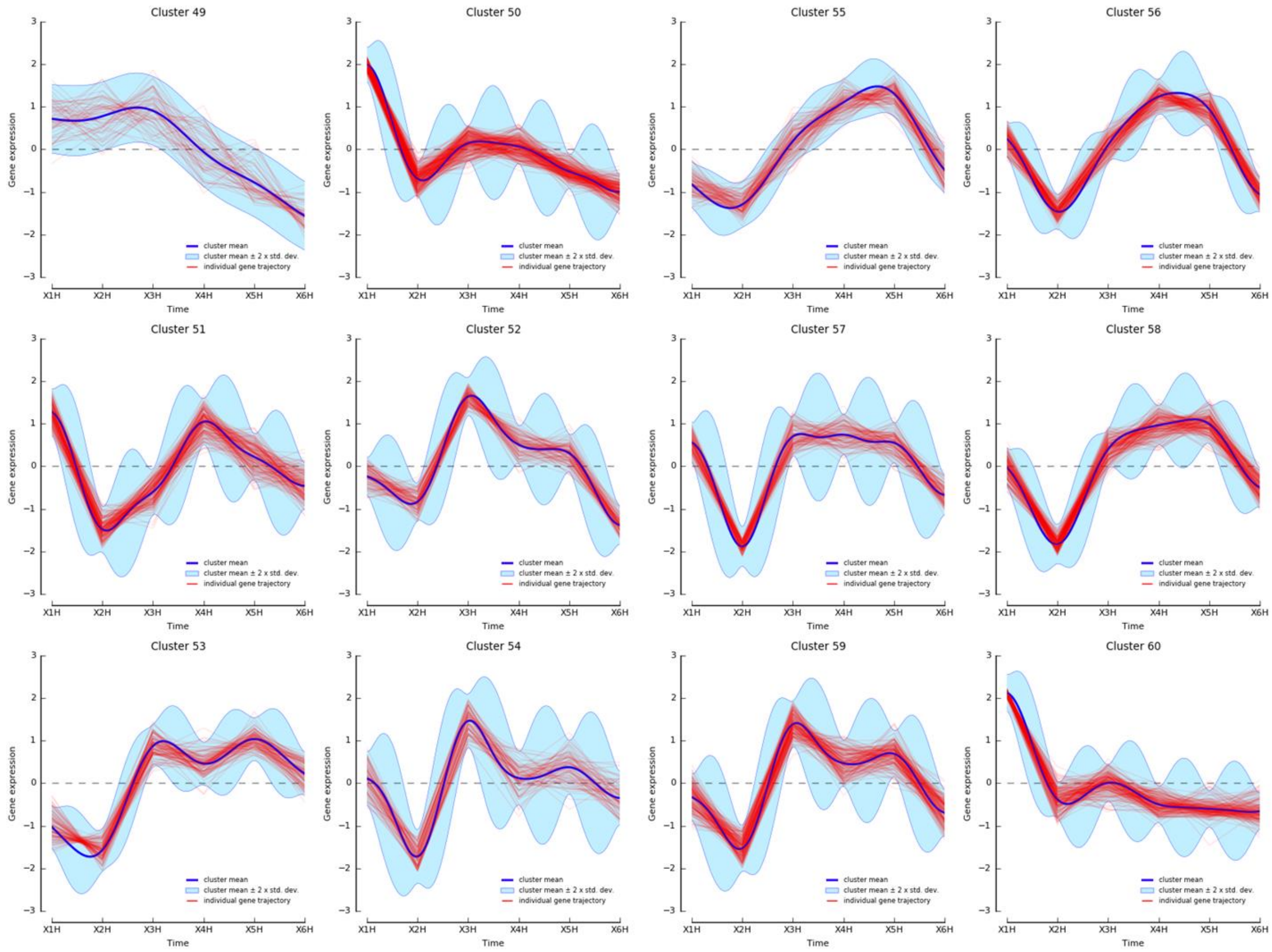


**Extended Figure 16 F. Clustered trajectories of expressed genes across the early embryonic development of *B. oleae*.**

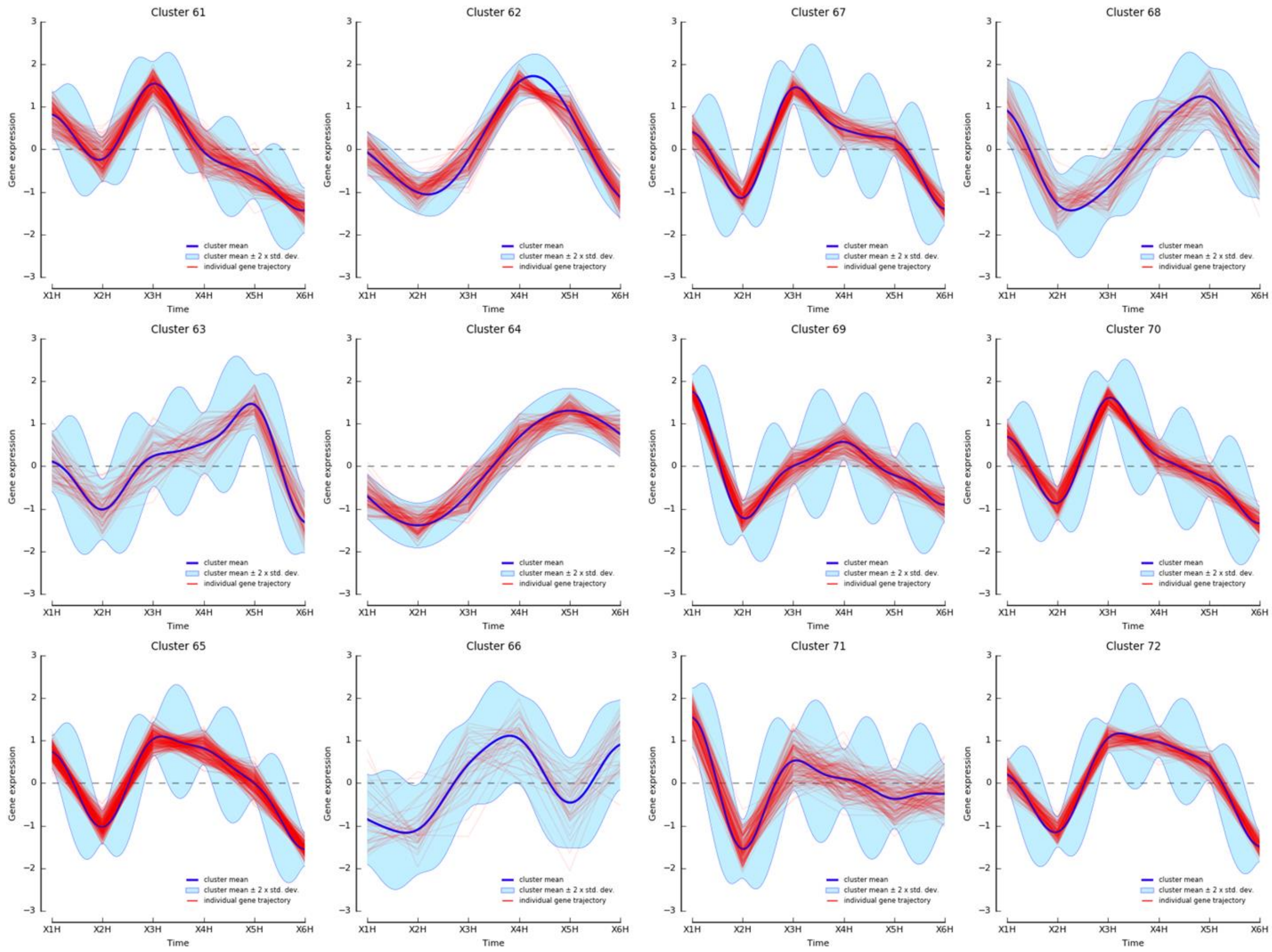


**Extended Figure 16 G. Clustered trajectories of expressed genes across the early embryonic development of *B. oleae*.**

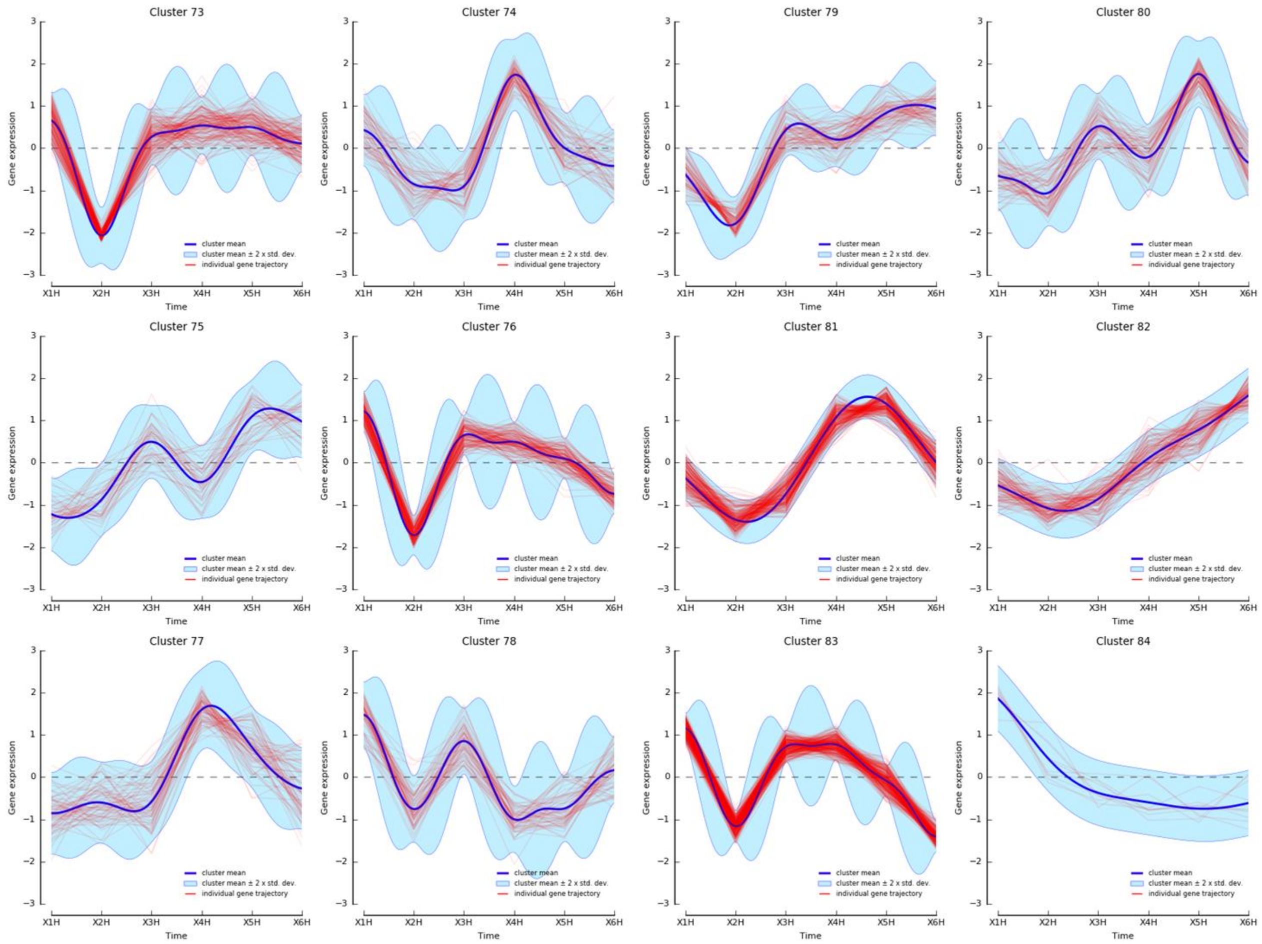


**Extended Figure 16 H. Clustered trajectories of expressed genes across the early embryonic development of *B. oleae*.**

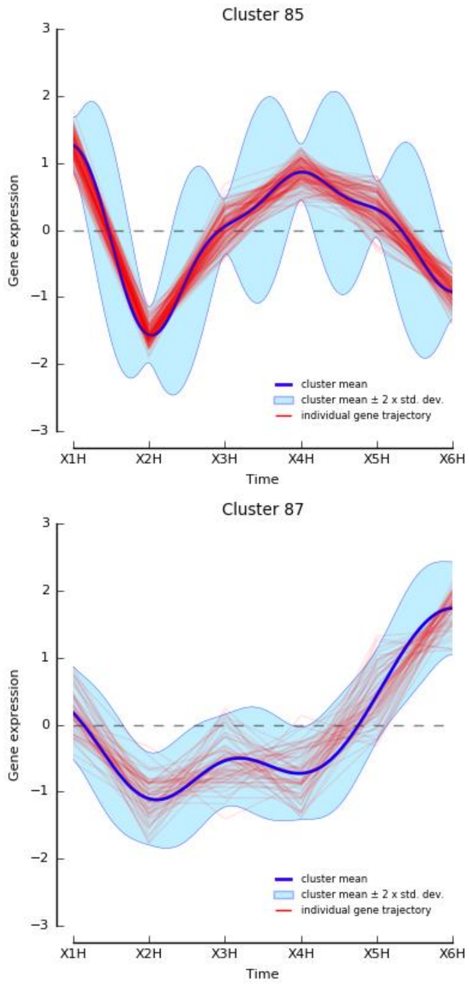


**Extended Figure 16 I. Clustered trajectories of expressed genes across the early embryonic development of *B. oleae*.**
