## Supplementary material for "Transcriptome landscape of the developing olive fruit fly embryo delineated by Oxford Nanopore long-read RNA-Seq"

### Protocol for cDNA synthesis and cDNA sequencing on the Oxford Nanopore Technologies (ONT) MinION platform

#### Materials and Reagents needed

#### General reagents

- RNase Zap (Thermo Fischer Scientific, AM9780)
- 1M Tris-HCl pH 8.0 (Thermo Fischer Scientific, AM9855G)
- Magnetic Stand for 1.5 mL tubes (e.g. Ambion P/N AM10026) and 0.2 mL tubes
- Filtered tips (10, 20, 100, 200, 1000 μL), and respective pipettes
- Eppendorf Centrifuge 5424R or 5424 (or equivalent)
- Micro-Centrifuge for 0.2 mL PCR tubes
- Qubit Assay Tubes (Thermo Fischer Scientific; Q32856)
- Qubit Fluorometer (Life Techonologies)
- VWR PCR 8-Tube Strip 0.2 mL (120 Strips) (VWR, 53509-304)
- Agilent TapeStation 2200 and the corresponding assay tubes
- Eppendorf DNA LoBind Tubes 1.5 mL (022431021)
- Mixer e.g HulaMixer (Thermo Fischer Scientific), Vortex Mixer (VWR)
- BioRad Thermo Cycler T100
- Agencourt AMPure XP beads (A63880, Beckman Coulter, 5 mL)
- Ethanol 100 % (reagent grade)

### 1 Reagents to assess the quality of the extracted RNA

#### Reagents for RNA Quantification

- Qubit RNA HS Assay Kit (Thermo Fischer Scientific, Q32852)

#### Reagents to examine the RNA profile

- Agilent RNA ScreenTape Ladder (Agilent; 5067-5578)
- Agilent RNA ScreenTape Sample Buffer (Agilent; 5067-5577)
- Agilent RNA ScreenTape (Agilent; 5067-5576)

### 2 Reagents for cDNA synthesis

#### Oligos and reconstitution buffer for cDNA synthesis primers:

- PolyT primer (V = A or C or G, N = A or C or G or T) (RNase-Free HPLC purification of the synthesized oligo is preferable. The oligo should be shipped lyophilized)

  5'-AAGCAGTGGTATCAACGCAGAGTATGCAACGCAACT$_{(30)}$VN -3'

- TSO oligo (TSO: Template-Switching oligonucleotide, the red marked bases are ribonucleotides. RNase-Free HPLC purification of the synthesized oligo is preferable. The oligo should be shipped lyophilized)
  5'- AAGCAGTGGTATCAACGCAGAGTGGATTCTATCACGCrGrGrG -3'

- THE RNA Storage Solution (Thermo Fischer Scientific, AM7000)

#### Reagents to remove DNA contamination in total RNA samples

- DNA-free DNA Removal Kit (Thermo Fischer Scientific, M1906)

**Enzymes and reagents for the cDNA synthesis reaction:**

- RNAse inhibitor 40 U/uL (Clontech, 2313A)
- Advantage UltraPure PCR Deoxynucleotide Mix (10 mM each dNTP) (Clontech, 639125)
- Water nuclease free PCR grade (eg. Affymetrix, 901578)
- SuperScript IV (Thermo Fischer Scientific, 18090010)
- Betaine (5M) (Sigma-Aldrich, B0300-1VL)
- $MgCl_2$ (1M) (Thermo Fisher Scientific, AM9530G)

**Reagents to spike RNA molecules of known abundance inside the sample RNA:**

- ERCC RNA Spike-In Mix (Thermo Fisher Scientific, 4456740)

**3 Reagents for cDNA amplification:**

**Primer for cDNA amplification:**

- cDNA amplification primer (Standard Desalting of the synthesized oligo can be ordered. The oligo can be shipped lyophilized or reconstituted at standard 100µM concentration.)

5'- /5Phos/ TCGTCGGCAGCGTCAAGCAGTGGTATCAACGCAGAGT-3'

**Enzymes for cDNA amplification:**

- Advantage 2 PCR Kit (Clontech, 639207)

**4 Reagents for cDNA Quality Control:**

**Reagents for cDNA quantification**

- Qubit HS DNA Assay Kit (Thermo, Q32851)

**Reagents to examine the cDNA profile**

- Agilent D5000 ScreenTape (Agilent, 5067-5588)
- Agilent D5000 Reagents (Agilent, 5067-5589)
- Agilent D5000 Ladder (Agilent, 5067-5590)

### 5 Reagents for cDNA library preparation for the nanopore platform

**End repair of the cDNA molecules**

- NEBNext End Repair Module (New England Biolabs, E6050S)

**d(A) tailing of the cDNA molecules**

- NEBNext dA-Tailing Module (New England Biolabs, E6053S)

**Ligate ONT adapters on the cDNA molecules**

- Ligation 1D Sequencing kit SQK-LSK108
- NEB Blunt/TA Master Mix (New England Biolabs, M0367S)
- Flow Cell Wash Kit (EXP-WSH002)

### 6 ONT MinION Sequencing

- MinION SpotON FLO-MIN106 flow cells
- MinION Mk1b

**Preparation of reagents**

1. The TSO oligo is reconstituted in "THE RNA Storage Solution" at a concentration of 1200 uM. *The information sheet from the manufacturer usually provides a dilution volume for a solution with a 100 uM oligo concentration. To create the solution with the 1200 uM oligo concentration, adjust the dilution volume accordingly by reducing 12X times the recommended volume presented on the information sheet*. Then 1 ul is diluted in 99ul of RNA storage solution (100X dilution; final concentration: 12 uM) and stored in aliquots at 5.6 ul per tube. The aliquots are stored at -80$^{O}$ C. The TSO ribonucleotides are prone to degradation. Loss of the ribonucleotides will lead in considerable reduction/absence of cDNA yield.
2. The PolyT primer is reconstituted in nuclease free $H_20$ at a concentration of 1200uM. *The information sheet from the manufacturer usually provides a dilution volume for a solution with a 100 uM oligo concentration. To create the solution with the 1200 uM oligo concentration, adjust the dilution volume accordingly by reducing 12X times the recommended volume presented on the information sheet*. Then 1 ul is diluted in 99ul of nuclease free $H_20$ and stored in aliquots at 7 ul per tube. The aliquots can be stored at -80$^{O}$ C.

**RNA quantification**

Total RNA was quantified using the “Qubit RNA HS Assay Kit” according to manufacturer instructions.

**Assess DNA contamination in the RNA extraction**

DNA contamination was measured using the Qubit dsDNA HS Reagent.

**Removal of DNA contamination from total RNA**

We used the DNA-free DNA Removal according to manufacturer instructions for the removal of DNA from RNA samples.

**Assess the profile of the extracted RNA**

Total RNA profile was determined using the Agilent RNA Screentape following manufacturer instructions except that the samples are not heated at 72 °C.

**Spike-In RNA**

ERCC (ERCC RNA Spike-In Mix 1) were added during the cDNA synthesis step. We aimed to obtain a final percentage of 5 % of our reads assigned to ERCCs assuming that ploy(A) fraction of total RNA is 5 %. We target the sequenced reads of the spiked-in RNA to be 5% of the total amount of sequenced reads). The amount of spiked RNA ($mass_{spiked\ RNA}$) that is going to be added in the reaction mix can be calculated as follows:

$$\mathbf{mass_{spiked\ RNA}} = \frac{\mathbf{fraction_{spiked\ reads}} \times \mathbf{fraction_{target\ RNA}} \times \mathbf{mass_{RNAinput}}}{\mathbf{Total\_RNA\_extracted}}$$

where:

**mass** $_{\textbf{spiked RNA}}$**:** mass (ngs) of spike-in RNA (SIRVs or ERCC) to be added in the sample.

**fraction** $_{\textbf{spiked reads}}$**:** desired fraction of sequenced spike-in RNA reads relative to the total amount of sequenced reads.

**fraction** $_{\textbf{target RNA}}$**:** fraction of the total RNA used in the sample, that is going to be synthesized into cDNA molecules.

**mass** $_{\textbf{RNA input}}$**:** mass (ngs) of RNA input per sample.

Then the volume (ul) of spike-in RNA to be used is calculated as follows:

$$\mathbf{volume_{spike-in\ RNA}} = \frac{\mathbf{mass_{spike-in\ RNA}}}{\mathbf{concentration_{spike-in\ RNA}}}$$

where

$\mathbf{concentration_{spike-in\ RNA}}$: concentration (ngs/ul) of the spike-in RNAs solution.

**volume$_{\text{spike-in RNA}}$:** volume (ul) from the spike-in RNAs solution to be added into the sample.

The value for the "mass $_{\text{RNA input}}$" is mass $_{\text{RNA input}}$ =300 ngs (We will use in the cDNA synthesis reactions 300 ngs of total RNA).

- For the ERCC RNA Spike-In Mix 1, the **mass $_{\text{spiked RNA}}$** = 0.45 ngs

1. The concentration of the stock solutions are:

- The "ERCC RNA Spike-In Mix 1" tube contains 10 µl of ERCC RNAs at a concentration of 103.515 fmoles/ul or 30.3 ng/ul .

- Prepare the appropriate dilution of each Spike-In Mix needed. In the new diluted solution the "mass $_{\text{spiked RNA}}$" for either the "ERCC RNA Spike-In Mix 1" or the "Spike-in RNA Variant (SIRVs) Control set 3 kit" should correspond, *if possible*, to 0.1 ul of the final diluted volume. So we need to have the following dilutions:
  - For the "ERCC RNA Spike-In Mix 1" we are going to dilute 6.72 times the stock solution. So in 5.72 ul of "THE RNA solution" add 1 ul from the "ERCC RNA Spike-In Mix 1" stock solution (new concentration= 4.5 ng/ul). Afterwards we will have to take Volume $_{\text{spike-in RNA}}$ = ((0.45 ngs)/(4.5 ng/ul))=0.1 ul of the diluted solution.

**cDNA Library generation and sequencing on MinION**

Generally, we followed the ONT "1D Strand switching cDNA by ligation (SQK-LSK108)" protocol but with custom cDNA synthesis protocol (as described below), and the end repair and d(A) tailing steps were performed separately. An overview of the protocol is presented in **Error! Reference source not found.** and it is the following:

1. cDNA synthesis and amplification
2. End-repair of cDNA molecules
3. dA-tail of cDNA molecules
4. Adapter ligation
5. Sequencing
6. Base-calling

**cDNA synthesis**

Our cDNA synthesis protocol involved a customized version of the Smart-seq protocol[1]. The protocol is based on the terminal deoxynucleotidyl transferase activity of the wild-type MMLV (Moloney murine leukemia virus) reverse transcriptase[2].

**Preparation of Master Mixes**

1. Thaw and vortex all reagents and keep master mixes on ice until use.
2. Label three 1.5 ml eppendorf tubes: "**pre-RT**", "**RT**", "**PCR**"
3. Always use fresh TSO primer as it is prone to degradation.
4. Prepare the "**pre-RT mix**" according to Table 1 below.

Table 1: **pre-RT mix**

| | **pre-RT mix** | Total RNA (ul /sample) |
|---|---|---|
| 1 | ERCC RNA Spike-In Mix 1 | x |
| 2 | RNase Inhibitor (40 U/uL * 125 uL = 5000U) | 0.05 |
| 3 | Poly-T primer (stock: 12 uM) | 0.7 |
| 4 | Superscript IV first-strand buffer (5×) | 0.4 |
| 5 | Nuclease free water | 0.19 |

| | | |
|---|---|---|
| 6 | dNTP Mix (stock: 10 mM each) | 0.56 |
| | **Total =** | **2** |

5. Pipette 2uL of **pre-RT mix** to a PCR tube and add 1uL of sample (**300ng of total RNA)**. Include a negative control (**1uL of water/RNA buffer**).
6. Incubate the samples in a thermocycler set according to Table 2 below.

**Table 2: pre-RT incubation**

| **Temperature** | **Time** | **Purpose** |
|---|---|---|
| 72°C | 3 min | Unfolding of RNA secondary structures, Poly-T primer binding |
| 4°C | 10 min | Poly-T primer binds |
| 25°C | 1 min | Poly-T primer binds more specifically |
| 4°C | Hold | |

7 . Prepare the “**RT mix** according to Table 3

Table 3: **RT mix**

| | **RT mix** | ul /sample |
|---|---|---|
| 1 | Nuclease free $H_20$ | 0.85 |
| 2 | Superscript IV first-strand buffer (5×) | 0.8 |
| 3 | DTT (stock: 100 mM) | 0.175 |
| 4 | TSO (stock: 12 µM) | 0.7 |
| 5 | RNAse inhibitor (stock: 40 U/ µl) | 0.175 |
| 6 | SuperScript IV reverse transcriptase (stock: 200 U/ ul) | 0.35 |
| 7 | Betaine (stock: 5 M) | 0.7 |
| 8 | $MgCl_2$ (stock: 100 mM) | 0.25 |
| | **Total =** | **4** |

8. Following pre_RT incubation, add 4 ul of RT mix to each sample, mix and briefly spin down.
9. Incubate the samples in a thermocycler set according to Table 4 below

Table 4: **SSIV RT protocol**

| **Temperature** | **Time** | **Cycle** | **Purpose** |
|---|---|---|---|
| 50°C | 10 min | 1 | RT and template-switching |
| 55°C | 30 sec | 10 | Unfolding of RNA secondary structures |
| 50°C | 30 sec | | Completion/continuation of RT |

| | | | |
|---|---|---|---|
| 60°C | 30 sec | 5 | Unfolding of RNA secondary structures |
| 55°C | 30 sec | | Completion/continuation of RT |
| 50°C | 30 sec | 1 | Finish template switching |
| 65°C | 30 sec | 5 | Unfolding of RNA secondary structures |
| 60°C | 30 sec | | Completion/continuation of RT |
| 50°C | 30 sec | 1 | Finish template switching |
| 70°C | 30 sec | 5 | Unfolding of RNA secondary structures |
| 65°C | 30 sec | | Completion/continuation of RT |
| 50°C | 30 sec | 1 | Finish template switching |
| 75°C | 30 sec | 5 | Unfolding of RNA secondary structures |
| 70°C | 30 sec | | Completion/continuation of RT |
| 50°C | 1 min | 1 | Final finish template switching |
| 80°C | 10 min | 1 | Enzyme inactivation |
| 4°C | Hold | 1 | |

10. Prepare the **PCR master mix** according to Table 5

### Table 5: PCR master mix

| | **PCR Mix** | (ul per 7 ul of RT reaction) |
|---|---|---|
| 1 | PCR-Grade Water | 47.6 |
| 2 | 10X Advantage 2 PCR Buffer (not SA, short amplicon) (Advantage 2 PCR Kit) | 7 |
| 3 | 50X dNTP Mix (Advantage 2 PCR Kit) | 2.8 |
| 4 | PCR primer (stock: 12 µM) | 2.8 |
| 5 | 50X Advantage 2 Polymerase Mix (Advantage 2 PCR Kit) | 2.8 |
| | **Total =** | **63** |

11. Following RT incubation, add 63 ul of PCR mix to each sample, mix and briefly spin down
12. Incubate the samples in a thermocycler set according to Table 6 below

**Table 6: PCR protocol**

| **Temperature** | **Time** | **Cycle** |
|---|---|---|
| 95°C | 1 min | 1 |
| 95°C | 20 sec | 5 |
| 58°C | 4 min | |
| 68°C | 6 min | |
| 95°C | 20 sec | 11 or 12 cycles , as many to produce around 1-2 ug of cDNA per 70 ul of PCR amplification reaction |
| 64°C | 30 sec | |
| 68°C | 6 min | |
| 72°C | 10 min | 1 |
| 4°C | Hold | 1 |

**13.** The amplified product is subsequently cleaned with Agencourt AMPure XP beads as is described below.

**Agencourt AMPure XP cleanup of cDNA amplification products**

1. Allow AMPure XP beads to equilibrate to room temperature for at least 30 minutes.
2. Vortex the beads until evenly mixed, then add 0.9X sample volume of Agencourt AMPure XP beads to the sample in the same tube as used for PCR.
3. Pipet the entire volume up and down to mix thoroughly. Place the sample tubes on a roler mix for 5 - 8 minutes to let the DNA bind to the beads. Briefly spin the samples to collect the liquid from the side of the tube.
4. Place the sample tubes on the magnetic separation device for ~2 minutes until the liquid appears completely clear, and there are no beads left in the supernatant.
5. While the samples are on the magnetic separation device, pipette out the supernatants. Keep the samples on the magnetic separation device. Add 200 µl of freshly made 80% ethanol to each sample without disturbing the beads. Wait for 30 seconds and carefully pipette out the supernatant containing contaminants.
6. DNA will remain bound to the beads during the washing process. Repeat step 4 once more. Briefly spin the samples to collect the liquid from the side of the wall.
7. Place the samples on the magnetic device for 30 seconds, then remove all the remaining ethanol with a pipette.
8. Place the samples at room temperature until the pellet appears dry (~ 5 minutes). You may see a tiny crack in the pellet when it is dry.
9. Once the beads are dry, add 51 µl of TE buffer to cover the bead pellet.
10. Remove the samples from the magnetic separation device and mix thoroughly to resuspend the beads. Incubate the sample with rotation at room temperature for 5 – 8 minutes.
11. Put the tubes on the magnet and after ~2 minutes recover the supernatant which should contain the cleaned amplified cDNA. Determine the quantity of the cDNA and profile using Qubit HS DNA Assay Kit and Agilent D5000 Tapestation, respectively, following manufacturer instructions.

**End -repair of DNA**

End repair of 1 µg of amplified cDNA was carried out using NEBNext End Repair Module (New England Biolabs, E6050S) following manufacturer instructions. This was followed by 0.9X Ampure XP beads cleanup (described above).

### dA-tailing reaction

d(A) tailing of the recovered end-repaired cDNA was carried out using NEBNext dA-Tailing Module (New England Biolabs, E6053S) following manufacturer instructions. This was followed by 0.9X Ampure XP beads cleanup (described above).

### Adapter ligation

Ligation of ONT sequencing adapters onto recovered d(A)-tailed cDNA (up to 1 µg) was carried out following ONT SQK-LSK-108 protocol. However, we increased the incubation time from 10 minutes to 1 - 4 hours at room temperature.

### ONT MinION sequencing kit

ONT SQK-LSK-108 protocol was followed for the sequencing part.

### Basecalling

We performed our basecalling off-line using Albacore version 2.0.2

### Data analysis

Basecalling
Albacore (ONT, version 2.0.2)
read_fast5_basecaller.py -r --flowcell SQK-LSK108 --kit SQK-LSK108 --input %s --save_path %s --worker_threads 23 -o fastq" %(input_dir,save_path))

Minionqc[3] (version 1.0)
Rscript ~/MinionQC.R -p 23 -i $('pwd')/files -o $('pwd')/results

Pauvre (version 0.1.2, https://github.com/conchoecia/pauvre)
pauvre marginplot --no-transparent --fastq ../Bo_E_1H_C010_10_pass.fastq > pauvre.out 2> pauvre.out

Porechop (version 0.2.3, https://github.com/rrwick/Porechop)
~/porechop --format fasta -t 47 -i $read5.fasta -o $read5.choped.fasta > porechop.stdout 2> porechop.stdout

Cutadapt[4] poly(A) trimming from read ends (version 1.15)
~/.local/bin/cutadapt --info-file=trim_info -f fasta -a "A[100]" -o $read2.cutadapt.fasta $read2.fasta

GMAP[5] (version 2018-03-25)
GMAP for alignment QC
~/gmap -t 23 -D $dirc -f samse -d $ref $read1 > $outsam.sam

GMAP for transcriptome assembly
~/gmap -t 23 -D $dirc --cross-species --max-intronlength-ends=10000 -n 1 -z sense_force -f samse -d $ref $read1 > $outsam1.sam 2> gmap.stdout

Minimap2[6] (version 2.9 (r720))
~/minimap2 -ax splice -t 23 $ref $reads1 > $outsam1.sam

Samtools[7] (version 1.3.2)

AlignQC[8] (version 1.2)
~/alignqc analyze $outsam1.sort.bam --specific_tempdir $dirc/tmp1 -r $ref -a $annotation -o alignqc.xhtml --output_folder $dirc/alignQC.ouput_b4_correction > alignqc.stdout

Canu[9] (Canu 1.7)
canu useGrid=false -correct gnuplotImageFormat=png corOutCoverage=10000 corMhapSensitivity=high corMinCoverage=0 correctedErrorRate=0.16 overlapper=minimap ovsMethod=sequential minReadLength=200 minOverlapLength=100 genomeSize=1500000000 -p Bo_E_all_pass_edited -d Bo_E_all_pass_edited -nanopore-raw Bo_E_all_pass_edited.fasta

LoRDEC[10] (v0.8, using GATB v1.4.1)

~/lordec-correct -2 $illumina_reads -T 47 -p -k 19 -s 3 -i $nanopore.fasta -o "$nanopore"_lordec_corrected.fasta

GFOLD[11] (v1.1.4)
gfold diff -norm NO -s1 Bo.E.2H -s2 Bo.E.1H -suf .abs_cnt3 -o Bo.E.2HvsBo.E.1H.abs.diff > Bo.E.2HvsBo.E.1H.abs.diff.stdout

cDNA_Cupcake (version 5.3, https://github.com/Magdoll/cDNA_Cupcake/wiki)
~/collapse_isoforms_by_sam.py --input $read1 -s $outsam.sorted.sam --dun-merge-5-shorter -o $pref
~/filter_by_count.py $pref.collapsed --min_count=2 >filter_by_count.stdout
~/filter_away_subset.py $pref.collapsed >filter_away_subset.stdout
~/filter_away_subset.py $pref.collapsed.min_fl_2
cDNA_Cupcake for assembly evaluation using 5-hour timepoint
~/collapse_isoforms_by_sam.py -c 0.95 -i 0.95 --input $read1 -s $sortedsam --dun-merge-5-shorter -o $pref

TAMA (version tc0.0, https://github.com/GenomeRIK/tama)
~/tama_collapse.py -d merge_dup -s $sortedsam -f $ref -p $pref -x no_cap -c 95 -i 95

TAPIS[12] (1.2.1)
alignPacBio.py -p 22 -v -K 10000 -o tapis_output $indexesDir $indexName $reference $reads
run_tapis.py -p -t 30 -o run_tapis_output $annotation tapis_output/$bamfile

SQANTI[13] (version 1.2)
sqanti_qc.py -z -t 47 -fl $fl_abundance -c $sj_covIllumina -e $isoExpression -x $gmapindex -o $output -d qc_output $isoforms.fa $gtf $ref
sqanti_filter.py -d filter_output -i "$isoforms"_corrected.fasta "$output"_classification.txt

### References


1 Ramskold, D. *et al.* Full-length mRNA-Seq from single-cell levels of RNA and individual circulating tumor cells. *Nat Biotechnol* **30**, 777-782, doi:10.1038/nbt.2282 (2012).

2 Zajac, P., Islam, S., Hochgerner, H., Lonnerberg, P. & Linnarsson, S. Base preferences in non-templated nucleotide incorporation by MMLV-derived reverse transcriptases. *PLoS One* **8**, e85270, doi:10.1371/journal.pone.0085270 (2013).

3 Lanfear, R., Schalamun, M., Kainer, D., Wang, W. & Schwessinger, B. MinIONQC: fast and simple quality control for MinION sequencing data. *Bioinformatics (Oxford, England)*, doi:10.1093/bioinformatics/bty654 (2018).

4 Martin, M. Cutadapt removes adapter sequences from high-throughput sequencing reads. *EMBnet.journal; Vol 17, No 1: Next Generation Sequencing Data AnalysisDO - 10.14806/ej.17.1.200* (2011).

5 Wu, T. D. & Watanabe, C. K. GMAP: a genomic mapping and alignment program for mRNA and EST sequences. *Bioinformatics (Oxford, England)* **21**, 1859-1875, doi:10.1093/bioinformatics/bti310 (2005).

6 Li, H. Minimap2: pairwise alignment for nucleotide sequences. *Bioinformatics (Oxford, England)*, doi:10.1093/bioinformatics/bty191 (2018).

7 Li, H. *et al.* The Sequence Alignment/Map format and SAMtools. *Bioinformatics (Oxford, England)* **25**, 2078-2079, doi:10.1093/bioinformatics/btp352 (2009).

8 Weirather, J. L. *et al.* Comprehensive comparison of Pacific Biosciences and Oxford Nanopore Technologies and their applications to transcriptome analysis. *F1000Research* **6**, 100, doi:10.12688/f1000research.10571.2 (2017).

9 Koren, S. *et al.* Canu: scalable and accurate long-read assembly via adaptive k-mer weighting and repeat separation. *Genome research* **27**, 722-736, doi:10.1101/gr.215087.116 (2017).

10 Salmela, L. & Rivals, E. LoRDEC: accurate and efficient long read error correction. *Bioinformatics (Oxford, England)* **30**, 3506-3514, doi:10.1093/bioinformatics/btu538 (2014).

11 Feng, J. *et al.* GFOLD: a generalized fold change for ranking differentially expressed genes from RNA-seq data. *Bioinformatics (Oxford, England)* **28**, 2782-2788, doi:10.1093/bioinformatics/bts515 (2012).

12 Abdel-Ghany, S. E. *et al.* A survey of the sorghum transcriptome using single-molecule long reads. *Nature communications* **7**, 11706, doi:10.1038/ncomms11706 (2016).

13 Tardaguila, M. *et al.* SQANTI: extensive characterization of long-read transcript sequences for quality control in full-length transcriptome identification and quantification. *Genome research*, doi:10.1101/gr.222976.117 (2018).
